# A mechanistic model of primer synthesis from catalytic structures of DNA polymerase α–primase

**DOI:** 10.1101/2023.03.16.533013

**Authors:** Elwood A. Mullins, Lauren E. Salay, Clarissa L. Durie, Noah P. Bradley, Jane E. Jackman, Melanie D. Ohi, Walter J. Chazin, Brandt F. Eichman

## Abstract

The mechanism by which polymerase α**–**primase (polα–primase) synthesizes chimeric RNA-DNA primers of defined length and composition, necessary for replication fidelity and genome stability, is unknown. Here, we report cryo-EM structures of polα–primase in complex with primed templates representing various stages of DNA synthesis. Our data show how interaction of the primase regulatory subunit with the primer 5′-end facilitates handoff of the primer to polα and increases polα processivity, thereby regulating both RNA and DNA composition. The structures detail how flexibility within the heterotetramer enables synthesis across two active sites and provide evidence that termination of DNA synthesis is facilitated by reduction of polα and primase affinities for the varied conformations along the chimeric primer/template duplex. Together, these findings elucidate a critical catalytic step in replication initiation and provide a comprehensive model for primer synthesis by polα–primase.

## Introduction

DNA polymerase α**–**primase (polα–primase) is unique among the replicative polymerases by virtue of its ability to perform *de novo* synthesis from a DNA template. Polα– primase initiates DNA synthesis by generating short (∼30-nucleotide) primers ^1–3^, which are required for leading and lagging strand replication by polε and polδ ^4^. Generation of primers is particularly important on the lagging strand, where discontinuous DNA synthesis is repeatedly initiated during genome replication and telomere elongation ^5^. As such, polα–primase is essential for replication of the genome, and defects in priming have been implicated in cellular ageing and an impaired DNA damage response ^6–10^.

Polα–primase possesses both DNA-dependent RNA polymerase (primase) and DNA polymerase (polα) activities. Primase initiates synthesis from nucleoside triphosphates (NTPs) to generate a 7–10-ribonucleotide primer ^11,12^, which is extended by ∼20 deoxyribonucleotides by polα (**Fig. 1A**) ^1,2^. The resulting chimeric RNA-DNA primers are extended by polδ to generate each Okazaki fragment. Polα lacks proofreading capability and is therefore mutagenic ^13^. The potentially deleterious effects of polα generated errors and incorporation of large amounts of RNA into the nuclear genome by primase are avoided by enzymatic removal of primers during Okazaki fragment maturation ^14–19^. Thus, the composition and short length of the primer synthesized by polα–primase is an important factor in genomic integrity.

**Figure 1.**
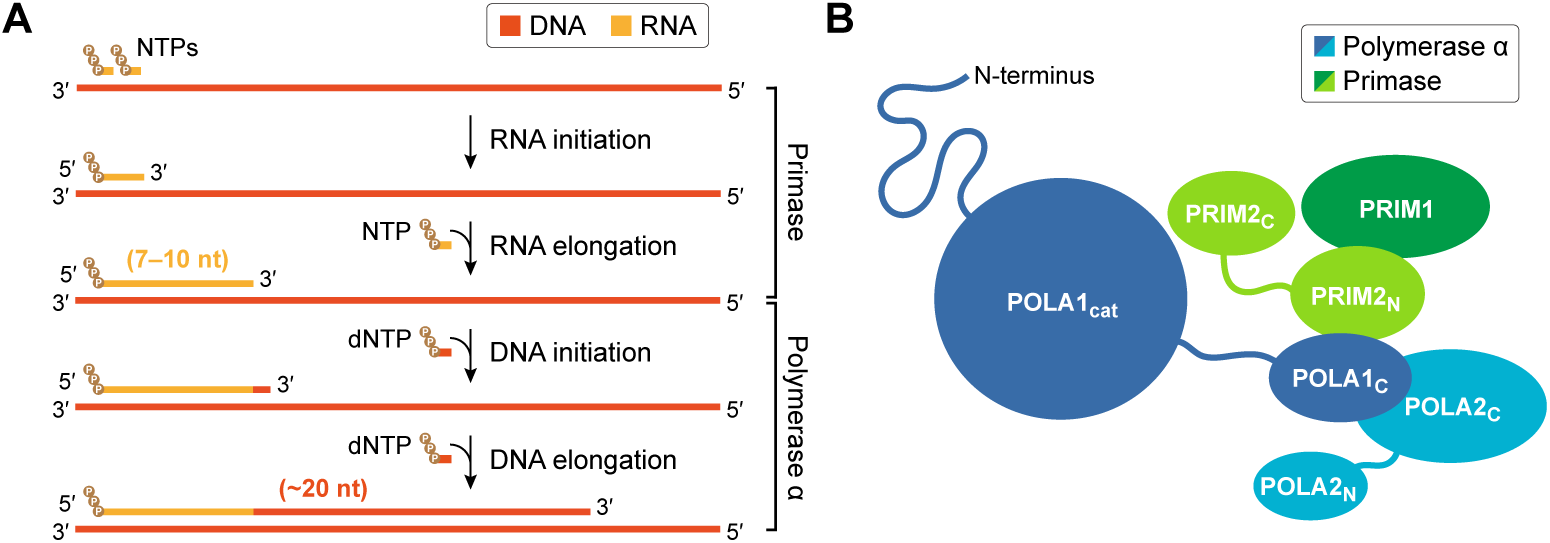
Chimeric primer synthesis. **A**. Stages of primer synthesis by polα–primase. Triphosphate moieties are depicted as brown circles. Primase and polymerase α synthesize RNA and DNA, respectively, to generate a short chimeric primer. **B**. Schematic of subunit and domain organization in the heterotetrameric polα–primase complex.

Polα–primase is a heterotetramer composed of two polymerase subunits—catalytic POLA1 (also referred to as p180) and regulatory POLA2 (p70, p68)—and two primase subunits—catalytic PRIM1 (p49, p48) and regulatory PRIM2 (p58) (**Fig. 1B**) ^20^. The primase dimer, which is necessary for normal RNA synthesis ^21^, is formed by interactions between PRIM1 and the N-terminal domain of PRIM2 (PRIM2_N_), with the C-terminal domain of PRIM2 (PRIM2_C_) tethered by a flexible linker ^22–29^. POLA1, which is functional on its own, relies on POLA2 for heterotetramer formation ^30–33^. Structural analysis of the yeast enzyme revealed a bilobal structure in which the catalytic domain of POLA1 (POLA1_cat_) is flexibly tethered via its C-terminal domain (POLA1_C_) to the tetramer core (TC), consisting of PRIM1, PRIM2_N_, POLA1_C_, and the C-terminal domain of POLA2 (POLA2_C_) ^26,30^. In the absence of nucleic acid substrates or protein partners, the human enzyme has only been observed in a compact, non-catalytic (“auto-inhibitory” or AI) configuration in which the active site of POLA1_cat_ is occluded by the tetramer core, as visualized by both X-ray crystallography and electron microscopy (EM) ^34,35^. The human enzyme has also been visualized in catalytically accessible configurations associated with the replisome ^36^ and the telomere maintenance CTC1–STN1–TEN1 (CST) complex ^37^. However, neither structure showed polα–primase engaged with primer/template.

Several models have emerged in recent years that largely attribute the counting ability of polα–primase to the PRIM2_C_ regulatory domain. Despite the catalytic center of primase residing in PRIM1, PRIM2_C_ is critical for all aspects of RNA synthesis, including regulation of RNA length ^23–25,38,39^. PRIM2_C_ has also been implicated in transfer of the RNA primer to polα by virtue of its interaction with the primer 5′-triphosphate and the template 3′-overhang ^21,34,40,41^. These interactions are believed to dictate RNA counting by spatially limiting the length of RNA synthesized by PRIM1, either via the PRIM2 interdomain linker ^41,42^ or via steric collision between PRIM2_C_ and PRIM2_N_ ^33,34^. While these models explain the upper limit of RNA length (i.e., 10 nucleotides), neither explains the range of RNA primers (7–10 nucleotides) extended by polα ^12^. PRIM2_C_ and the primer 5′-triphosphate have also been shown to be important for DNA synthesis ^43,44^, although the role of PRIM2_C_ in the transition from RNA to DNA synthesis and in DNA counting by polα is unknown. However, a recent model for DNA synthesis proposed a similar spatial limitation imposed by the POLA1 and PRIM2 interdomain linkers ^44^. Alternatively, structures of POLA1_cat_ bound to RNA/DNA and DNA/DNA duplexes afforded a model for DNA counting whereby preferential binding to relatively A-form duplexes by polα leads to decreased affinity and eventual dissociation from the primer/template as a result of a progressive increase in B-form character as DNA is synthesized ^32,45^.

The molecular details by which polα–primase synthesizes chimeric primers of defined length and composition have remained a matter of debate because there are few structures of the heterotetramer bound to primer/template ^46,47^, and no structures showing the intermediate and final stages of DNA synthesis. Here, we present cryo-EM structures of polα–primase alone and bound to primed templates representing DNA synthesis at early, intermediate, and final stages, beginning at the point immediately following transfer of the RNA primer to polα. Our structures reveal that dissociation of POLA1_cat_ and PRIM2_C_ from the AI complex enables a series of configurational and conformational changes that span a remarkable range of motion, both in the absence and the presence of nucleic acid substrates. At the onset of DNA synthesis, we observe a direct interaction between POLA1_cat_ and PRIM2_C_ that provides an explanation for how polα can extend RNA primers ranging in length from 7–10 nucleotides, and that suggests primer transfer is largely defined by competition between PRIM1 and POLA1_cat_ for the primer 3′- end. We also observe association of PRIM2_C_ with chimeric RNA-DNA substrates corresponding to intermediate and late stages of DNA synthesis. Complementary biochemical experiments comparing the synthesis activities of the full polα–primase complex and various subcomplexes are consistent with PRIM2_C_ acting as a limited DNA processivity factor, reducing the proportion of both shorter and longer primers ^44^. Additionally, we provide direct structural evidence linking primer composition, duplex conformation, and substrate affinity, with central roles for POLA1_cat_ and PRIM2_C_. Together, these findings establish a comprehensive model for chimeric primer synthesis by polα–primase.

## Results

### Polα–primase exists in an equilibrium between compact and extended configurations

Prior structures in the absence of nucleic acid substrates or protein partners have shown polα–primase in one of two states ^26,34,35,37,47–49^, suggesting the heterotetramer undergoes a dramatic structural transition from a compact, auto-inhibitory state to an extended, catalytically competent configuration prior to primer synthesis. To better understand the pre-existing equilibrium between the two states, we first analyzed the protein alone using single-particle cryo-EM. *Xenopus laevis* polα–primase was selected based on the stability and the quality of the purified complex (**Fig. 2A**) and the high sequence conservation with human polα–primase (69% identity and 90% similarity) (**Supplemental Figs. S1–S4**). We removed the N-terminal tail of POLA1 (residues 1–334) to further improve the stability of the complex. In the absence of nucleic acid substrates, 2D classification sorted 69% of particles into classes with density for all four subunits in the compact AI configuration, 30% into classes lacking density for POLA1_cat_, and the remaining 1% into a class with density for only POLA1_cat_ (**Fig. 2B and Supplemental Figs. S5 and S6**). We initially refined a partial 3D reconstruction of the AI complex to a resolution of 2.8 Å that exhibited poor density for PRIM1 (**Supplemental Figs. S5 and S7 and Supplemental Table S1**). To improve the interpretability of the map in this region, we identified a subset of particles that yielded a 3.2-Å reconstruction with clear density for all four subunits (**Fig. 2C, Supplemental Figs. S5 and S7, and Supplemental Table S1**). Only the small N-terminal domain of POLA2 (POLA2_N_) was not well defined. As previously observed in lower-resolution structures ^34,35,47,48^, the AI complex is stabilized by extensive contacts between POLA1_cat_ and every domain in the tetramer core, as well as PRIM2_C_. Conversely, PRIM2_C_ contacts only POLA1_cat_, PRIM2_N_, and the interdomain linkers in POLA1 (residues 1242–1275) and PRIM2 (residues 262–278). These interactions sterically occlude substrate recognition by both POLA1_cat_ and PRIM2_C_. To better understand the structural rearrangements necessary to bind nucleic acid substrates, dissociation of POLA1_cat_ and PRIM2_C_ from the AI complex was examined by 3D variability analysis, using all particles initially aligned on the AI complex during preliminary 3D refinement (**Supplemental Figs. S5 and S8**). Two principal motions were observed: (1) a hinging motion in PRIM2_N_ that pulls PRIM1 away from POLA1_cat_, and (2) a bending/twisting motion in POLA1_cat_ that pulls the exonuclease domain away from POLA2_C_ and that begins to pull the palm domain away from PRIM2_N_. Interactions involving the thumb domain were unchanged, as were all interactions involving PRIM2_C_.

**Figure 2.**
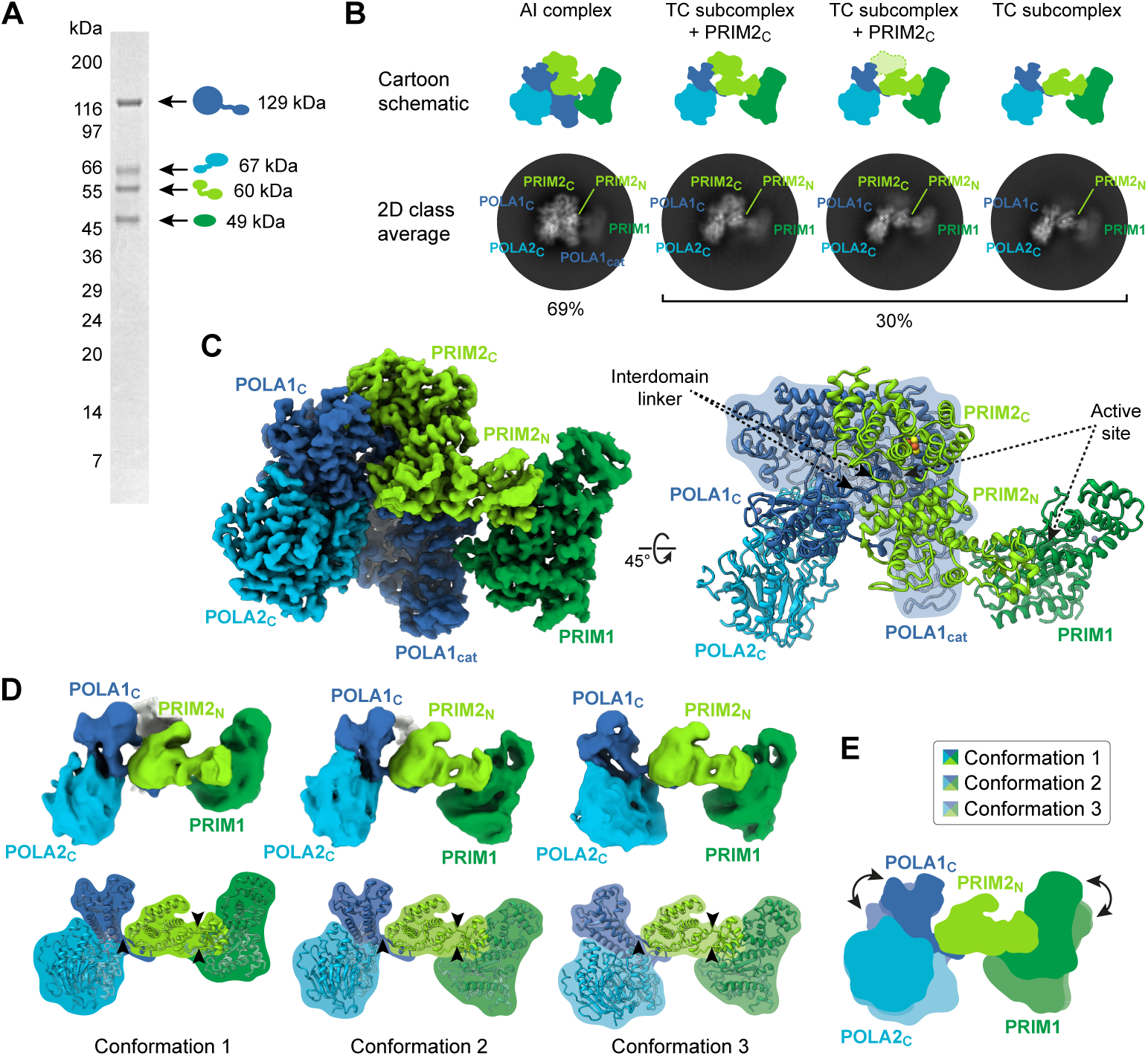
Conformational and configurational dynamics. **A**. Characterization of purified *Xenopus laevis* polα–primase by denaturing gel electrophoresis. Subunits are colored as in Fig. 1B (POLA1, blue; POLA2, cyan; PRIM1, dark green; PRIM2, light green). The POLA1 construct lacks residues 1–334. **B**. Representative 2D class averages and corresponding subunit schematics. The percentage of particles belonging to the AI complex or the TC subcomplex is indicated below the class averages. Class averages were selected to show PRIM2_C_ at various degrees of association with the tetramer core. All final 2D class averages are shown in **Supplemental Fig. S6**. **C**. 3D reconstruction of polα–primase in the auto-inhibitory configuration. *Left*, cryo-EM density map. *Right*, molecular model refined against the density map. POLA1_cat_ is outlined in blue. **D**. Distinct conformations of the tetramer core. *Top*, cryo-EM density maps. *Bottom*, molecular models docked into the density maps. Models were docked as three rigid bodies extracted from the AI complex. The points at which the complex were split and then reassembled are indicated with arrowheads. **E**. Schematic overlay aligned on the central rigid body of the three tetramer core conformations shown in panel D. The range of motion resulting from conformational dynamics is represented with curved arrows.

Without the conformational restrictions imposed by engagement of POLA1_cat_ in the AI complex, conformational heterogeneity is increased in the tetramer core. The subset of particles grouped into 2D classes that lacked density for POLA1_cat_ could be further divided into classes with clear, partial, or no density for PRIM2_C_ (**Fig. 2B and Supplemental Figs. S5 and S6**). 3D reconstructions of the tetramer core were generated from particles selected through multiclass ab initio reconstruction and heterogeneous refinement. These reconstructions revealed three distinct conformations characterized by hinging motions in which the PRIM1 and POLA1_C_/POLA2_C_ lobes move relative to one another by pivoting around flexible loops in PRIM2_N_ (residues 153–157 and 216–219) and POLA1_C_ (residues 1439–1443), respectively (**Fig. 2D,E and Supplemental Fig. S5**). Together, these data demonstrate that in the absence of nucleic acid substrates, polα–primase exists predominantly in a compact auto-inhibitory state, but also adopts a series of extended configurations in which POLA1_cat_ and PRIM2_C_ are disengaged from the other domains in the complex. Based on 3D variability analysis of the AI complex, along with inspection of the 2D class averages, we speculate that dissociation of POLA1_cat_ precedes dissociation of PRIM2_C_, and initiates the structural transition away from an auto-inhibitory state.

### PRIM2_C_ remains bound to the primer during DNA synthesis

Polα–primase possesses dual enzymatic activities that enable synthesis of chimeric RNA-DNA primers. To investigate the molecular bases for the transition from RNA to DNA synthesis and the low processivity of polα, we designed substrates to visualize polα–primase at two key points during DNA synthesis—immediately after intramolecular handoff of the RNA primer from primase to polα (DNA initiation, DI) and after polα has elongated the primer with DNA (DNA elongation, DE). The DI primer is comprised of nine ribonucleotides and the chimeric DE primer is comprised of nine ribonucleotides followed by 11 deoxyribonucleotides (**Fig. 3A, Supplemental Fig. S9, and Supplemental Table S2**). Both primers include the critically important 5′-triphosphate group that would be introduced by primase during RNA initiation. Ternary complexes containing polα–primase, primer/template, and incoming nucleotide were trapped using 2′,3′-dideoxycytidine to terminate the primer strand and prevent phosphodiester bond formation with dGTP.

**Figure 3.**
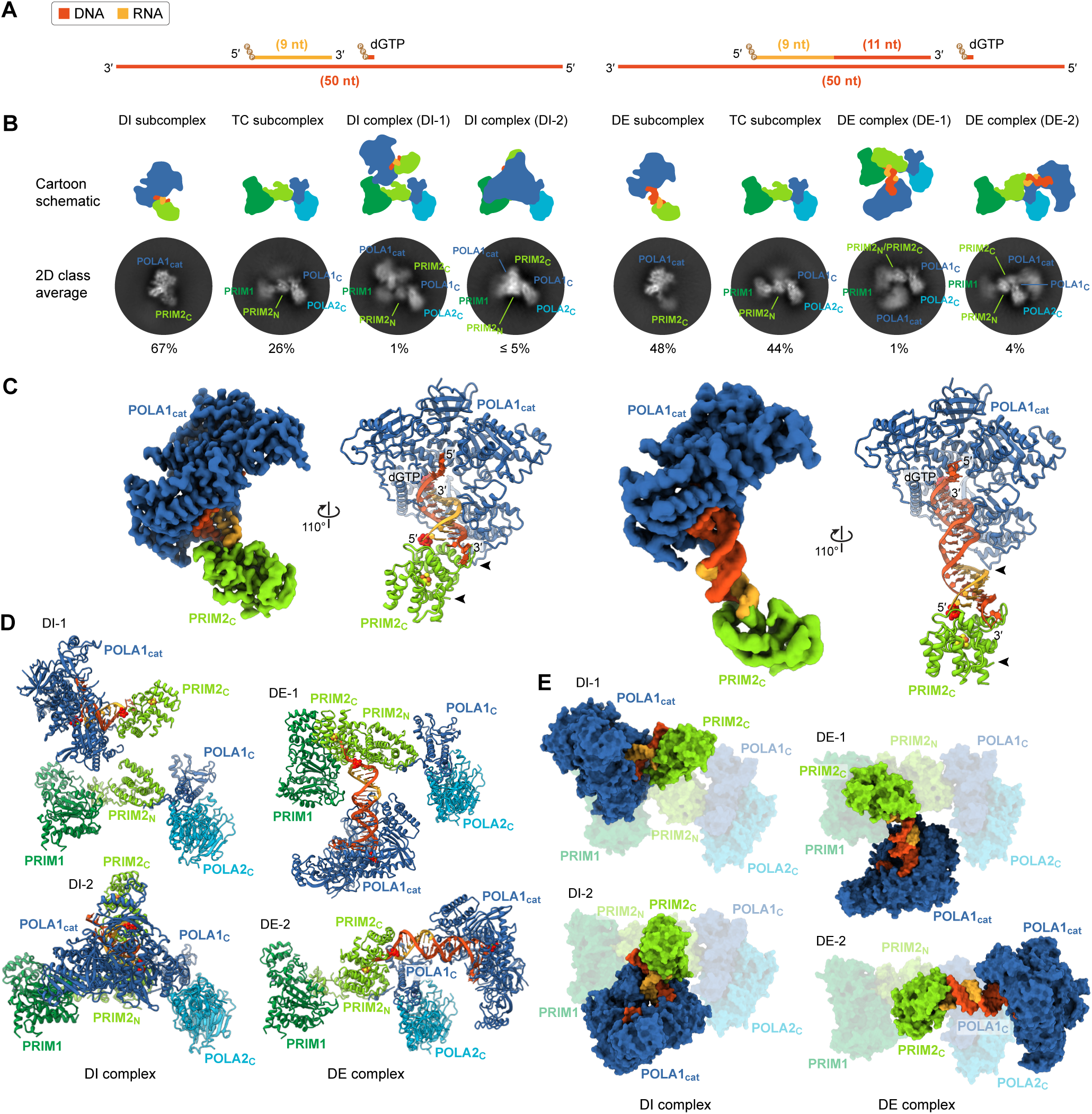
Configurational rearrangements during DNA initiation and DNA elongation. In each panel, DI and DE (sub)complexes are positioned on the left and the right, respectively. **A**. Substrate schematics. Triphosphate moieties are depicted as brown circles. In both substrates, the primers terminate in a 2′,3′-dideoxycytidine to prevent incorporation of dGTP. **B**. Representative 2D class averages and corresponding subunit schematics. The percentage of particles belonging to the DI or DE subcomplexes, the TC subcomplex, or one of the distinct configurations of the DI and DE complexes (DI-1, DI-2, DE-1, or DE-2) is indicated below the class averages. ∼3% of particles from the DI dataset could not be reliably assigned to a group. 0% and 3% of particles from the DI and DE datasets, respectively, are consistent with the AI complex. All final 2D class averages are shown in **Supplemental Figs. S11 and S13**. **C**. 3D reconstructions and molecular models of POLA1_cat_ and PRIM2_C_ bound to the 3′- and 5′-ends of the primers, respectively. The positions of the interdomain linkers are indicated with arrowheads. **D,E**. Distinct configurations of the DI and DE complexes. Models were constructed from four rigid bodies, comprised of the POLA1_cat_/PRIM2_C_ models shown in panel C and the PRIM1/PRIM2_N_/POLA1_C_/POLA2_C_ models shown in Fig. 2, and docked into the low-resolution 3D reconstructions shown in **Supplemental Figs. S19 and S20**. **D**. DI and DE complexes oriented as in the 2D class averages. **E**. DI and DE complexes aligned on the tetramer core.

Relative to polα–primase in the absence of substrate, the 2D class averages determined in the presence of the DI and DE substrates showed increased configurational heterogeneity and dramatic configurational rearrangements (**Fig. 3B and Supplemental Figs. S10–S13**). The vast majority of the class averages lacked clear density for part of the complex, indicating a large degree of flexibility and conformational/configurational heterogeneity remains when substrates are bound. These class averages could be divided into two groups: those with density for POLA1_cat_ and PRIM2_C_ bound to substrate (hereafter referred to as the DNA synthesis assembly, or DSA), and those with density for everything else (i.e., the tetramer core). With the DI substrate, 2D classes showed 67% of particles with clear density for only the DNA synthesis assembly, 26% for only the tetramer core, and ∼6% for polα–primase in various configurations. With the DE substrate, 2D classes showed a similar distribution, but with fewer particles aligned on the DNA synthesis assembly and more particles aligned on the tetramer core. Only 0 and 3% of particles with the DI and DE substrates, respectively, were found to be in the AI configuration.

To better characterize interactions with the substrates, we performed focused 3D refinement using masks around POLA1_cat_, PRIM2_C_, and the primer/template (**Fig. 3C, Supplemental Figs. S10, S12, S14, and S15, and Supplemental Tables S3 and S4**). With both the DI and DE substrates, the 3.3-Å and 4.4-Å reconstructions showed POLA1_cat_ and PRIM2_C_ bound to opposing ends of the duplexes (**Fig. 3C**), indicating that interactions with PRIM2_C_ are maintained during transfer of the primer to polα and throughout DNA synthesis. Focused 3D refinement of only POLA1_cat_ and the DNA/DNA portion of the DE substrate improved the resolution of the reconstruction to 3.5 Å (**Supplemental Figs. S12 and S15 and Supplemental Table S4**). In both DSA structures, POLA1_cat_ is in a closed ternary complex with the primer 3′-end and the incoming dGTP, and PRIM2_C_ is bound to the primer 5′-triphosphate. With the DI substrate, POLA1_cat_ and PRIM2_C_ are in contact with one another, albeit with the entire subunit interface consisting of a single interaction between two loops (**Fig. 3C and Supplemental Fig. S16**). With the longer DE substrate, the two domains are separated by a full turn of the RNA-DNA/DNA duplex. 3D variability analysis of these DI and DE subcomplexes revealed dissociation of PRIM2_C_ from a subset of the particles, as well as bending of the oligonucleotide duplexes (**Supplemental Figs. S17 and S18**). With the shorter DI substrate, this bending altered the protein-protein interface between POLA1_cat_ and PRIM2_C_.

Prompted by the seeming malleability and small size of the subunit interface, we modeled POLA1_cat_ and PRIM2_C_ with shorter RNA primers (**Supplemental Fig. S16**). With 8-mer and 7- mer primers, we observed only minor clashes between side chains, which appear unlikely to prevent DNA initiation. Conversely, we saw severe steric overlap between POLA1_cat_ and PRIM2_C_ with a 6-mer primer. These models explain the previous observation that polα can extend RNA primers as short as seven nucleotides in length ^12^. Additionally, binding of the nascent primer by PRIM2_C_ during DNA elongation explains the earlier observation that the primer 5′-triphosphate affects not only RNA synthesis but also DNA synthesis ^43^, as well as the recent finding that PRIM2_C_ increases the processivity of polα–primase ^44^.

For simultaneous binding of the primer ends to be maintained throughout DNA synthesis, as the primer increases in length, large-scale protein conformational changes are necessary. Because of the extreme conformational and configurational heterogeneity inherent in the polα– primase complex, we were unable to generate reconstructions of the full substrate-bound complex by multiclass ab initio reconstruction and heterogeneous refinement. However, we were able to generate reconstructions from subsets of particles manually selected from 2D class averages (**Supplemental Figs. S19 and S20 and Supplemental Tables S3 and S4**). Rigid-body docking of the DSA and TC subcomplexes into these low-resolution reconstructions enabled us to model two configurations of the complete polα–primase complex bound to DI substrate and two configurations bound to DE substrate (**Fig. 3D,E, Supplemental Figs. S19 and S20, and Supplemental Tables S3 and S4**). Comparison of the four substrate-bound configurations showed that the DNA synthesis assembly rotates by as much as 180° with respect to the tetramer core, with the pivot point occurring between PRIM2_C_ and PRIM2_N_ (**Fig. 3E**). In addition to this swiveling, the DNA synthesis assembly rotates by nearly 180° with respect to the helical axis of the oligonucleotide duplexes. In all models of the four-subunit complex, the substrate-bound configurations appear to be stabilized by interactions between the DNA synthesis assembly and the tetramer core. However, the variability of these interactions, as well as the large proportion of 2D class averages showing density for only one or the other subcomplex, suggests these protein-protein contacts are weak and transient. Thus, in the presence of primer/template substrates representing two key stages of DNA synthesis, polα– primase adopts a wide range of configurations characterized by concerted movement of POLA1_cat_, PRIM2_C_, and the oligonucleotide duplex relative to the tetramer core.

### Primase enhances an intrinsic counting ability in polα

Whether counting of DNA nucleotides is an inherent property of polα or only a property of the complete polα–primase complex is unclear ^32,34,44,50,51^. Our finding that PRIM2_C_ remains bound to the primer after transfer to polα reveals a means by which primase might participate in counting during DNA synthesis. We theorized that binding of the substrate by PRIM2_C_ would alter DNA elongation by indirectly tethering POLA1_cat_ to the primer via the tetramer core. To investigate this theory, we compared the RNA and DNA synthesis activities of the polα–primase complex with a complex in which the interdomain linker between POLA1_cat_ and POLA1_C_ was severed, constructed by mixing purified POLA1_cat_ and ΔPOLA1_cat_ (PRIM1/PRIM2/POLA1_C_/POLA2) (**Fig. 4A and Supplemental Fig. S21**). As a control, we measured the RNA and DNA synthesis activities of POLA1_cat_ and ΔPOLA1_cat_ separately. Primer extension was performed by adding radiolabeled ATP or dATP and unlabeled NTPs and dNTPs to a 5′-triphosphorylated 6-ribonucleotide primer annealed to a 99-deoxyribonucleotide template (**Fig. 4B and Supplemental Table S2**). POLA1_cat_ alone generated extension products of predominantly 17–33 nucleotides, with the main product being 30 nucleotides in length (**Fig. 4A,C**). Similar to previous observations with primase alone ^21,29^, ΔPOLA1_cat_ rapidly formed products of nine nucleotides before abruptly terminating at a 10-mer primer (**Fig. 4A**). The severed complex generated by combining POLA1_cat_ and ΔPOLA1_cat_ afforded products representing an admixture of the individual control reactions (**Fig. 4A,C and Supplemental Fig. S22**). In contrast, intact polα–primase generated fewer extension products outside of 30 nucleotides (**Fig. 4A,C**), and produced the 30-mer product at a dramatically faster rate (**Fig. 4D**). However, the total activity of polα–primase was notably less than that of the other constructs. We speculate this reduced activity results from a substantial proportion of the polα– primase molecules being in the AI configuration, and therefore unable to perform primer elongation without undergoing a dramatic configurational transition. Collectively, these data demonstrate an intrinsic—albeit approximate—counting ability in polα that is improved by the physical linkage between POLA1_cat_ and the remainder of the complex, reinforcing the preference of polα–primase to generate a 30-nucleotide primer.

**Figure 4.**
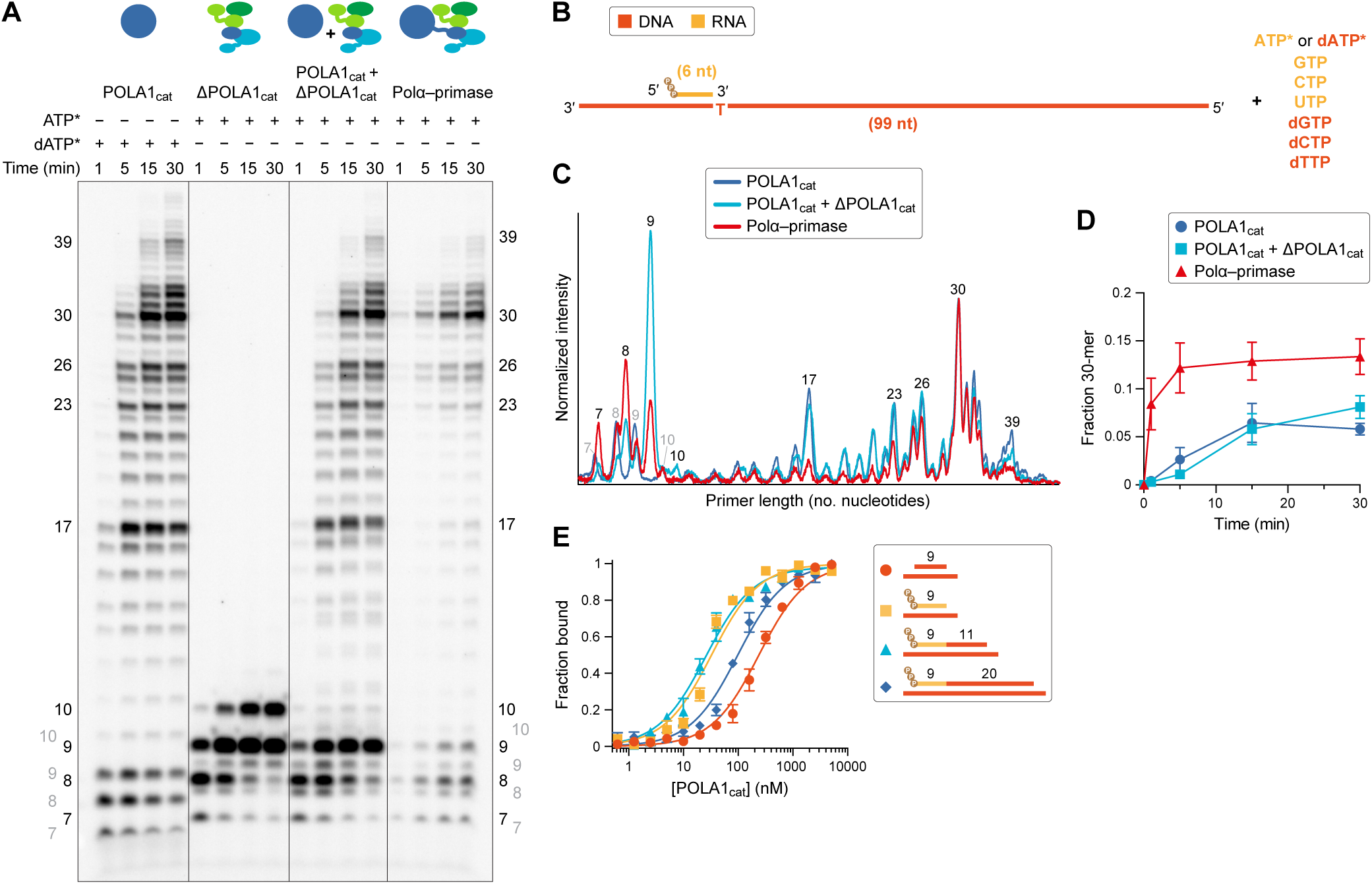
Primer/template binding and elongation. **A**. Separation and detection of RNA and RNA-DNA elongation products by denaturing gel electrophoresis and autoradiography. Intermediate and doublet bands result from a mixture of RNA and DNA incorporation at positions 7–10 (**Supplemental Fig. S22**). **B**. Substrate schematic. Triphosphate moieties are depicted as brown circles. [α-^32^P]-labelled ATP and [α-^32^P]-labelled dATP are only incorporated at position 7 in the primer. **C**. Quantitation of the 30-min time points from the gel in panel A. The intensities of the elongation products are normalized to the intensity of the 30-mer. **D**. Kinetics of 30-mer formation. **E**. Binding of DNA/DNA, RNA/DNA, and RNA-DNA/DNA duplexes by POLA1_cat_. Data in panels D and E are presented as the mean ± SEM from three replicate experiments.

To better understand the approximate counting ability of polα, we performed binding assays with POLA1_cat_ and DNA/DNA, RNA/DNA, and two RNA-DNA/DNA duplexes (**Fig. 4E and Supplemental Table S5**). Consistent with previous findings ^32,45^, we observed significantly tighter binding of an RNA/DNA substrate than a corresponding DNA/DNA substrate. Addition of 11 nucleotides of DNA to the RNA primer, creating an RNA-DNA/DNA substrate, had no effect on binding affinity. However, a further addition of nine nucleotides of DNA, producing a longer RNA-DNA/DNA duplex, resulted in significantly weaker binding, approaching that of the DNA/DNA substrate. For both RNA-DNA/DNA duplexes, the DNA/DNA regions were sufficiently long to fully accommodate the binding footprint of the polymerase catalytic core. Thus, these findings indicate the affinity of POLA1_cat_ for chimeric primers is dependent on not only the length of the DNA/DNA region of the duplex but also the proximity of the duplex end to the RNA/DNA region, providing a basis for counting and the termination of DNA synthesis.

### Primer/template conformation modulates substrate recognition

Consistent with the results from our binding assays, the intrinsic counting ability of polα has been proposed to result from conformation-dependent recognition of RNA/DNA and DNA/DNA duplexes ^32,45^. Because of the relatively high resolutions of the DSA reconstructions, we were able to determine the conformations of the 5′-triphosphorylated RNA/DNA and RNA-DNA/DNA substrates in the DI and DE subcomplexes and to evaluate the details of their recognition by POLA1_cat_ and PRIM2_C_. In both structures, POLA1_cat_ was observed in a closed conformation in which the fingers domain was fully engaged with an incoming dGTP and active-site residues within the palm domain (Asp859, Phe860, and Asp1000) coordinated a catalytic Mg^2+^ ion (**Fig. 5A,B and Supplemental Fig. S23**). The thumb domain was also fully engaged with both the RNA/DNA and RNA-DNA/DNA duplexes (**Fig. 5A,C**). To our knowledge, the DE subcomplex represents the first structure of POLA1_cat_ with the thumb and fingers domains engaged with a DNA/DNA duplex and an incoming dNTP (**Supplemental Fig. S24**). In both subcomplexes, POLA1_cat_ induces a local A-form conformation in the duplexes that positions and orients the 2′,3′-dideoxycytidine at the 3′-end of the primers for phosphodiester bond formation with dGTP (**Fig. 5B,E,F**). Similar catalytic arrangements have been observed in structures of polδ and polε ^52,53^. However, POLA1_cat_ possesses a wider substrate-binding cleft that results from structural differences in the thumb domain (**Supplemental Fig. S25**). This wider cleft has been proposed to complement the wider diameter of an A-form RNA/DNA duplex ^32,45^. In both of our structures, we observed intermediate AB-form conformations ^54,55^, or nearly intermediate AB-form conformations, over most of the region of the duplexes bound by POLA1_cat_ (**Fig. 5E,F**).

**Figure 5.**
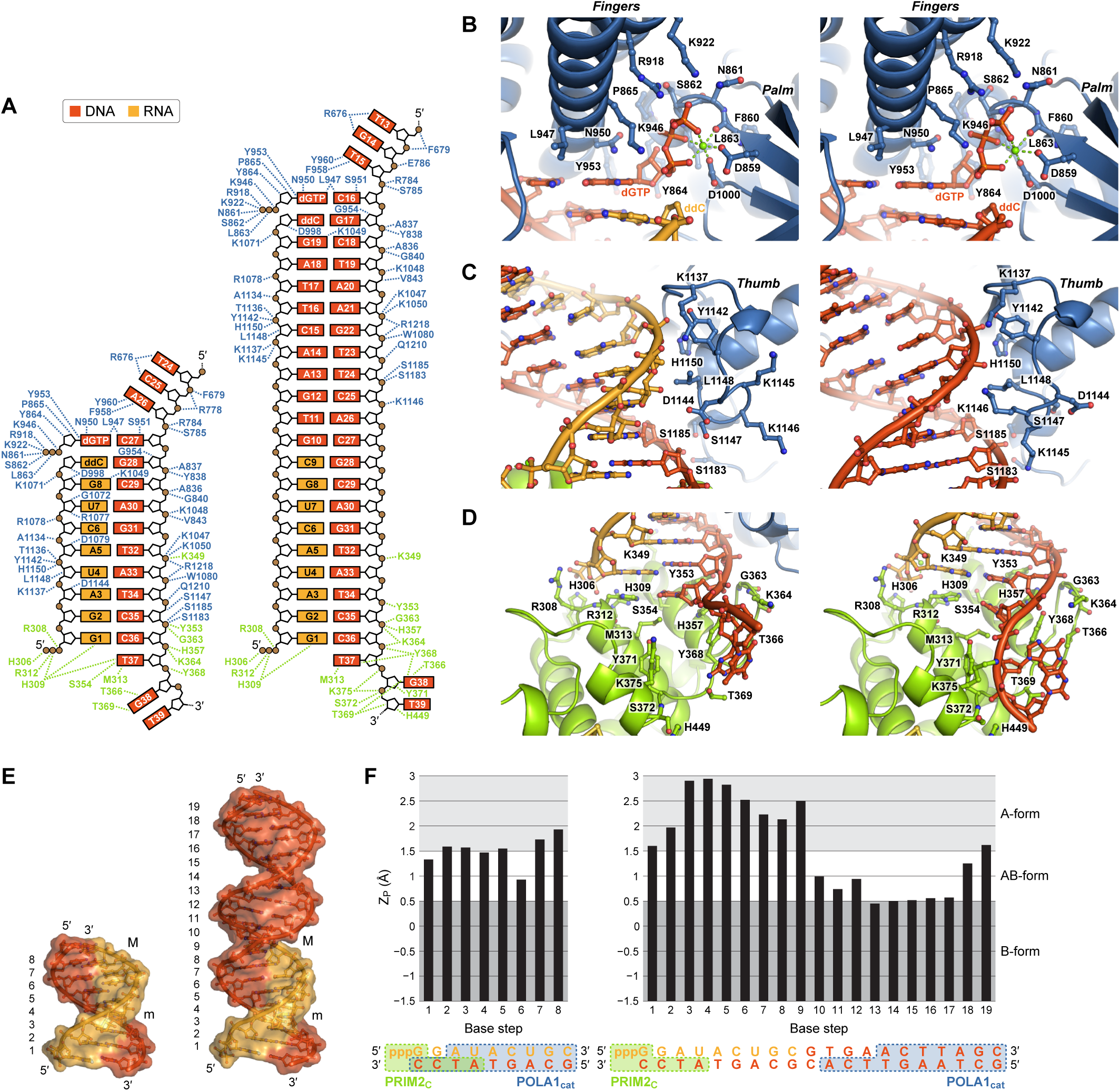
Substrate recognition and duplex remodeling. In each panel, the DI and DE structures are positioned on the left and the right, respectively. **A**. Schematic of substrate-binding contacts in POLA1_cat_ (blue) and PRIM2_C_ (green). **B**. Recognition of an incoming dGTP by POLA1_cat_. Phosphodiester bond formation is prevented by a 2′,3′-dideoxycytidine (ddC) at the end of the primers. **C**. Recognition of RNA/DNA and DNA/DNA duplexes through substrate-adaptive conformational changes in POLA1_cat_. **D**. Recognition of distinct primer/template conformations by PRIM2_C_. **E**. RNA/DNA and RNA-DNA/DNA duplexes extracted from the DI and DE subcomplexes. Major (M) and minor (m) grooves are indicated. Numbers to the left of the duplexes indicate the base step starting from the 5′-end of the primers. **F**. Analysis of duplex conformation. Z_P_ is the interstrand phosphate-phosphate distance after projection onto the helical (Z) axis. The regions of the duplexes bound by POLA1_cat_ (blue) and PRIM2_C_ (green) are shown below the plots.

Importantly, small but significant differences in the interactions between the thumb domain and the DI and DE substrates illustrate how POLA1_cat_ accommodates different minor groove geometries (**Fig. 5A,C,E,F**). With the DI substrate, three aspartate residues (Asp998, Asp1079, and Asp1144) form hydrogen-bonding interactions with 2′-hydroxyl groups along the RNA primer. With the DE substrate, conformational changes at the tip of the thumb domain (residues 1142–1148) pull Asp1144 away from the chimeric primer while directing Lys1145 and Lys1146 toward the relatively narrow minor groove in the same DNA/DNA region of the duplex. These new hydrogen-bonding interactions likely partially offset the loss of RNA-specific hydrogen bonds with the DE substrate. While RNA/DNA and DNA/DNA duplexes generally adopt A-form and B-form geometries ^56^, respectively, the intermediate AB-form conformations observed with both the DI and DE substrates suggest POLA1_cat_ remodels the primer/template during DNA synthesis. The local propensity of the duplex to adopt this intermediate conformation is likely correlated with substrate affinity and, consequently, with processivity ^32,45^.

PRIM2_C_ has been shown to increase the processivity of polα–primase during DNA synthesis ^44^. The two DSA structures suggest that this increased processivity is the result of continued association of PRIM2_C_ with the primer/template during DNA elongation. In both the DI and DE subcomplexes, interactions between PRIM2_C_ and the primer strand are identical (**Fig. 5A,D**). Three positively charged residues (His306, Arg308, and Arg312) at the N-terminus of helix α15 surround the 5′-triphosphate and offset the negative charge, and His309 stacks against the 5′-terminal nucleobase. Surprisingly, interactions between PRIM2_C_ and the template strand are markedly different in the two subcomplexes, despite PRIM2_C_ binding the same RNA/DNA portion of both substrates. In the DI structure, the DNA template is positioned such that only the last paired nucleotide, which is not coplanar with the 5′-terminal nucleotide in the RNA primer, and the first unpaired nucleotide form extensive contacts with PRIM2_C_. Rotation of the phosphodeoxyribose backbone swings the second and third unpaired nucleotides away from PRIM2_C_ and toward POLA1_cat_. In the DE structure, the DNA template is positioned such that the last paired nucleotide, which is coplanar with the 5′-terminal nucleotide in the RNA primer, and the first, second, and third unpaired nucleotides form extensive contacts with PRIM2_C_. Relative to the DI substrate, interactions with several residues (Tyr353, His357, Lys364, and Tyr368) in the flexible template-binding loop (residues 359–372) are shifted by one nucleotide toward the 5′-end of the template strand as a result of increased base pair inclination in the DE substrate. Virtually identical interactions were observed in an X-ray crystal structure of human PRIM2_C_ bound to a short RNA/DNA duplex ^34^ (**Supplemental Fig. S26**).

The unique interactions that we observe with the DI substrate appear to result from local duplex remodeling by POLA1_cat_, which causes the RNA/DNA duplex to have less A-form character, as quantified by the Z_P_ parameter ^54,55,57^, than either the RNA/DNA portion of the DE substrate or the RNA/DNA duplex in the crystal structure (**Fig. 5D,E,F and Supplemental Fig. S26**). In addition to reduced A-form character, the DI substrate has lower base pair inclination, which pulls the 3′-overhang away from PRIM2_C_. The affinity of PRIM2_C_ for the primer/template was previously shown to depend on both the 5′-triphosphate and the 3′-overhang ^34,41^. Thus, tight binding of the substrate appears to require substantial inclination of the base pairs at the end of duplex. Together, these data indicate that like POLA1_cat_, the affinity of PRIM2_C_ for the primer/template is modulated by the nucleic acid conformation, which is imposed by both the DNA composition of the primer and binding by the polymerase catalytic core.

### Loose association of full-length primers allows for reversion to the auto-inhibitory configuration

The finding that substrate recognition by both POLA1_cat_ and PRIM2_C_ is dependent on duplex conformation has implications for the mechanism of DNA counting. To investigate how duplex conformation affects the preference of polα–primase to generate 30-nucleotide primers, we designed a substrate to visualize the complex at the end of DNA synthesis (DNA termination, DT). The DT primer is comprised of nine ribonucleotides followed by 20 deoxyribonucleotides (**Fig. 6A, Supplemental Fig. S9, and Supplemental Table S2**) and includes the same 5′-triphosphate group as the DI and DE primers. A ternary DT complex containing polα–primase, primer/template, and incoming nucleotide was trapped using 2′,3′- dideoxycytidine to terminate the primer strand and prevent phosphodiester bond formation with dGTP.

**Figure 6.**
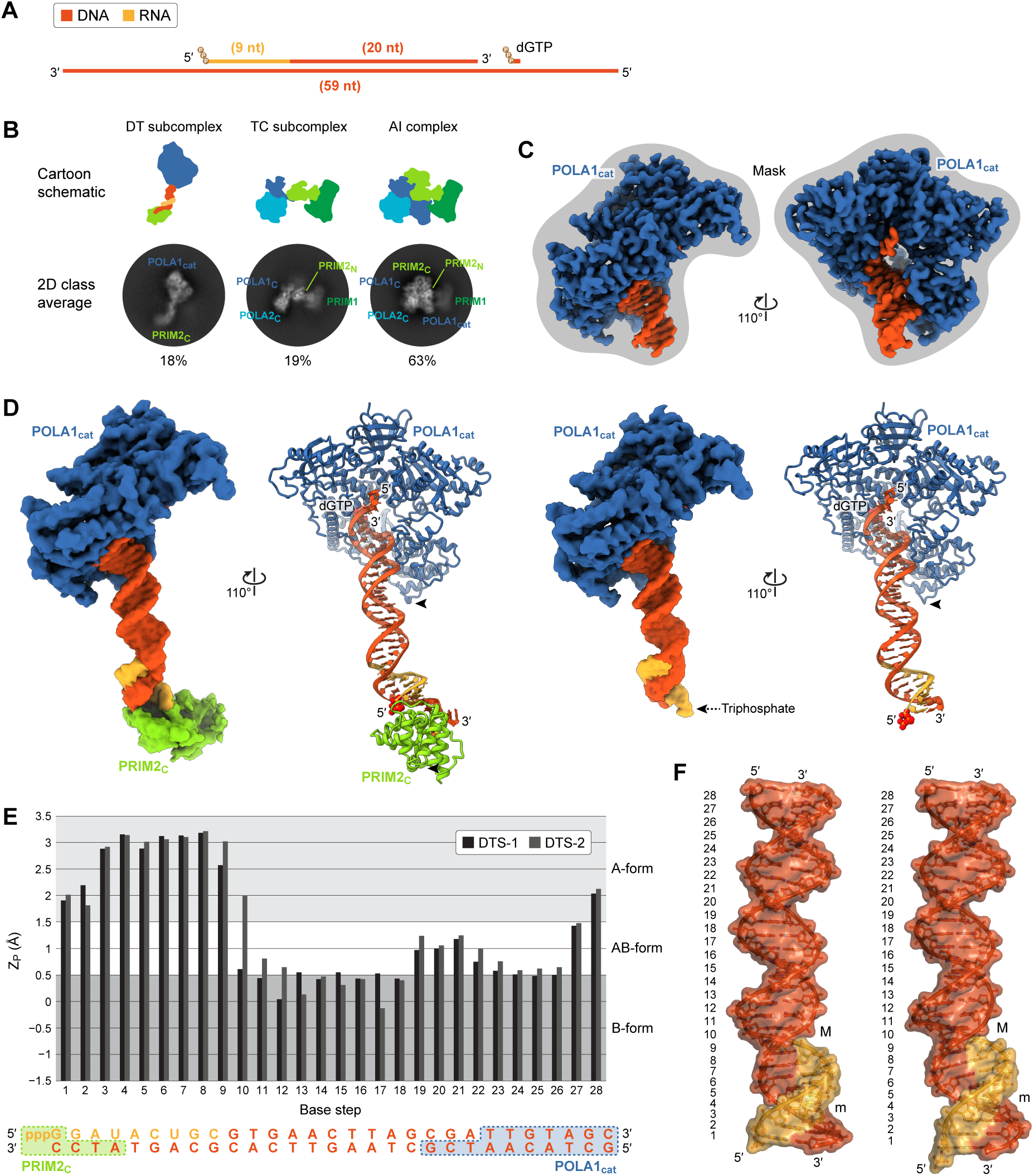
Configurational rearrangements upon DNA termination. **A**. Substrate schematic. Triphosphate moieties are depicted as brown circles. The primer terminates in a 2′,3′- dideoxycytidine to prevent incorporation of dGTP. **B**. Representative 2D class averages and corresponding subunit schematics. The percentage of particles belonging to the DT subcomplex, the TC subcomplex, or the AI complex is indicated below the class averages. No 2D class averages were observed with clear density for the complete DT complex. All final 2D class averages are shown in **Supplemental Fig. S28**. **C**. Masked 3D reconstruction of POLA1_cat_ bound to a full-length primer/template. **D**. 3D reconstructions and molecular models of POLA1_cat_ bound to a full-length primer/template with (DT subcomplex 1, left) and without (DT subcomplex 2, right) simultaneous association of PRIM2_C_. The positions of the interdomain linkers are indicated with arrowheads. **E**. Analysis of duplex conformation. Z_P_ is the interstrand phosphate-phosphate distance after projection onto the helical (Z) axis. The regions of the duplexes bound by POLA1_cat_ (blue) and PRIM2_C_ (green) are shown below the plots. **F**. RNA-DNA/DNA duplexes extracted from DT subcomplex 1 (left) and DT subcomplex 2 (right). Major (M) and minor (m) grooves are indicated. Numbers to the left of the duplexes indicate the base step starting from the 5′-end of the primers.

As with the DI and DE substrates, a substantial proportion of 2D class averages showed density for only the DNA synthesis assembly or the tetramer core (**Fig. 6B and Supplemental Figs. S27 and S28**), consistent with large-scale conformational and configurational dynamics. Unlike with the shorter substrates, however, no class averages showed density for the complete polα–primase complex bound to substrate. This level of heterogeneity suggests that the interdomain linkers in POLA1 and PRIM2 remain slack when bound to a full-length primer, as taught linkers would seemingly reduce the overall flexibility of the DT complex. Surprisingly, 63% of particles were grouped into 2D classes with density clearly representative of the AI complex. Protein-protein contacts in this configuration sterically occlude substrate recognition by both POLA1_cat_ and PRIM2_C_, indicating the DT substrate is not bound in most particles. With the DI and DE substrates, only 0 and 3% of particles, respectively, were found to be in the AI configuration, while in the absence of substrate, 69% of particles were found to be in this configuration. While this type of comparative quantitation of cryo-EM data has the potential for bias, this dramatic increase in AI particles in the presence of DT substrate suggests substantially weaker binding of the longer chimeric primer by both POLA1_cat_ and PRIM2_C_.

To better understand the structural basis for weaker association of the DT substrate, we performed focused 3D refinement of the DT subcomplex. Using a mask around POLA1_cat_ and the DNA/DNA region of the primer/template, we generated a 2.9-Å reconstruction with clear density for POLA1_cat_ in a closed conformation, with the thumb and fingers domains fully engaged with the duplex and an incoming dGTP (**Fig. 6C, Supplemental Figs. S27 and S29, and Supplemental Table S6**). 3D variability analysis of the complete DT subcomplex, including POLA1_cat_, PRIM2_C_, and the full-length primer/template, revealed duplex bending similar to that seen in the DE subcomplex, as well as dissociation of PRIM2_C_ from the substrate (**Supplemental Fig. S30**). However, the degree to which PRIM2_C_ is disengaged from the primer/template appears to be substantially greater with the DT substrate. Consistent with this observation, 3D classification sorted most particles aligned on the DT subcomplex into classes with little to no density for PRIM2_C_ (**Supplemental Fig. S27**). Following classification, focused 3D refinement provided 3.8- and 3.6-Å reconstructions of the DT subcomplex with (DT subcomplex 1, or DTS-1) and without (DT subcomplex 2, or DTS-2) PRIM2_C_ engaged with the substrate (**Fig. 6D, Supplemental Figs. S27 and S29, and Supplemental Table S6**). The overall conformations of the duplexes in these structures, as quantified by the Z_P_ parameter, are quite similar (**Fig. 6E**). However, the base pair inclination in the RNA/DNA portion of DTS-2 appears to be reduced relative to DTS-1 (**Fig. 6F and Supplemental Fig. S31**). While the degree of certainty associated with this observation is limited by the local resolution of the reconstructions (**Supplemental Fig. S29**), this conformational difference is consistent with our conclusion that tight substrate binding by PRIM2_C_ is dependent on the base pair inclination at the end of the duplex. Conversely, the origins for weaker binding of the DT substrate by POLA1_cat_ are not apparent in the structures. In particular, all structures of the DNA synthesis assembly reveal POLA1_cat_ has remodeled the bound region of the duplex—whether it is RNA/DNA or DNA/DNA—to achieve similar intermediate AB-form conformations (**Supplemental Fig. S31**). However, remodeling of this type requires an input of free energy, which in turn reduces the free energy associated with binding affinity. Thus, primer/template duplexes predisposed to adopt intermediate AB-form conformations, like those in the remodeled substrates, require less free energy to be remodeled, and therefore will be bound with higher affinity.

## Discussion

Multiple, seemingly conflicting models have been proposed to explain the counting abilities of primase and polα ^32–34,41,42,44–47^. Our structures of polα–primase create a framework to reconcile and integrate these models, providing a comprehensive mechanistic understanding of nucleotide counting (**Fig. 7A,B**). We show that in the absence of nucleic acid substrates or protein partners, polα–primase exists primarily in a compact, auto-inhibitory state, but spontaneous dissociation of POLA1_cat_ and PRIM2_C_ initiates a series of dramatic structural rearrangements necessary for chimeric primer synthesis across two spatially separate active sites. In all current models of RNA synthesis, primer formation is initiated by PRIM1 and PRIM2_C_ bringing together two NTPs on a single-stranded DNA template. In the linker model, elongation of the primer continues until halted by full extension of the PRIM2 interdomain linker ^41,42,47^. In the collision model, elongation is ceased by a steric clash between PRIM2_N_ and PRIM2_C_ ^34^. As previously discussed ^34^, the linker model is not consistent with domain arrangement in the tetramer core. Conversely, the collision model is supported by structural data and structure-based modeling ^34^, and provides a convincing rationale for termination of RNA synthesis, but strictly at 9–10 nucleotides. Both models fail to explain previous observations showing extension of RNA primers as short as seven nucleotides in length by polα ^12^.

**Figure 7.**
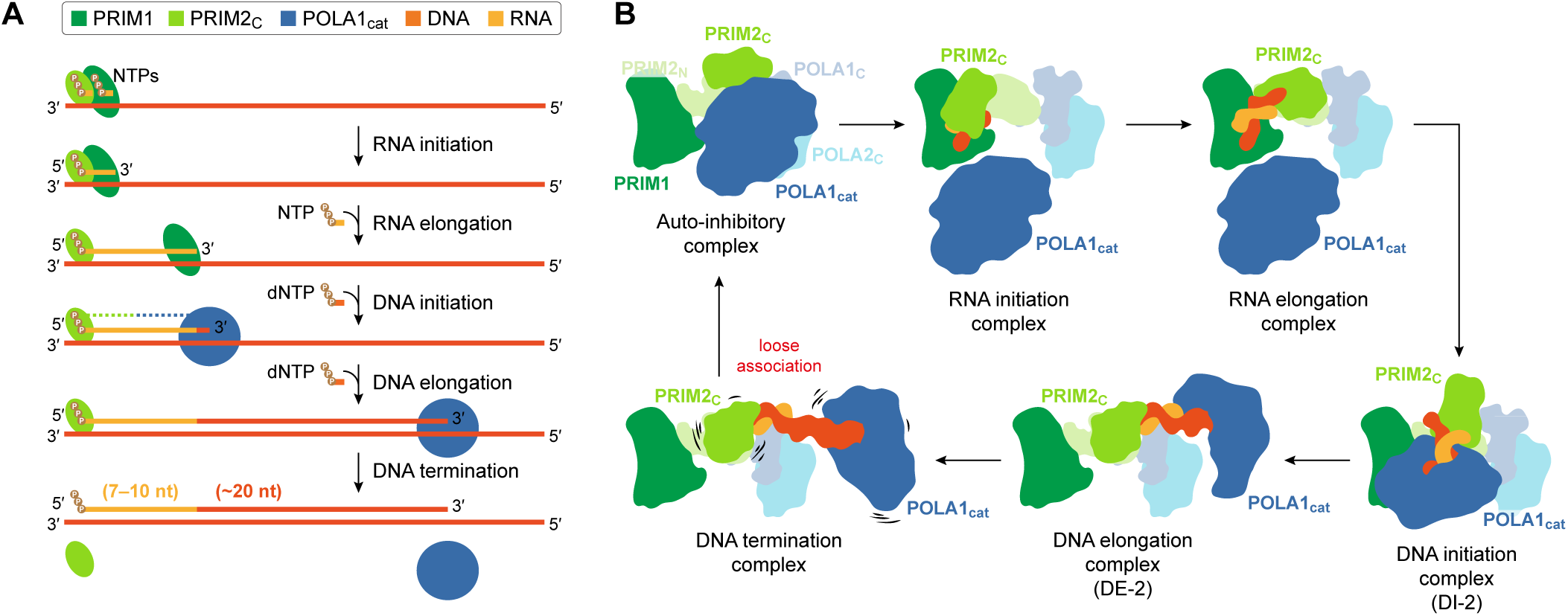
Structural rearrangements during chimeric primer synthesis. **A**. Stages of RNA and DNA synthesis. PRIM2_C_ remains bound to the 5′-end of the primer during all stages of synthesis. The transition from RNA elongation to DNA initiation occurs when the 3′-end of the primer is transferred from PRIM1 to POLA1_cat_. After generation of a full-length primer, DNA elongation is terminated by dissociation of polα–primase from the primer/template. Protein-protein contacts between PRIM2_C_ and POLA1_cat_ are represented by a dashed line. **B**. Subunit schematics showing configurational rearrangements throughout the catalytic cycle of primer synthesis. Schematics are aligned on the tetramer core (PRIM1/PRIM2_N_/POLA1_C_/POLA2_C_). The RNA initiation and RNA elongation configurations were derived from hypothetical models in which the positions of POLA1_cat_ are indeterminate and the positions of PRIM2_C_ are dictated by the geometric requirements of phosphodiester bond formation in the active site of PRIM1. The DNA initiation and DNA elongation configurations correspond to DI-2 and DE-2, respectively (Fig. 3). The DNA termination configuration was derived from a hypothetical model in which the DE subcomplex in DE-2 was replaced by DT subcomplex 1 while retaining the position of PRIM2_C_. Loose association of POLA1_cat_ and PRIM2_C_ with the full-length primer/template is indicated with wobble lines. At each stage of primer synthesis, the configuration depicted is likely representative of a broad configurational/conformational distribution.

Based on our structure of the DI subcomplex, as well as previous biochemical data, we propose a competition model for RNA counting and primer transfer. PRIM2_C_ has been suggested to anchor the primer during its transfer between primase and polα active sites by remaining bound to the primer 5′-end ^34,37,44^. Moreover, the highly distributive nature of RNA and DNA synthesis by PRIM1 and POLA1_cat_ in the absence of PRIM2_C_ ^29,44,58^ suggests that PRIM2_C_ acts as a hub to facilitate re-association of PRIM1 and POLA1_cat_ with the primer. Our structures confirm that PRIM2_C_ remains bound to the 5′-end of the primer after transfer to polα, and provide evidence that a dynamic protein-protein interface allows PRIM2_C_ and POLA1_cat_ to simultaneously bind RNA primers of seven or more nucleotides, while transfer of RNA primers less than seven nucleotides in length to POLA1_cat_ would be sterically blocked by PRIM2_C_. Thus, the likely driving factor in the primase-to-polymerase switch is a competition between PRIM1 and POLA1_cat_ for the 3′-end of the primer, rather than termination of RNA synthesis at a maximal length of 10 nucleotides. In this competition model, steric collision between PRIM2_N_ and PRIM2_C_ would serve only as a failsafe to terminate RNA synthesis, ensuring longer stretches of RNA (> 10 nts) are not incorporated into the genome, should POLA1_cat_ not outcompete PRIM1 for the primer.

Our finding that PRIM2_C_ remains bound to the primer after transfer to polα has far-reaching implications for DNA synthesis, including the preference of polα–primase to generate chimeric primers of ∼30 nucleotides. Simultaneous association of both PRIM2_C_ and POLA1_cat_ is a prerequisite for the linker model of DNA termination, which posits that the POLA1 and PRIM2 interdomain linkers prevent separation of PRIM2_C_ and POLA1_cat_ beyond the distance associated with an ∼30-mer primer ^44,47^. However, we observed seemingly weaker binding of chimeric primers by both POLA1_cat_ and PRIM2_C_ as the DNA composition of the primer increased (as measured by direct binding assays using POLA1_cat_ and implied by the proportion of particles in the AI configuration). Moreover, if termination were defined only by the length of the interdomain linkers in POLA1 and PRIM2, we would expect to observe a stable complex between polα– primase and the DT substrate, likely with well-ordered, taught linkers. Additionally, the linker model is unable to provide a rationale for the intrinsic counting ability of POLA1_cat_, which we and others have observed ^32,50^. By contrast, the affinity model of DNA termination, which proposes that POLA1_cat_ preferentially recognizes A-form duplexes typical of RNA/DNA hybrids, and loses affinity for the substrate as the duplex becomes more B-form throughout DNA synthesis ^32,45^, provides a mechanistic explanation for the innate counting ability of the polymerase catalytic core. Our data expand upon the affinity model by suggesting a role for preferential binding of A-form duplexes by PRIM2_C_, and by providing a possible structural basis for this specificity. Thus, we propose that DNA termination is the result of both POLA1_cat_ and PRIM2_C_ losing affinity for the primer/template as the B-form propensity of the duplex increases during DNA synthesis. Because the linker model requires both POLA1_cat_ and PRIM2_C_ to remain bound to the substrate, the interdomain linkers are likely to play, at most, a failsafe role in restricting primer length, and thereby limiting the mutagenic potential of polα.

The dramatic structural rearrangements that we observe are consistent with the idea that flexible tethering of POLA1_cat_ and PRIM2_C_ to the tetramer core is necessary for transfer of the nascent primer from primase to polα ^26,34^. As we demonstrate here, these rearrangements are also necessary for simultaneous binding of the primer ends by POLA1_cat_ and PRIM2_C_ during DNA elongation, as the primer increases in length. Presumably, this plasticity also facilitates interaction with the many protein partners with which polα–primase is known to interact at replication forks and telomeres. At replication forks, polα–primase interacts with several proteins associated with the replisome, including MCM10, AND-1, RPA, RFC, polδ, and the CMG helicase ^19,36,59^, as well as multiple nucleosome core proteins ^60,61^. Despite this extensive network of contacts, polα–primase appears to remain conformationally dynamic ^62,63^. Moreover, recent structures of polα–primase bound to a replisome complex revealed a manner of flexible association ^36^ that is unlikely to restrict even the extraordinarily broad range of motion that we observed in our structures.

Polα–primase appears to remain similarly dynamic at telomeres, as three distinct configurations of the polα–primase/CST complex bound to single-stranded DNA have been observed ^37,48,49^. The first structure, termed a recruitment complex, showed polα–primase in the AI configuration with CST primarily associated via interactions with POLA1_C_ ^48^. The second structure, designated a pre-initiation complex, showed polα–primase in an extended configuration in which POLA1_cat_ and the tetramer core are scaffolded by CST ^37^. In this configuration, the PRIM1 and POLA1_cat_ active sites are locked in opposing orientations, with PRIM2_C_ free to move between them. This arrangement was proposed to facilitate RNA synthesis and transfer of the RNA primer from primase to polα. Consistent with this notion, CST has been shown to enhance primer synthesis, without substantially affecting primer length ^37,64,65^. Moreover, docking of our DSA structures into the pre-initiation complex revealed additional interactions between the thumb domain of POLA1_cat_ and the CTC1 and STN1 subunits of CST (**Supplemental Fig. S32**), which may stabilize the closed, substrate-bound conformation of the polymerase catalytic core, supporting POLA1_cat_ in its competition with PRIM1 for the RNA primer. The positions of our structures in these models, however, suggest DNA elongation would lead to minor and then severe steric clashes between PRIM2_C_ and the tetramer core as the primer reaches lengths of 10 and 18 nucleotides, respectively. In this configuration, these clashes are likely to prevent synthesis of a full-length primer. This conclusion is supported by the third structure of polα–primase/CST, which showed POLA1_cat_ associated with CST, as in the pre-initiation complex, but with the tetramer core disengaged ^49^. Together, these observations suggest the tetramer core is not tightly bound to CST and dissociates prior to or during DNA elongation by POLA1_cat_. Based on the evolutionarily relationship between CST and RPA ^66^, we believe RPA likely scaffolds polα–primase in a similar manner, explaining RPA-dependent stimulation of primer synthesis by polα–primase ^67^. How these interactions with RPA and CST are coordinated with those of other proteins at replication forks and telomeres, as well as the effects these interactions may have on the conformational and configurational dynamics of polα–primase and the mechanism of chimeric primer synthesis, remain to be determined.

## Conclusion

Determining structures of polα–primase with multiple catalytic substrates, along with performing structure-based biochemical and biophysical analyses, has enabled us to propose a comprehensive model for primer synthesis. This model is built upon the large body of available structural and biochemical data while unifying and refining prevailing understanding of the mechanistic basis for polα–primase function. Our structural data reveal that polα–primase both remodels and reads the conformation of its primer/template substrate, which explains its unique ability to regulate primer length (i.e., count). These data also reveal that polα–primase retains a remarkable degree of conformational and configurational flexibility that is crucial to the intricate functions of this complex and dynamic protein machine. A greater appreciation of the type and degree of intrinsic flexibility that we have observed for polα–primase is anticipated as active eukaryotic replisomes and other cellular machinery come into view at ever increasing resolution.

## Methods

### Cloning and expression

The genes encoding *Xenopus laevis* PRIM1 (NCBI accession NP_001080819.1), PRIM2 (NCBI accession XP_018121015.1), POLA1 (NCBI accession NP_001082055.1), and POLA2 (NCBI accession NP_001086972.1) were initially cloned into pFastBac-HTa or pFastBac-HTb expression vectors (Invitrogen). PRIM1 was then subcloned into a modified pET-19b expression vector (Novagen) encoding an HRV 3C-cleavable decahistidine fusion tag. PRIM2 was subcloned into a modified pRSFDuet-1 expression vector (Novagen). And, POLA1 and POLA2 were subcloned into an extensively modified, polygenic pFastBac-1 expression vector (Invitrogen) encoding TEV-cleavable StrepII-GFP (POLA1) and hexahistidine-MBP (POLA2) fusion tags. Protein expression and purification were facilitated by truncating the gene encoding POLA1 to remove the N-terminal tail (residues 1–334). To generate a POLA1_cat_ construct, the catalytic core of POLA1 (residues 335–1254) was subcloned into a modified pET-27b(+) expression vector (Novagen) encoding an HRV 3C-cleavable hexahistidine-SUMO fusion tag. To generate a ΔPOLA1_cat_ construct, the C-terminal domain of POLA1 (residues 1262–1458) and POLA2 were subcloned into a modified, polycistronic pCDFDuet-1 expression vector (Novagen) encoding an HRV 3C-cleavable hexahistidine (POLA1_C_) fusion tag. Production of baculovirus and expression of POLA1/POLA2 in *Spodoptera frugiperda* Sf9 cells were performed by the Expression and Microbiology Core at UC Berkeley. PRIM1/PRIM2, POLA1_cat_, and POLA1_C_/POLA2 were expressed in *Escherichia coli* Rosetta 2(DE3) cells. Cultures were grown at 37°C in LB medium supplemented with 34 mg/L chloramphenicol, 100 mg/L ampicillin (PRIM1/PRIM2), 30 mg/L kanamycin (PRIM1/PRIM2 and POLA1_cat_), and 70 mg/L streptomycin (POLA1_C_/POLA2). Upon reaching mid-log phase, cultures were further supplemented with 100 mg/L ammonium iron(III) citrate (PRIM1/PRIM2) and 5 μM ZnSO_4_ (PRIM1/PRIM2 and POLA1_C_/POLA2) and cooled to 16°C. Protein overexpression was then induced by addition of IPTG to a final concentration of 0.5 mM. After overnight incubation, cells were harvested by centrifugation. Cell pellets were stored at −80°C. *Acanthamoeba castellanii* TLP2 was expressed as previously described ^68–70^. Protein constructs are depicted in **Supplemental Fig. S33**.

### Protein purification

Cell pellets were thawed in lukewarm water and resuspended in buffer L [50 mM sodium phosphate pH 7.5, 1 M NaCl, 5% (v/v) glycerol, 1 mM MgCl_2_, 10 mM imidazole, and cOmplete EDTA-free protease inhibitor cocktail (Roche)]. Both insect cells and bacterial cells were lysed by Dounce homogenization followed by gentle sonication. The two lysates were then combined and cleared by centrifugation prior to application onto a Ni-NTA Agarose column (Qiagen) equilibrated in buffer L lacking protease inhibitor cocktail. The column was washed with 15 column volumes (cv) of buffer N1 [50 mM sodium phosphate pH 7.5, 1 M NaCl, 5% (v/v) glycerol, 1 mM MgCl_2_, and 20 mM imidazole], followed by 5 cv of buffer N2 [50 mM HEPES•NaOH pH 7.5, 300 mM NaCl, 5% (v/v) glycerol, 1 mM MgCl_2_, and 20 mM imidazole]. Protein was eluted by linearly transitioning over 15 cv to buffer N3 [50 mM HEPES•NaOH pH 7.5, 300 mM NaCl, 5% (v/v) glycerol, 1 mM MgCl_2_, and 500 mM imidazole]. Fractions containing polα–primase were pooled and supplemented with 1 mM TCEP before fusion tags were removed by overnight cleavage at 4°C. The following day, protein was diluted three-fold in buffer H1 [50 mM HEPES•NaOH pH 7.5, 5% (v/v) glycerol, 1 mM MgCl_2_, and 1 mM TCEP] and passed through a 0.22-μm filter before being applied to a heparin Sepharose column (Cytiva) equilibrated in a 9:1 mixture of buffer H1 and buffer H2 [50 mM HEPES•NaOH pH 7.5, 1 M NaCl, 5% (v/v) glycerol, 1 mM MgCl_2_, and 1 mM TCEP]. The column was then washed with 5 cv of the same 9:1 mixture before protein was eluted by linearly transitioning over 10 cv to only buffer H2. Fractions containing polα–primase were pooled and again applied to a Ni-NTA Agarose column equilibrated in buffer L lacking protease inhibitor cocktail. Protein was eluted in 10 cv of the same buffer, and fractions containing polα–primase were pooled and supplemented with 1 mM TCEP. Protein was then concentrated by diafiltration, cleared by centrifugation, and applied to a Superose 6 Increase column (Cytiva) equilibrated in buffer S [20 mM HEPES•NaOH pH 7.5, 150 mM NaCl, 5 mM MgCl_2_, and 1 mM TCEP]. Protein was eluted in 1.5 cv of the same buffer, and fractions containing polα–primase were pooled and concentrated by diafiltration. Aliquots were used immediately or frozen in liquid nitrogen and stored at −80°C. POLA1_cat_ and ΔPOLA1_cat_ were purified in the same manner as polα–primase. AcaTLP2 was purified as previously described ^68–70^.

### Oligonucleotide synthesis and purification

RNA, DNA, and chimeric RNA-DNA oligonucleotides were purchased from Integrated DNA Technologies and resuspended in buffer A (10 mM Mes•NaOH pH 6.5 and 40 mM NaCl). To generate 5′-triphosphorylated primers, 500 μM 5′-monophosphorylated primer and 550 μM template were annealed by heating to 85°C and then slowly cooling to 22°C, producing a double-stranded substrate for the tRNA-modifying enzyme AcaTLP2 ^70,71^. AcaTLP2 reaction mixtures were prepared with 50 μM enzyme, 100 μM primer/template, 150 μM ATP, 1 mM GTP, 25 mM Hepes•NaOH pH 7.5, 125 mM NaCl, 10 mM MgCl_2_, and 3 mM DTT and incubated at 22°C. After 1 h, an equal volume of quenching/loading buffer containing 80% (v/v) formamide, 100 mM EDTA pH 8.0, and 220 μM competitor oligonucleotide was added. Samples were then heated to 85°C and slowly cooled to 22°C to denature the primer and the template. Oligonucleotides were separated by denaturing urea-polyacrylamide gel electrophoresis (PAGE) as previously described ^72^, except electrophoresis was performed at 30 W for 9 h. For each reaction, the band corresponding to the triphosphorylated primer was located by UV-shadowing and cut from the gel. The primer was then extracted into 24 mL of 300 mM NaOAc pH 5.5 using a crush-and-soak procedure in which the sample was placed at −80°C until frozen, thawed in a water bath at 42°C, and then rocked overnight at 4°C. The following day, the supernatant was decanted and overnight extraction was repeated using a second 24-mL volume of 300 mM NaOAc pH 5.5. The two volumes were combined, passed through a 0.22-μm filter, and split into four 12-mL aliquots. 36 mL of cold 100% ethanol was then added to each aliquot. Samples were thoroughly mixed, incubated at −80°C for 2 h, warmed to −20°C, and centrifuged at 12,000×g and 4°C for 1 h. After decanting the supernatant, pellets were air-dried for ∼15 min and resuspended in 1 mL of 300 mM NaOAc pH 5.5, using the same 1 mL of buffer for all pellets. The sample was then split into two 0.5-mL aliquots, and 1.5 mL of cold 100% ethanol was added to each. Samples were again thoroughly mixed, incubated at −80°C for 2 h, warmed to −20°C, and centrifuged at 12,000×g and 4°C for 1 h. After decanting the supernatant, pellets were air-dried for ∼15 min and then gently rinsed with 200 μL of cold 95% (v/v) ethanol. The ethanol was removed and the pellet was air-dried for an additional ∼15 min. The triphosphorylated primer was then resuspended in 100 μL of buffer A, using the same 100 μL of buffer for both pellets, and stored at −80°C. Oligonucleotide sequences are provided in **Supplemental Table S2**.

### Cryo-EM sample preparation and data collection

Polα–primase was concentrated to ≥ 3 μM by diafiltration. To generate RNA-primed templates, 110 μM 5′-triphosphorylated RNA primer and 100 μM DNA template were annealed by heating to 85°C and then slowly cooling to 4°C. Substrate complexes were prepared by incubating 3 μM polα–primase, 3.6 μM primer/template, and 1 mM dGTP on ice for 30 min. Using a Vitrobot Mark IV (FEI), 2.5 μL of sample was applied to glow-discharged, holey-carbon R1.2/1.3 200-mesh copper grids (Quantifoil) at 4°C and 100% humidity. Grids were immediately blotted with filter paper (Ted Pella) and flash-frozen in liquid ethane. Grids prepared without substrate or with DT substrate were loaded into a Titan Krios G4 electron microscope (Thermo Scientific) operated at 300 kV for automated data collection using SerialEM ^73^. Movies of dose-fractionated frames were acquired with a K3 BioQuantum direct electron detector (Gatan) functioning in counting mode. Each of the datasets was collected from a single grid in a single imaging session. Grids prepared with DI or DE substrates were loaded into a Talos Arctica electron microscope (Thermo Scientific) operated at 200 kV for automated data collection using Leginon ^74^. Movies of dose-fractionated frames were acquired with a K2 Summit direct electron detector (Gatan) functioning in counting mode. Each of the datasets was collected in three to four imaging sessions, using a different grid for each session. Additional details for each of the datasets are provided in **Supplemental Tables S1, S3, S4, and S6**.

### Cryo-EM data processing

For pre-processing of all datasets, motion-correction of raw movies was performed using MotionCor2 ^75^ and patch CTF estimation was performed for motion-corrected micrographs using cryoSPARC ^76^. All initial processing steps were also performed in cryoSPARC. Particles from each dataset were picked by a combination of blob picking, template picking, and neural-network picking using Topaz ^77^. Templates for template picking were selected from the highest-resolution 2D class averages resulting from blob picking, while Topaz was trained from a small set of manually picked particles. Particles were extracted with a box size of either 400 (substrate-free and DT datasets) or 360 (DI and DE datasets) pixels and then downsampled to 120 pixels (pixel size of 2.73 Å) by Fourier truncation. 2D classification was first performed separately for particles picked by each method. Selected particles were then combined and duplicate and triplicate particles were removed before final 2D classification was performed. For the DT dataset, particles were sorted manually before initial 3D reconstructions were generated by ab initio reconstruction. For all other datasets, initial 3D reconstructions were generated by multiclass ab initio reconstruction followed by heterogeneous refinement. Satisfactory 3D reconstructions could only be obtained for the AI complex (substrate-free and DT datasets), the DNA synthesis assembly (DI, DE, and DT datasets), and the tetramer core (all datasets). Because of extreme configurational and conformational heterogeneity, satisfactory 3D reconstructions could not be obtained for the full polα–primase complex bound to a primer/template substrate. For reconstructions of the tetramer core, conformational heterogeneity was reduced through additional iterations of multiclass ab initio reconstruction and heterogeneous refinement. For all 3D reconstructions, homogeneous refinement was initially performed using downsampled particles and then repeated after particles were recentered and re-extracted either without downsampling or with reduced downsampling. For the AI complex from the substrate-free dataset and the DNA synthesis assembly from the DI, DE, and DT datasets, all subsequent processing steps were performed in RELION-3.1 ^78^. Particle alignments were first improved by focused 3D refinement, before heterogeneity was reduced through focused, alignment-free 3D classification (following particle subtraction, as necessary). CTF refinement was also performed to correct for per-particle defocus differences, as well as for imperfect microscope alignment during data collection. Using refined CTF parameters, focused 3D refinement was repeated for each final set of particles. To improve high-resolution features in the reconstructions, despite local quality differences, maps were sharpened using DeepEMhancer ^79^. Data-processing workflows are outlined in **Supplemental Figs. S5, S10, S12, and S27**.

A modified workflow was employed to obtain reconstructions of the full polα–primase complex bound to primer/template substrates. Particles corresponding to configurations DI-1, DI-2, DE-1, and DE-2 were manually selected by examination of 2D class averages and downsampled to 64 pixels (pixel size of 5.12 Å) by Fourier truncation. Ab initio reconstruction and non-uniform refinement were then performed to generate low-resolution 3D reconstructions. For particles initially assigned to configuration DI-2, multiclass ab initio reconstruction and heterogeneous refinement revealed a possible third configuration of the DI complex. All processing steps in the modified workflow were performed in cryoSPARC. Modified data-processing workflows are outlined in **Supplemental Figs. S19 and S20**.

### Model building and refinement

Homology models of *X. laevis* polα–primase subunits were generated from corresponding *Homo sapiens* X-ray crystal structures using SWISS-MODEL ^80^. For the AI complex, POLA1, POLA2_C_, and PRIM2 homology models (from PDB accession 5EXR) were docked as rigid bodies into the high-resolution partial reconstruction. Following manual modification, atomic coordinates and isotropic temperature factors were refined against the cryo-EM density for all non-hydrogen atoms using Ramachandran, secondary structure, and rotamer restraints. Hydrogen atoms were placed in riding positions and were not refined against the density. Both the refined model and a PRIM1 homology model (from PDB accession 4BPU) were then docked as rigid bodies into the complete reconstruction. After manual adjustment, atomic coordinates and isotropic temperature factors were refined as described above, except reference restraints were applied based on the *X. laevis* PRIM2/POLA1/POLA2_C_ structure and the *H. sapiens* PRIM1 structure. For the DI subcomplex, POLA1_cat_ and PRIM2_C_ homology models (from PDB accessions 4QCL and 5F0Q) were docked as rigid bodies into the corresponding reconstruction. Following manual modification of the protein and de novo addition of the substrate, atomic coordinates and isotropic temperature factors were refined as described above, except base pairing and base stacking restraints were added for the oligonucleotide duplex. For the DE subcomplex, the refined POLA1_cat_ model from the DI subcomplex was docked as a rigid body into the high-resolution partial reconstruction. After manual adjustment of the protein and de novo addition of the substrate, atomic coordinates and isotropic temperature factors were refined as described above for the DI subcomplex. The refined models of POLA1_cat_ from the partial DE structure and PRIM2_C_ from the DI structure were then docked as rigid bodies into the complete reconstruction. The oligonucleotide substrate from the partial DE subcomplex and the 3′-overhang and first base pair from the *H. sapiens* PRIM2_C_ crystal structure (PDB accession 5F0Q) were also docked as rigid bodies. Following manual modification of the protein, including adjustment of the template-binding residues in PRIM2_C_ to match those in the *H. sapiens* crystal structure, and de novo addition of the intervening region of the substrate, atomic coordinates and isotropic temperature factors were refined as described above for the DI subcomplex, except reference restraints were applied based on the *X. laevis* POLA1_cat_ and PRIM2_C_ structures. For the DT subcomplex, the refined POLA1_cat_ model from the DE subcomplex was docked as a rigid body into the high-resolution partial reconstruction. After manual adjustment of the protein and de novo addition of the substrate, atomic coordinates and isotropic temperature factors were refined as described above for the DI subcomplex. The refined models of POLA1_cat_ from the partial DT structure and PRIM2_C_ from the complete DE structure were then docked as rigid bodies into the reconstruction of DTS-1. The oligonucleotide substrate from the partial DT subcomplex and the 3′-overhang and first nine base pairs from the DE subcomplex were also docked as rigid bodies. Following manual modification of the protein and de novo addition of the intervening region of the substrate, atomic coordinates and isotropic temperature factors were refined as described above for the complete DE subcomplex. Similarly, the refined model of POLA1_cat_ from the partial DT structure, the oligonucleotide substrate from the partial DT structure, and the 3′-overhang and first nine base pairs from the complete DE structure were docked as rigid bodies into the reconstruction of DTS-2. Following manual modification of the protein and de novo addition of the intervening region of the substrate, atomic coordinates and isotropic temperature factors were refined as described above for the complete DE subcomplex. For all structures, manual protein modification and de novo substrate addition were performed in Coot ^81^, while docking and refinement were performed in PHENIX ^82^. If possible, rotamers were determined from the cryo-EM density. In instances in which the correct conformation of the side chain was ambiguous, rotamers were selected to match those in higher-resolution cryo-EM structures from this study or those in previously published crystal structures. In instances in which both the correct conformation was ambiguous and an appropriate reference structure was unavailable, rotamers were manually selected to avoid steric clashes, and if possible, form favorable electrostatic interactions. Final models were validated using MolProbity ^83^.

Structures of the complete polα–primase complex in configurations DI-1, DI-2, DE-1, and DE-2 and of the TC subcomplex in configurations 1, 2, and 3 were generated from rigid bodies. For all structures, the TC subcomplex was reconstituted from three rigid bodies taken from the AI complex, while the DI and DE subcomplexes were each maintained as a single rigid body. Docking into low-resolution reconstructions was performed in ChimeraX ^84^.

Refinement and validation statistics are provided in **Supplemental Figs. S7, S14, S15, S19, S20, and S29 and Supplemental Tables S1, S3, S4, and S6**. Figures were prepared in PyMOL (https://www.pymol.org), UCSF Chimera ^85^, or UCSF ChimeraX.

### Primer elongation

To generate an RNA-primed template for elongation assays, 110 μM 5′- triphosphorylated RNA primer and 100 μM DNA template were annealed by heating to 85°C and then slowly cooling to 4°C. Reaction mixtures were then prepared with 50 nM POLA1_cat_, ΔPOLA1_cat_, POLA1_cat_/ΔPOLA1_cat_, or polα–primase; 500 nM primer/template; 0.1 μM [α-^32^P]-ATP (3,000 Ci/mmol; PerkinElmer) or [α-^32^P]-dATP (3,000 Ci/mmol; PerkinElmer); 9.9 μM ATP or dATP; 200 μM GTP, CTP, and UTP; 20 μM dGTP, dCTP, and dTTP; 20 mM HEPES•NaOH pH 7.5; 150 mM NaCl; 5 mM MgCl_2_; and 1 mM TCEP and incubated at 22°C. 8-μL aliquots were periodically withdrawn, quenched with 2 μL of 100 mM EDTA pH 8.0, and incubated at 65°C for 15 min. To denature the primer and the template, 10 μL of loading buffer [80% (w/v) formamide, 10 mM EDTA pH 8.0, 1 mg/mL bromophenol blue, 1 mg/mL xylene cyanol, and 1.5 μM competitor oligonucleotide] was added and samples were incubated at 85°C for 10 min and then slowly cooled to 22°C. Elongation products were separated by denaturing urea-PAGE as previously described ^72^, except 18% polyacrylamide (19:1 acrylamide/bisacrylamide) gels were run at 20 W for 7 h and visualized by phosphorimaging using an Amersham Typhoon biomolecular imager (Cytiva). Experiments were performed in triplicate.

### Primer/template binding

Binding reactions were performed in 20 mM HEPES•NaOH pH 7.5, 150 mM NaCl, 5 mM MgCl_2_, 1 mM TCEP, 20 μM dGTP, and 0.05% (v/v) Tween-20 with 10 nM Cy5-labeled primer/template and 0.15–5,120 nM POLA1_cat_. Immediately after mixing, samples were centrifuged at 20,000×g and 4°C for 5 min, incubated on ice for 10 min, and then incubated at 20°C for 10 min. Microscale thermophoresis data were collected using a Monolith NT.115 instrument (NanoTemper). All experiments were performed using the red fluorescence laser (λ_excitation_ = 600–640 nm; λ_emission_ = 660–720 nm) at 80% power and the infrared laser (λ = 1480 nm) at medium power. Temperature-related intensity changes were monitored at 20°C with a pre-IR phase of 3 s, followed by an IR-on phase of 20 s, and a post-IR phase of 1 s. Data quality was assessed by monitoring fluorescence quenching and photobleaching, as well as protein aggregation and adsorption. Experiments were performed in triplicate. For each protein-substrate binding titration, normalized fluorescence values [F_norm_ = (F_1 (IR-hot)_)/(F_0 (IR-cold)_)] were tracked as parts per thousand and converted to ΔF_norm_ [F_norm (n)_−F_norm (initial)_]. ΔF_norm_ values for individual isotherms were normalized to generate the fraction of bound protein (*f*_B_). Dissociation constants (*K*_d_) were derived from fits to the data using a one-site binding model [*f*_B_ = *x*/(*K*_d_+*x*), where *x* is the total concentration of POLA1_cat_].

### Data availability

Structures were deposited in the Protein Data Bank under accession codes 8G99 (AI complex, partial), 8G9F (AI complex, complete), 8G9L (DI subcomplex), 8G9N (DE subcomplex, partial), 8G9O (DE subcomplex, complete), 8UCU (DT subcomplex, partial), 8UCV (DT subcomplex 1), and 8UCW (DT subcomplex 2). Maps were deposited in the Electron Microscopy Data Bank under accession codes EMD-29862 (AI complex, partial), EMD-29864 (AI complex, complete), EMD-29871 (DI subcomplex), EMD-29872 (DE subcomplex, partial), EMD-29862 (DE subcomplex, complete), EMD-42140 (DT subcomplex, partial), EMD-42141 (DT subcomplex 1, complete), EMD-42142 (DT subcomplex 2, complete), EMD-29888 (TC subcomplex, conformation 1), EMD-29889 (TC subcomplex, conformation 2), and EMD-29891 (TC subcomplex, conformation 3).

## Acknowledgments

This work was funded by National Institutes of Health grants R35GM136401 (B.F.E.), R35GM118089 (W.J.C.), R01GM087543 (J.E.J.), and PO1CA92584. We thank Miaw-Sheue Tsai and members of the Expression and Microbiology Core at UC Berkeley for baculovirus production and insect cell protein expression, and members of the Eichman and Chazin research groups for helpful discussions. Cryo-EM data were collected at the Center for Structural Biology Cryo-EM Facility at Vanderbilt University and the Life Sciences Institute Cryo-EM Facility at the University of Michigan. The content is solely the responsibility of the authors and does not necessarily represent the official views of the National Institutes of Health.

## Author contributions

B.F.E. conceived of the study; B.F.E. and W.J.C. oversaw the study; E.A.M., L.E.S., N.P.B., and B.F.E. designed experiments; J.E.J. provided reagents; E.A.M., L.E.S., C.L.D., and N.P.B. collected data; E.A.M., L.E.S., N.P.B., M.D.O., W.J.C., and B.F.E. analyzed data; E.A.M., L.E.S., N.P.B., W.J.C., and B.F.E. wrote the manuscript.

