## Supplementary Information for "A mechanistic model of primer synthesis from catalytic structures of DNA polymerase α–primase"

```
XlPOLA1
XlPOLA1     1  .........MSDSGSFAASRSRREKTEKSGRKEALERLKRAKAGEKVKYEVEQVSSIYEEVDEAEYSKLVR.DRQDDDWI
HsPOLA1     1  MAPVHGDDSLSDSGSFVSSRARREKKSKKGRQEALERLKKAKAGEKYKYEVEDFTGVYEEVDEEQYSKLVQ.ARQDDDWI
ScPOLA1     1  .....................MSSKSEKLEKLRKLQAARNGTSIDDYEGDESDGDRIYDEIDEKEYRARKRQELLHDDFV
TtPOLA1     1  ...........................MSDKLTRLERLNKEVKKQNKLKQHSKNNRFDDDMDIEAYEDDEQ.........

XlPOLA1
XlPOLA1    71  VDDDGTGYVE.....DGREIFDDDLEDNALADSGKGAKGAP..KDKTNVKKSSVSKPNNIKSMFMASAVKKTTDKAVDLS
HsPOLA1    80  VDDDGIGYVE.....DGREIFDDDLEDDALDADEKGKDGKARNKDKRNVKKLAVTKPNNIKSMFIACAGKKTADKAVDLS
ScPOLA1    60  VDDDGVGYVDRGVEEDWREVDNSSSDEDTGNLASKDSKRKKNIKREKDHQITDMLRTQHSKSTLLAHAKKSQKKSIPIDN
TtPOLA1    45  ........IE......EDDFIDDTQED......KKYKKYREIEDEFDQEIEEEEELN.......KK...KKTKNTILNYT

XlPOLA1
XlPOLA1   144  KDDLLGDL...........LQDLKSQAVPITPPPVITLKKKKLAGSPLNPFSVPPTAPKVL.PTSVKRLPAVTKPGHPAA
HsPOLA1   155  KDGLLGDI...........LQDLNTETPQITPPPVMILKKKRSIGASPNPFSVHTAT.............AVPSGKIASP
ScPOLA1   140  FDDILGEFESGEVEKPNILLPSKLRENLNSSPTSEFKSSIKRVNGNDESSHDAGISKKVKIDPDSSTDKYLEIESSPLKL
TtPOLA1    96  NTTAVTNNK..........KKAISKQIPDIDIEEIMKLTERKKKIEQEEAQLLQEEQ.............ELLEQEKREE

XlPOLA1
XlPOLA1   212  QSKASVPRQIKKEPKAELISSAVGPLKVEAQVKEEDSGMVEFDDGDFDEPME.EDVEITPVDSSTIKTQAQSIKCVKEEN
HsPOLA1   211  VSRKEPPLTPVPLKRAEFAGDDV...QVESTEEEQESGAMEFEDGDFDEPMEVEEVDLEPMAAKAWDKESEPAEEVKQEA
ScPOLA1   220  QSRKLRYANDVQDLLDDVENSPVVATKRQNVLQDTLLANPPSAQSLADEEDD.EDSDEDIILKRRTMRSVTTTRRVNIDS
TtPOLA1   153  EEKKRISQEAKSILREDCASSKT....ANSSKGKVDQNILNAINRDFSDDSN....TVDSISEFQKLKSLAQKANLANES

                                                                       β1              β2
XlPOLA1
XlPOLA1   291  .IKEEKSSFITSATLNESCWDQIDEAEP..MTTEIQVDSSHLPLVTGADGSQVFRFYWLDAYEDQYSQPGVVYLFGKVWI
HsPOLA1   288  DSGKGTVSYLGSFLPDVSCWDIDQEGDSSFSVQEVQVDSSHLPLVKGADEEQVFHFYWLDAYEDQYNQPGVVFLFGKVWI
ScPOLA1   299  RSNPSTSPFVTAPGTPIGIKGLTPSKSLQSNTDVATLAVNVKKEDVVDPETDTFQMFWLDYCEVNN....TLILFGKVKL
TtPOLA1   225  .LKQSKVSNTEINLTNLSISQVKKINDYK...........N........EDGSVDAYLYDYFYDAQVKPDKIYAFAKVQN

                  β3         β4      β5      β6        α1      α2         β7
XlPOLA1
XlPOLA1   368  ESADAYVSCCVSVKNIERTVYLLPRENRVQLSTGKDTGAPVSMMHVYQEFNEAVAEKYKIMKFKSKKVDKDYAFEIPDVP
HsPOLA1   368  ESAETHVSCCVMVKNIERTLYFLPREMKIDLNTGKETGTPISMKDVYEEFDEKIATKYKIMKFKSKPVEKNYAFEIPDVP
ScPOLA1   375  K.DDNCVSAMVQINGLCRELFFLPREG..KTPT..D.........IHEEIIPLLMDKYGLDNIRAKPQKMKYSFELPDIP
TtPOLA1   285  KQTNAFDTCVIQIDTIIRNLFFYPSSDTVTEQQIKN.........EIAELLKKEQTSRKNVEFLGAFVDKNYAFELPIPR

                 β8        TT          TT        β9     TT     α3         β10              β11
XlPOLA1
XlPOLA1   448  ASSEYLEVRYSADSPQ.....LPQDLKGETFSHVFGTNTSSLELFLLSRKIKGPSWLEIKSPQLSSQP.MSWCKVEAVVT
HsPOLA1   448  EKSEYLEVKYSAEMPQ.....LPQDLKGETFSHVFGTNTSSLELFLMNRKIKGPCWLEVKSPQLLNQP.VSWCKVEAMAL
ScPOLA1   441  SESDYLKVLLPYQTPKSSRDTIPSDLSSDTFYHVFGGNSNIFESFVIQNRIMGPCWLDIKGADFNSIRNASHCAVEVSVD
TtPOLA1   356  GKSRWYQVVMSYEYEV.....ISPDTKGQYFSYCVGSTYSALETFLITKKITGPSWVRFQNVKDTTSC.ITNRKLEFRVD

                η1   β12             β13          TTT        β14                   β15
XlPOLA1
XlPOLA1   522  RPDQ...VSVVKDLAPPPVVLSLSMKT......VQNAKTHQNEIVAIAALVHHTFPLDKAPPQPPFQTHFCVLSKL.ND
HsPOLA1   522  KPDL...VNVIKDVSPPPLVVMAFSMKT......MQNAKNHQNEIIAMAALVHHSFALDKAAPKPPFQSHFCVVSKP.KD
ScPOLA1   521  KPQN...ITPTTTKTMPNLRCLSLSIQT......LMNPKENKQEIVSITLSAYRNISLDSPIPENIKPDDLCTLVRPPQS
TtPOLA1   430  YTNQSNIQVLQKQLPTPPLSVVCISLKTSQQIVLSQKKKEYKKEIFNLNMKYHEGINIDNSNKDE..LNQFKSISFITHI

                     α4           β16          α5               β17          α6            η2
XlPOLA1
XlPOLA1   592  CIFPYDYNEAVKQKN.ANIEIALTERTLLGFFLAKIHKIDPDVIVGHDIYGFDLEVLLQRINSCKVPFWSKIGRLRRSVM
HsPOLA1   592  CIFPYAFKEVIEKKN.VKVEVAATERTLLGFFLAKVHKIDPDIIVGHNIYGFELEVLLQRINVCKAPHWSKIGRLKRSNM
ScPOLA1   592  TSFPLGLAALAKQKLPGRVRLFNNEKAMLSCFCAMLKVEDPDVIIGHRLQNVYLDVLAHRMHDLNIPTFSSIGRRLRRTW
TtPOLA1   508  DPTKKQDSITKKGTLPETTKFCLNELNLLEQFLVHFNEIDPDIVVAHDLYSTVFEIILTRIREKGIRKWNLLSKLINIGS

                              η3      β18   α7                α8               α9    η4
XlPOLA1
XlPOLA1   671  PKLGGRSG.....FAERNAACGRIICDIEISAKELIR..CKSYHLSELVHQILKAERVVIPPENIRNAYNDSV.HLLYML
HsPOLA1   671  PKLGGRSG.....FGERNATCGRMICDVEISAKELIR..CKSYHLSELVQQILKTERVVIPMENIQNMYSESS.QLLYLL
ScPOLA1   672  PEKFGRGNSNMNHFFISDICSGRLICDIANEMGQSLTPKCQSWDLSEMYQVTCEKEHKPLDIDYQNPQYQNDVNSMTMAL
TtPOLA1   588  SDIPKYGSS...TFKTKMAMKGRLLVDTLLSSQEFVN..CVEYTLEALAQKLFKIEIPRIDAKAYQQKFATYK.LLNSLV

                    α10                  α11            α12            α13
XlPOLA1
XlPOLA1   743  ENTWIDAKFILQIMCELNVLPLALQITNIAGNVMSRTLMGGRSERNEYLLLHAFTENNFIVPD........KPVFKKMQQ
HsPOLA1   743  EHTWKDAKFILQIMCELNVLPLALQITNIAGNIMSRTLMGGRSERNEFLLLHAFYENNYIVPD........KQIFRKPQQ
ScPOLA1   752  QENITNCMISAEVSYRIQLLTLTKQLTNLAGNAWAQTLGGTRAGRNEYILLHEFSRNGFIVPD........KEGNRSRAQ
TtPOLA1   662  DDTYQDIDYALRIMYHLQIVPLTKQLTSICGNIWMGSLQNQRAERNEMLLLHKFNQLNYVYPDNFKNLPESYKKKHKNAQ

                                                   β19     β20        α14
XlPOLA1
XlPOLA1   815  TTVEDNDDMGTDQNKNK.SRKKAAYAGGLVLEPKVGFYDKFILLLDFNSLYPSIIQEYNICFTTVHREAP..........
HsPOLA1   815  KLGDEDEEIDGDTNKYKKGRKKAAYAGGLVLDPKVGFYDKFILLLDFNSLYPSIIQEFNICFTTVQRVAS..........
ScPOLA1   824  KQRQNEENADAPVNSKK.....AKYQGGLVFEPEKGLHKNYVLVMDFNSLYPSIIQEFNICFTTVDRNK...........
TtPOLA1   742  IRKQYEEDEDQAQGNKNPKKKENKYKGGQVFEPEKGLYNEYIVLLDFNSLYPSIIQEFNVCFTTCVRDPIPLEMQMAPFL
```

**(cont.)**

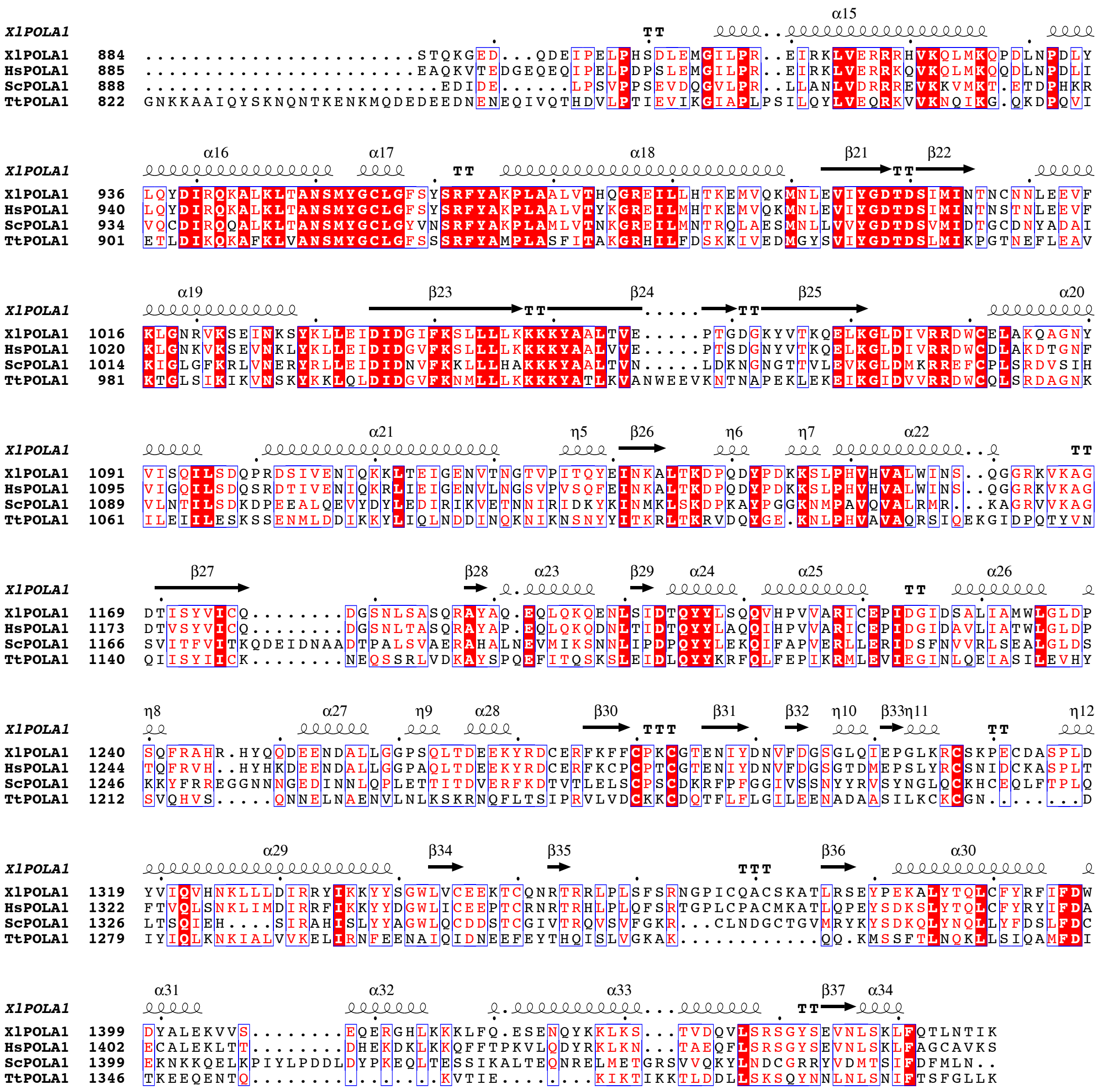

**Supplemental Figure S1. Sequence conservation of POLA1.** The sequence alignment of POLA1 from *Xenopus laevis* (Xl), *Homo sapiens* (Hs), *Saccharomyces cerevisiae* (Sc), and *Tetrahymena thermophila* (Tt) was constructed in Clustal Omega and ESPript. Secondary structural elements above the alignment were taken from the AI complex of *Xenopus* polα–primase.

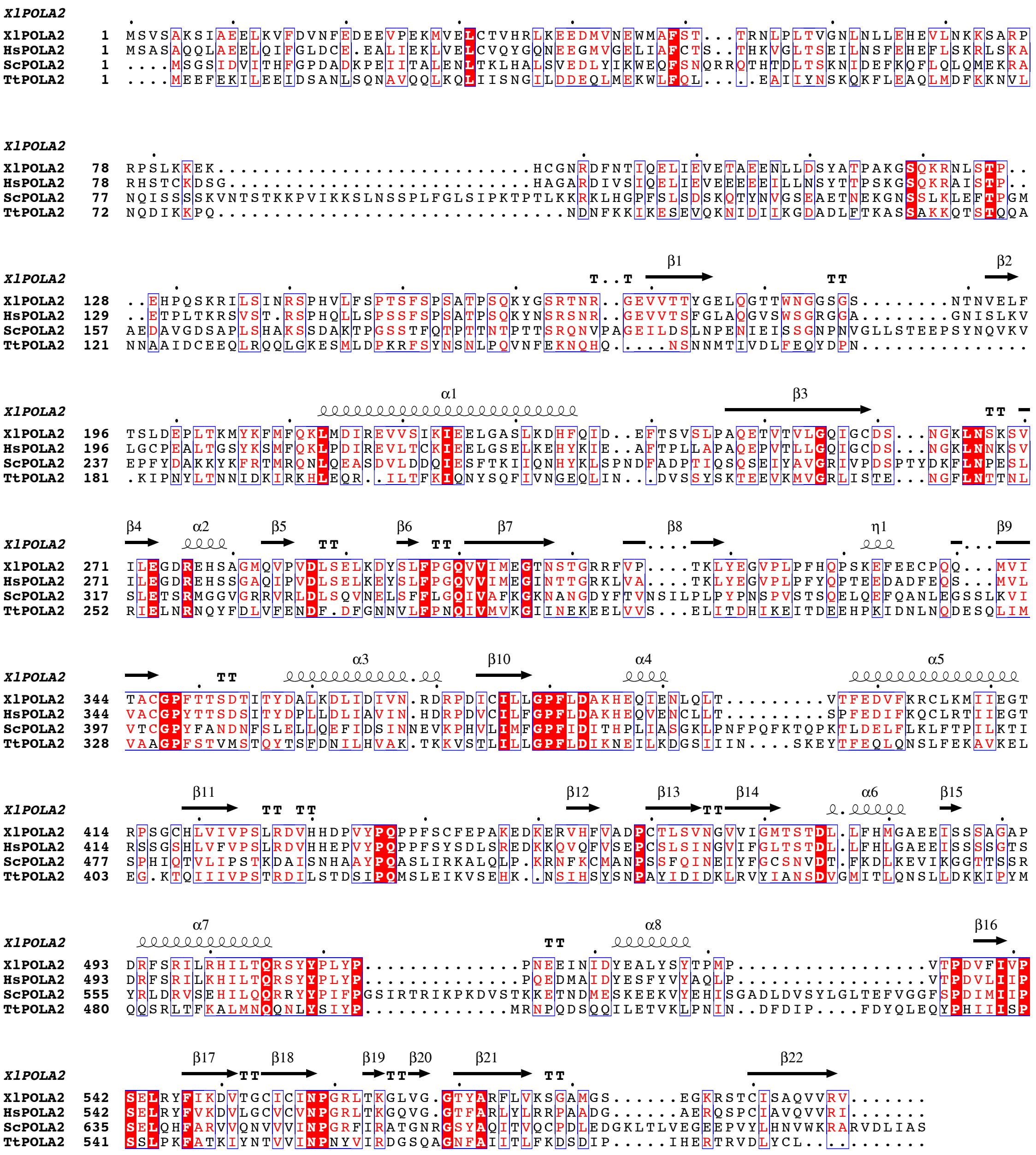


**Supplemental Figure S2. Sequence conservation of POLA2.** The sequence alignment of POLA2 from *Xenopus laevis* (Xl), *Homo sapiens* (Hs), *Saccharomyces cerevisiae* (Sc), and *Tetrahymena thermophila* (Tt) was constructed in Clustal Omega and ESPript. Secondary structural elements above the alignment were taken from the AI complex of *Xenopus* polα–primase.

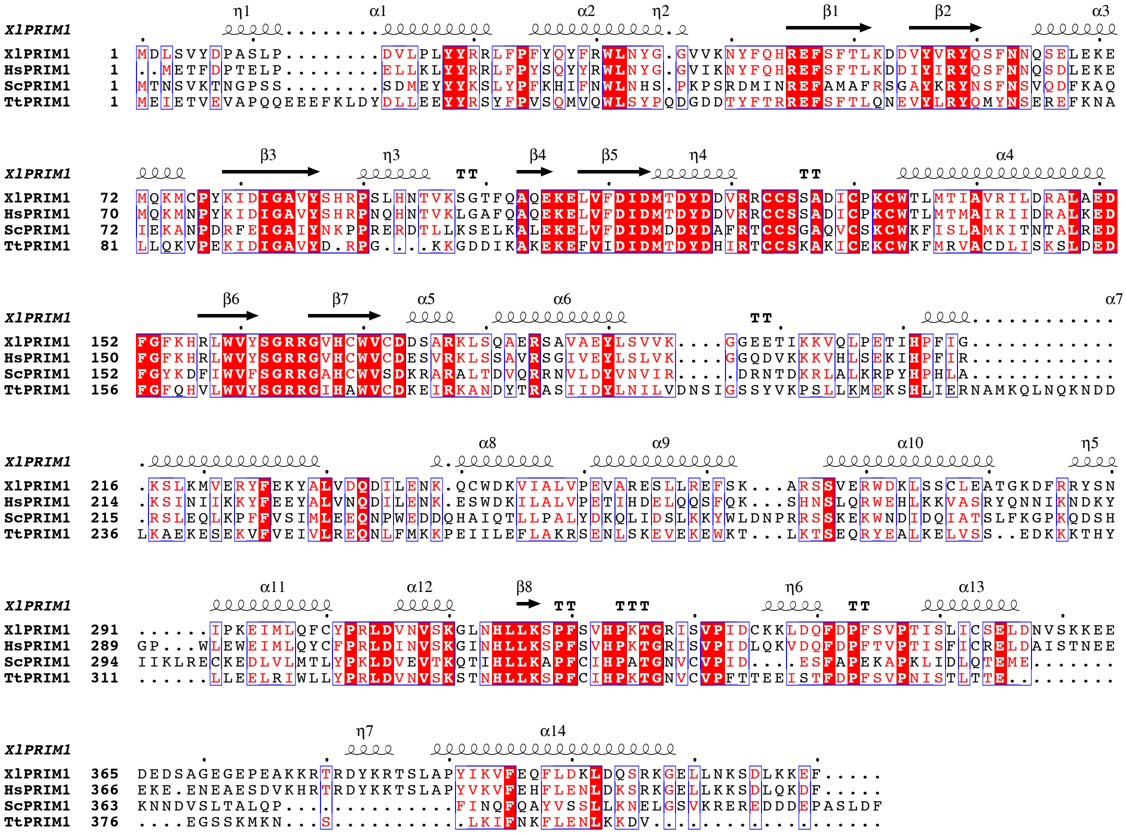


**Supplemental Figure S3. Sequence conservation of PRIM1.** The sequence alignment of PRIM1 from *Xenopus laevis* (Xl), *Homo sapiens* (Hs), *Saccharomyces cerevisiae* (Sc), and *Tetrahymena thermophila* (Tt) was constructed in Clustal Omega and ESPript. Secondary structural elements above the alignment were taken from the AI complex of *Xenopus* polα–primase.

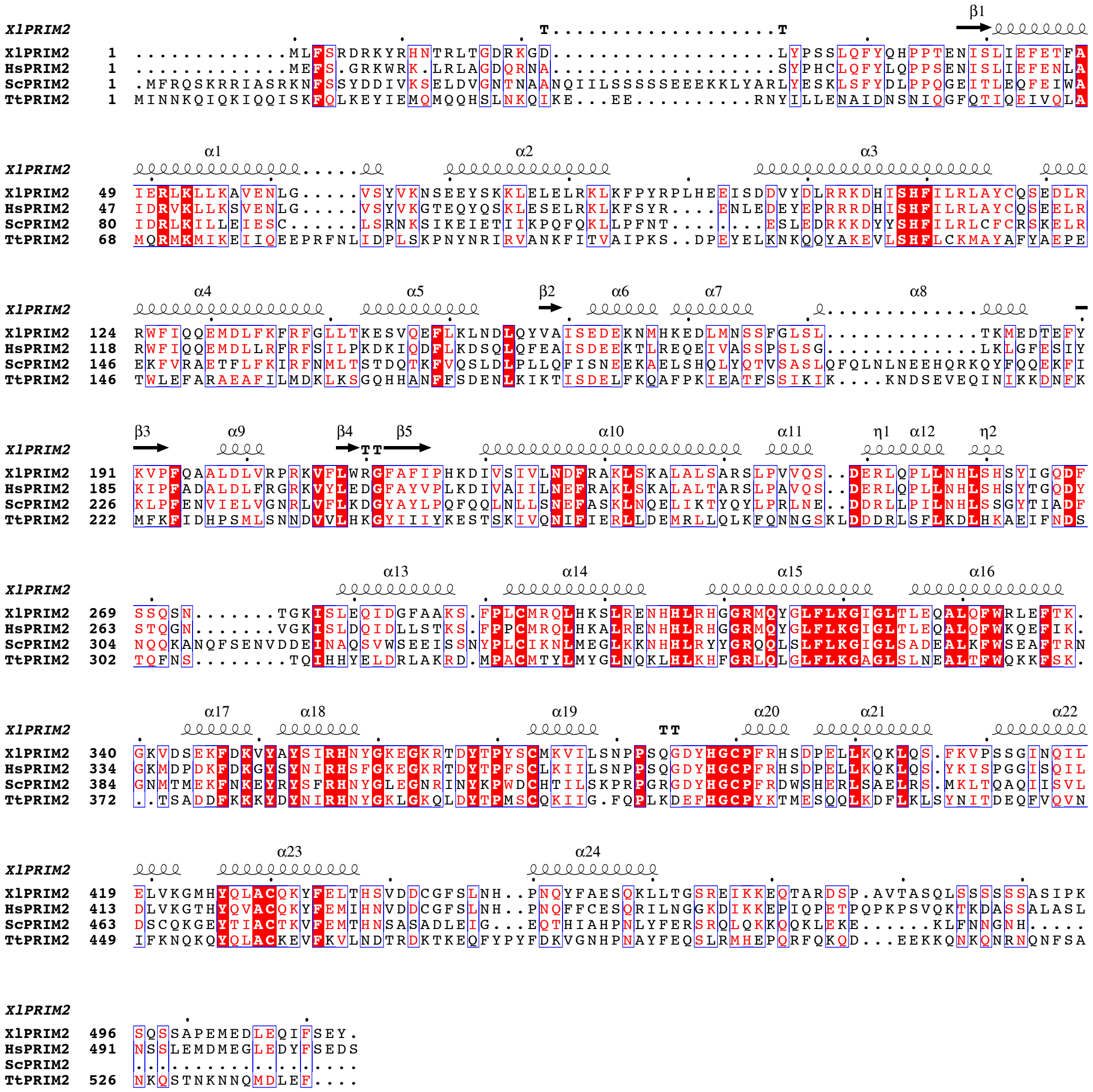

**Supplemental Figure S4. Sequence conservation of PRIM2.** The sequence alignment of PRIM2 from *Xenopus laevis* (Xl), *Homo sapiens* (Hs), *Saccharomyces cerevisiae* (Sc), and *Tetrahymena thermophila* (Tt) was constructed in Clustal Omega and ESPript. Secondary structural elements above the alignment were taken from the AI complex of *Xenopus* polα–primase.

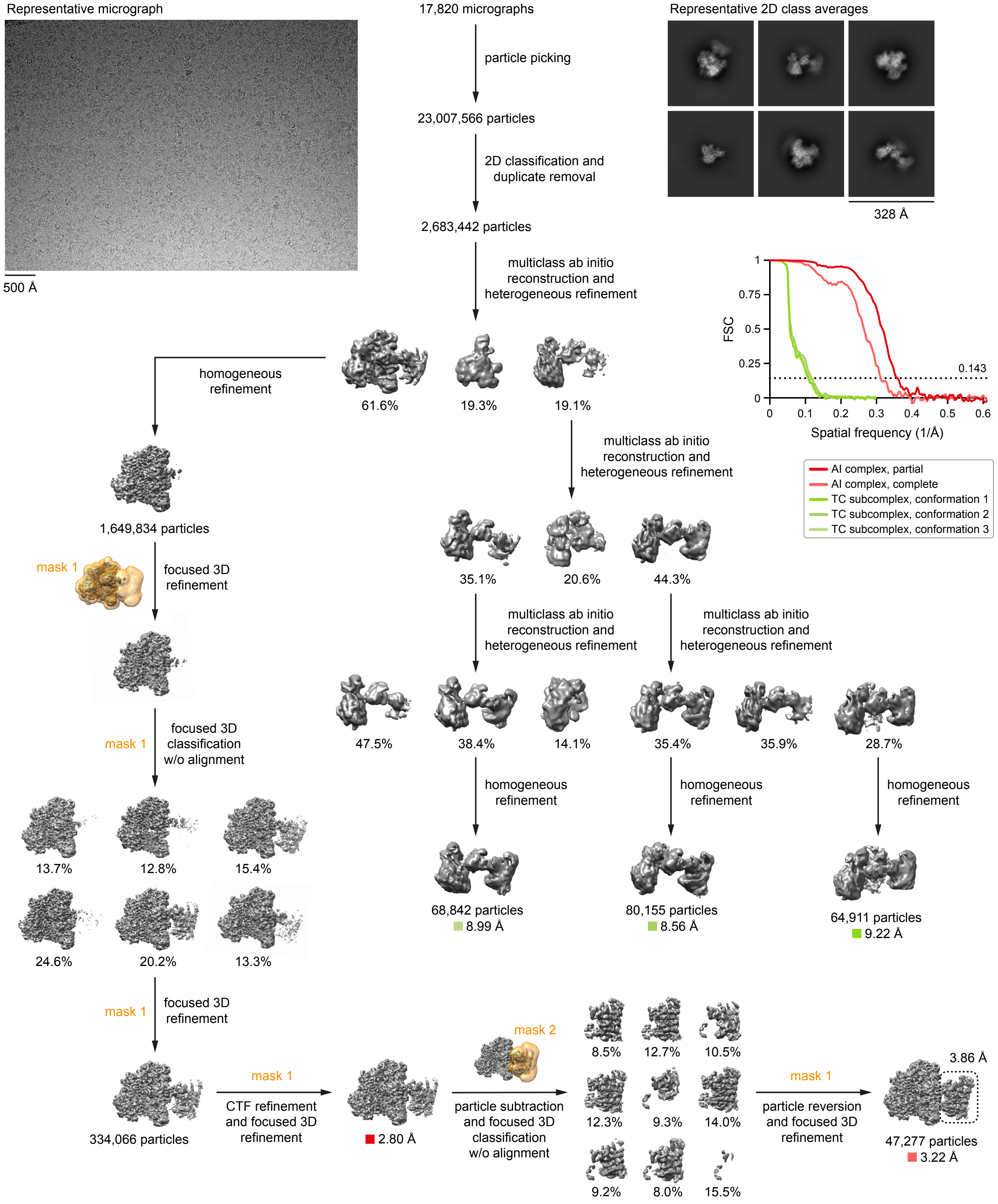


**Supplemental Figure S5. Data-processing workflow for the substrate-free dataset.** All processing steps were performed in cryoSPARC or RELION. 3D reconstructions and masks are colored gray and orange, respectively. Gold-standard Fourier shell correlation (FSC) curves were calculated from independent half-reconstructions. All final 2D class averages are provided in Supplemental Fig. S6. Additional validation metrics are provided in Supplemental Fig. S7 and Supplemental Table S1.

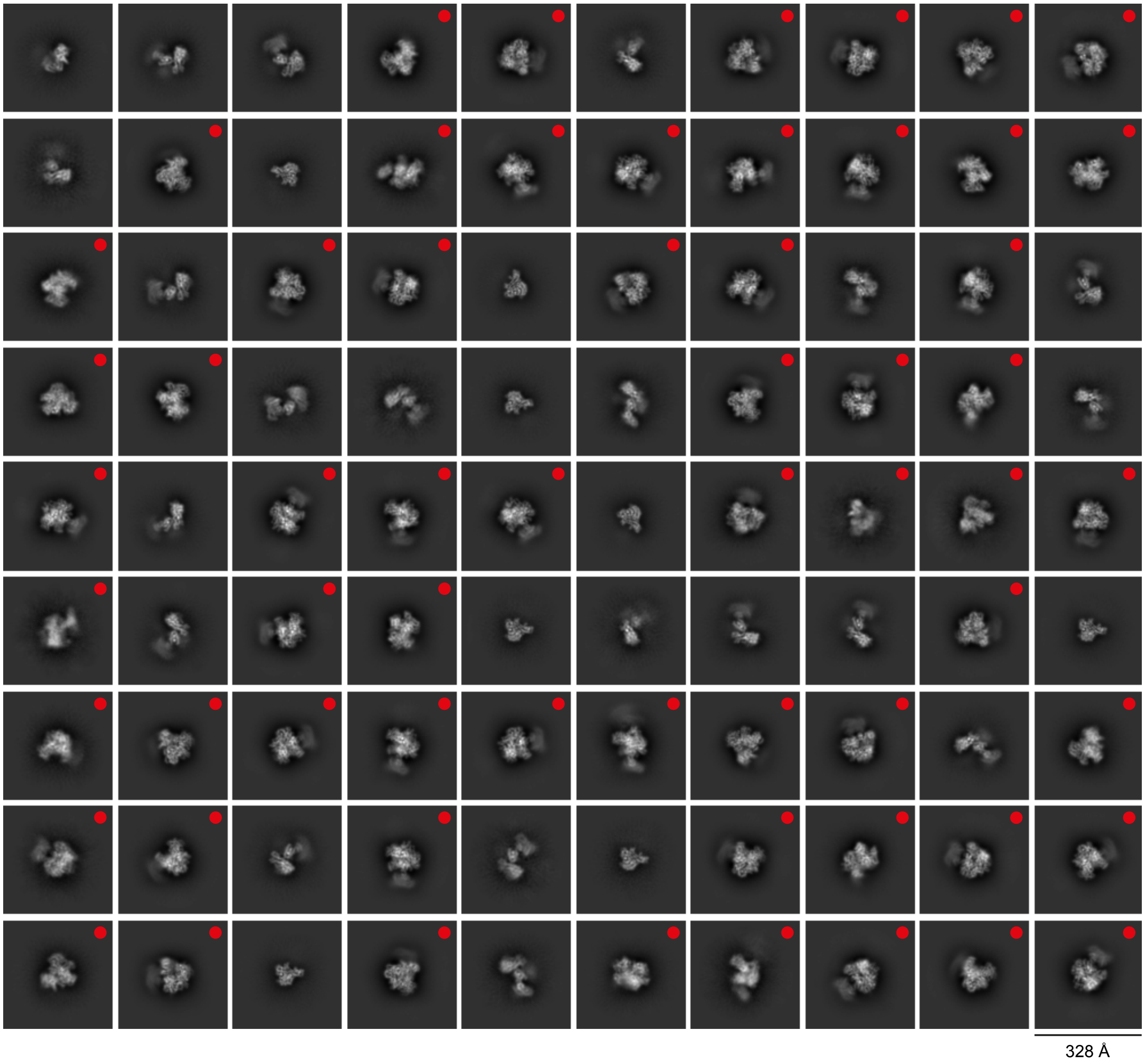


**Supplemental Figure S6. Final 2D class averages from the substrate-free dataset.** 2D class averages show the full AI complex (68% of classes, 69% of particles), the TC subcomplex (31% of classes, 30% of particles), or only POLA1$_{cat}$ (1% of classes, 1% of particles). 2D class averages showing the AI complex are indicated with red dots.

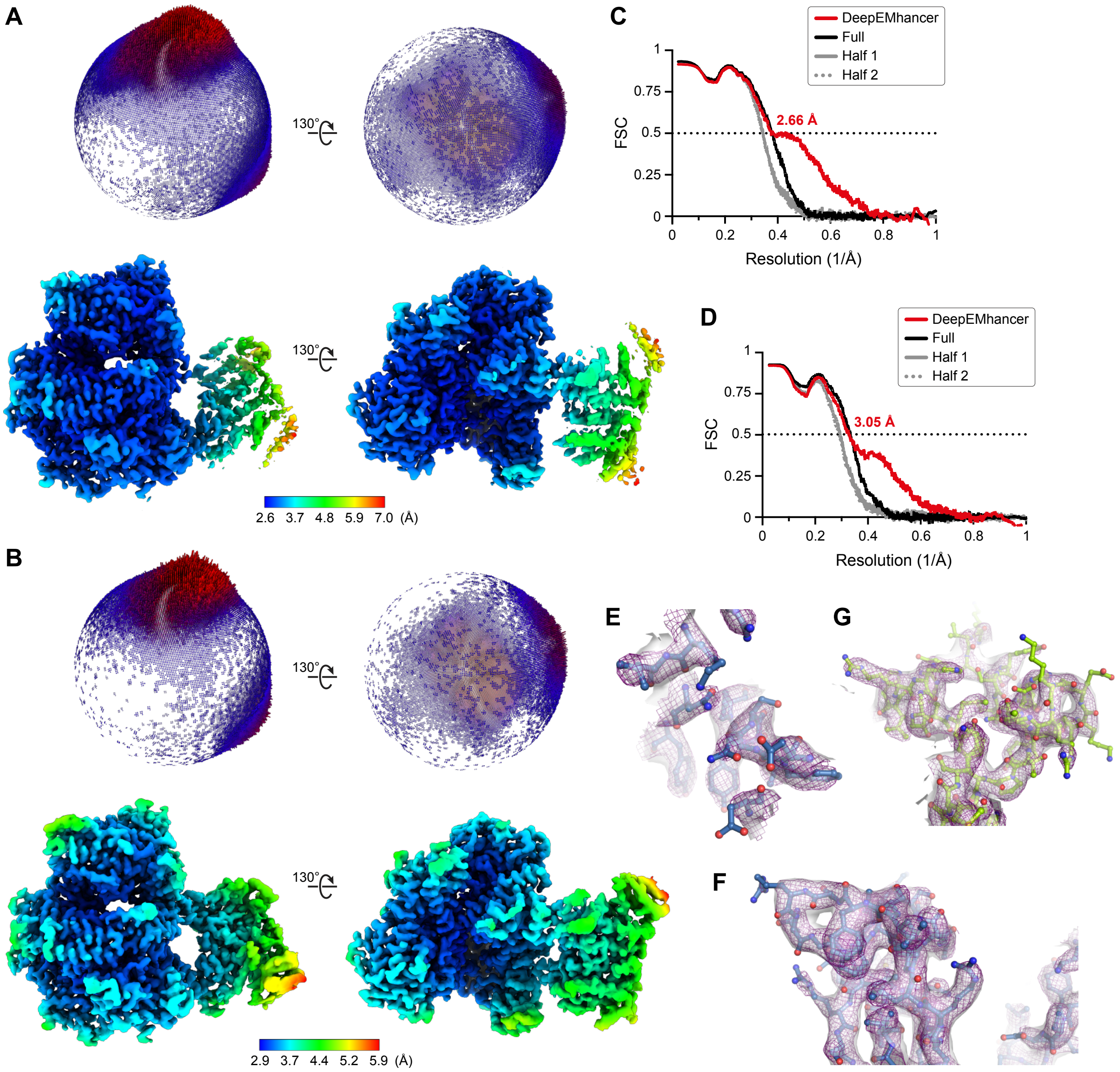


**Supplemental Figure S7. Validation of 3D reconstructions from the substrate-free dataset. A**,**B**. Euler angle distributions and local resolution estimations for the partial (A) and complete (B) reconstructions of the AI complex. **C**,**D**. Model-to-map Fourier shell correlation (FSC) curves for the partial (C) and complete (D) structures. Prior to correlation calculations, structures were refined against the DeepEMhancer-sharpened map, the unsharpened full map, or half map 1. Correlation with half map 2 was calculated relative to the structures refined against half map 1. **E**–**G**. Model-to-map fits for the regions of the AI complex shown in Supplemental Fig. S23. Cryo-EM densities from the DeepEMhancer-sharpened map and the unsharpened full map are shown as a purple mesh and a white surface, respectively.

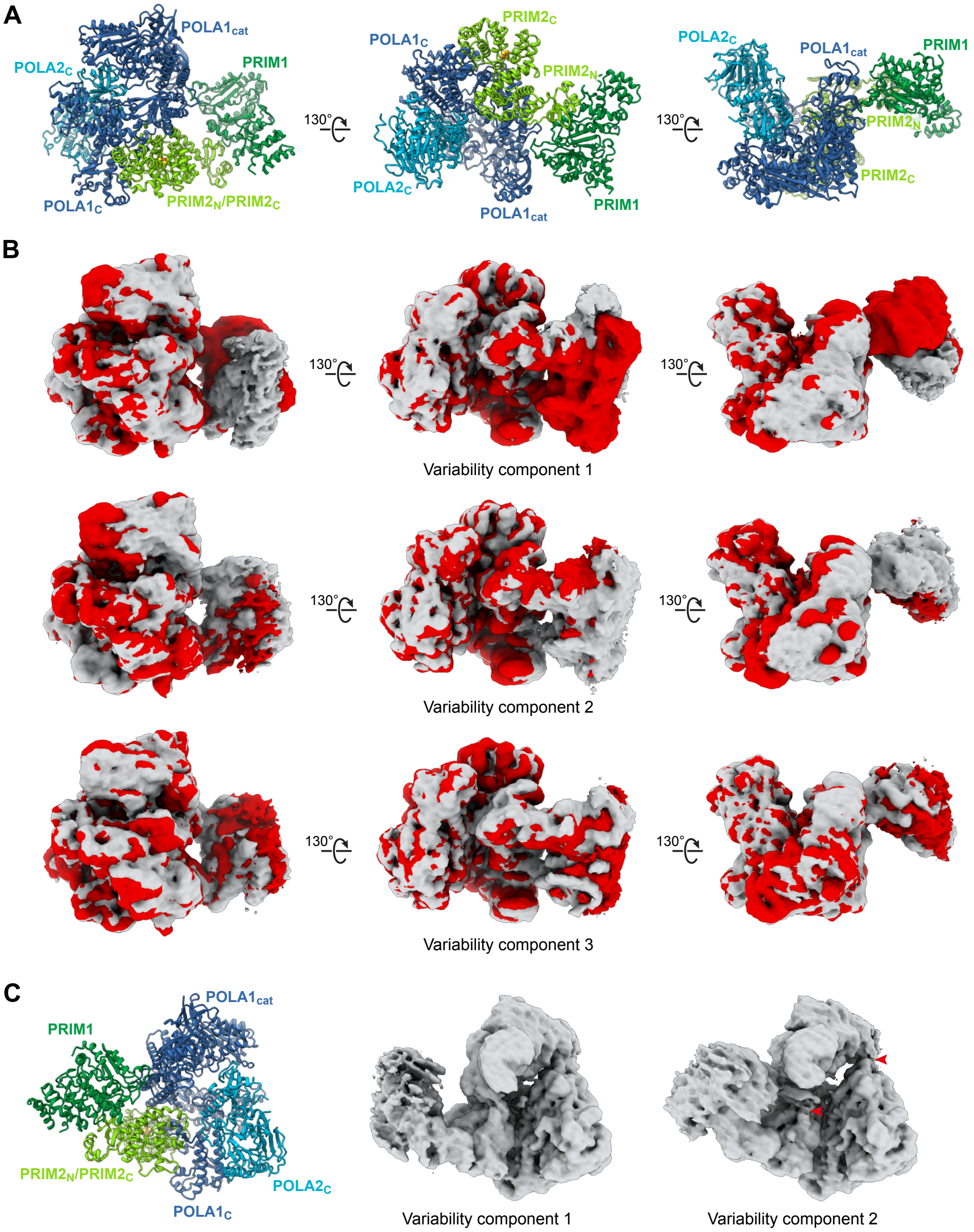


**Supplemental Figure S8. Conformational dynamics in the auto-inhibitory complex. A**. Structural representations of the AI complex oriented as in panel B. **B**. 3D variability analysis of the 1,649,834 particles aligned by focused 3D refinement (Supplemental Fig. S5). Maps were filtered to a resolution of 5 Å. Initial and final frames of variability components 1 (top), 2 (middle), and 3 (bottom) are shown in gray and red, respectively. Variability component 1 resolves a bending motion in PRIM2$_{N}$ that moves PRIM1 away from POLA1$_{cat}$, accompanied by a small bending/twisting motion in POLA1$_{cat}$. Component 2 resolves a larger bending/twisting motion in POLA1$_{cat}$ that pulls the exonuclease domain away from POLA2$_{C}$ and that begins to pull the palm domain away from PRIM2$_{N}$. Component 3 resolves a flexing motion (molecular breathing) that spans the AI complex. In all variability components, interactions involving PRIM2$_{C}$ appear to be unchanged. **C**. Comparison of POLA1$_{cat}$ disengagement in variability components 1 and 2. The points at which POLA1$_{cat}$ begins to separate from POLA2$_{C}$ and PRIM2$_{N}$ are indicated with arrowheads.

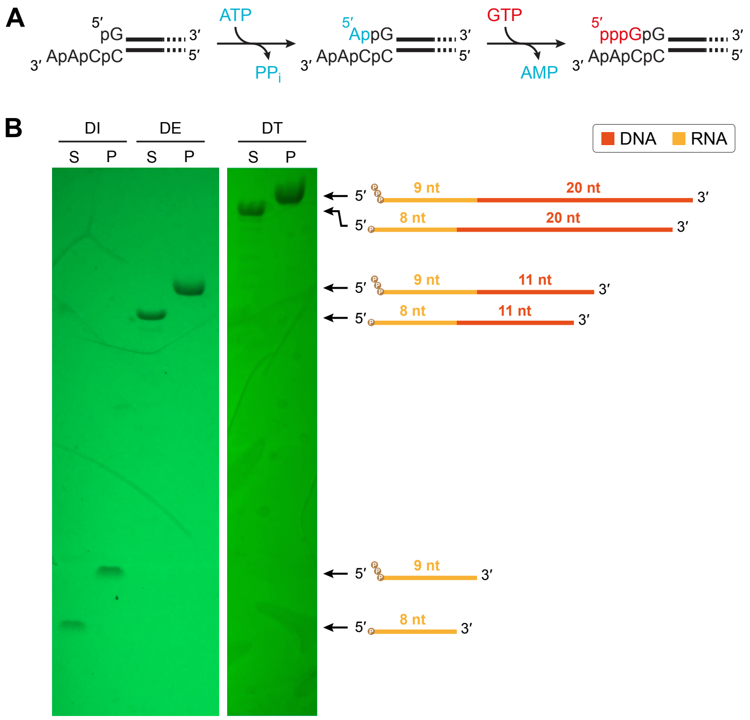


**Supplemental Figure S9. Preparation of 5′-triphosphorylated RNA and RNA-DNA primers. A**. Chemical reaction of AcaTLP2. **B**. Comparison of AcaTLP2 substrates (S) and products (P) for the DI, DE, and DT primers. Oligonucleotides were separated by denaturing urea-PAGE and visualized by UV-shadowing. Phosphate groups are depicted as brown circles.

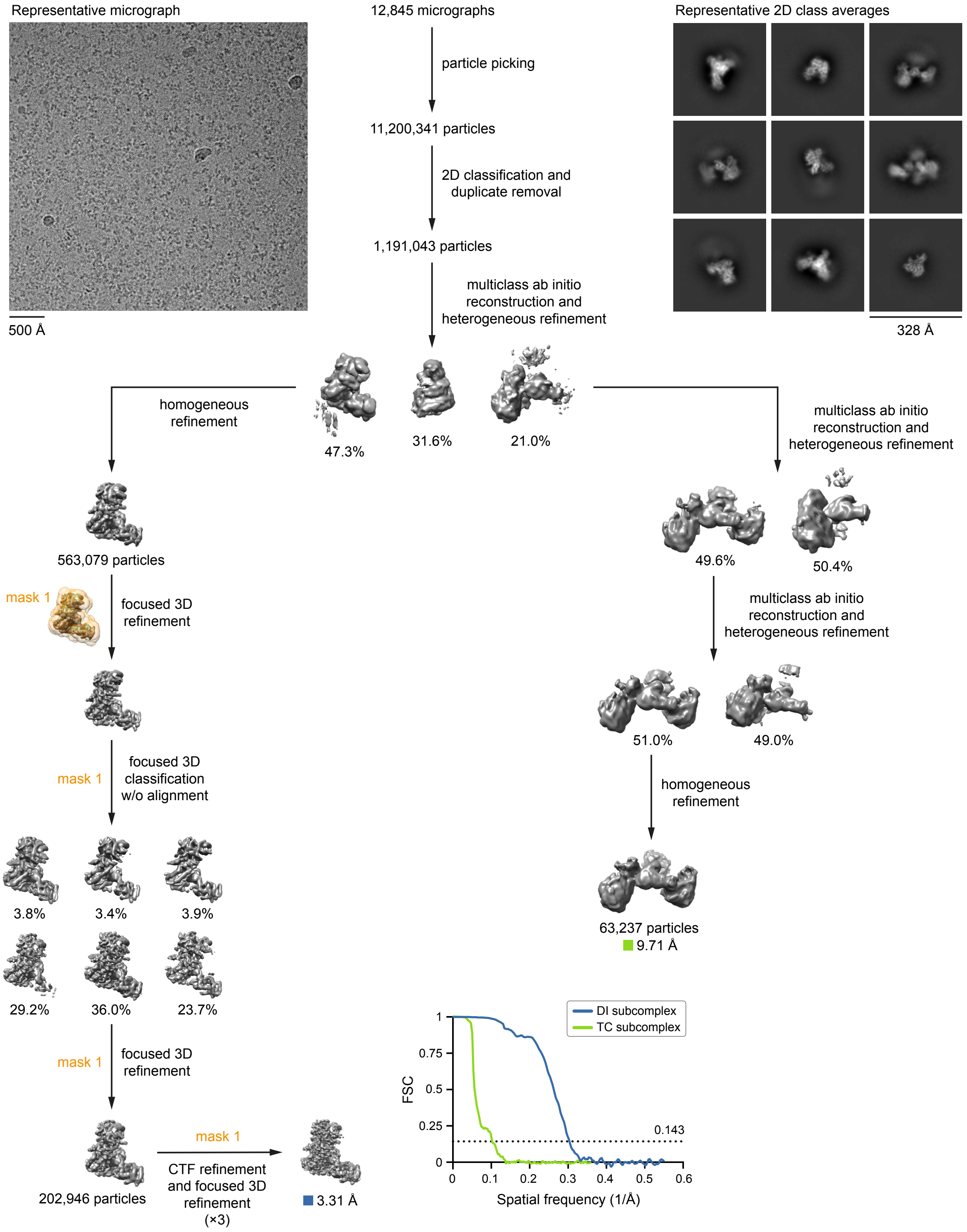


**Supplemental Figure S10. Data-processing workflow for the DNA initiation dataset.** All processing steps were performed in cryoSPARC or RELION. 3D reconstructions and masks are colored gray and orange, respectively. Gold-standard Fourier shell correlation (FSC) curves were calculated from independent half-reconstructions. All final 2D class averages are provided in Supplemental Fig. S11. Additional validation metrics are provided in Supplemental Fig. S14 and Supplemental Table S3.

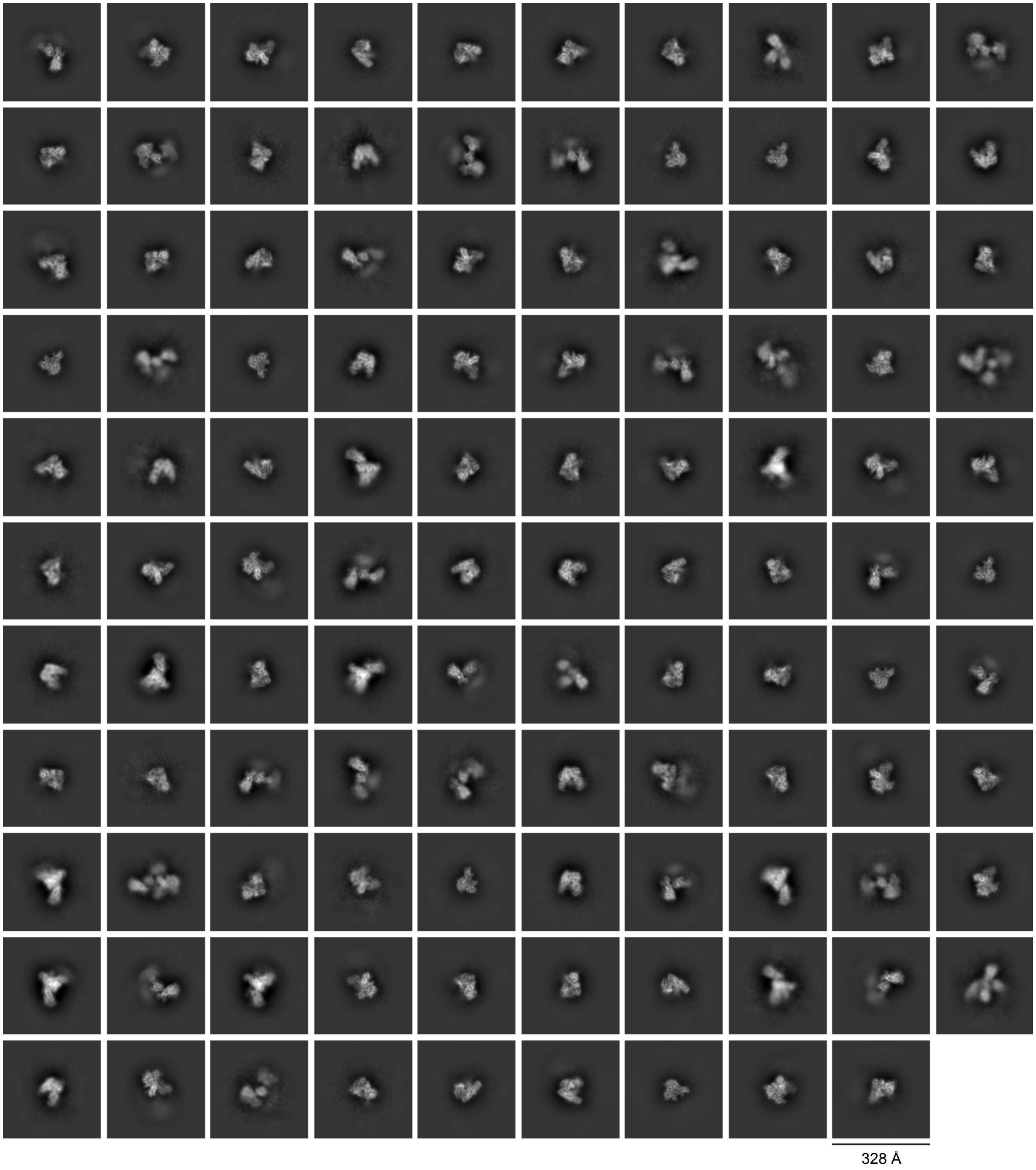


**Supplemental Figure S11. Final 2D class averages from the DNA initiation dataset.** 2D class averages show the DI subcomplex (59% of classes, 67% of particles), the TC subcomplex (26% of classes, 26% of particles), the DI complex in configuration 1 (4% of classes, 1% of particles), the DI complex in configuration 2 (≤ 9% of classes, ≤ 5% of particles), or “other” complexes/configurations (≥ 3% of classes, ≥ 1% of particles). No 2D class averages show the AI complex.

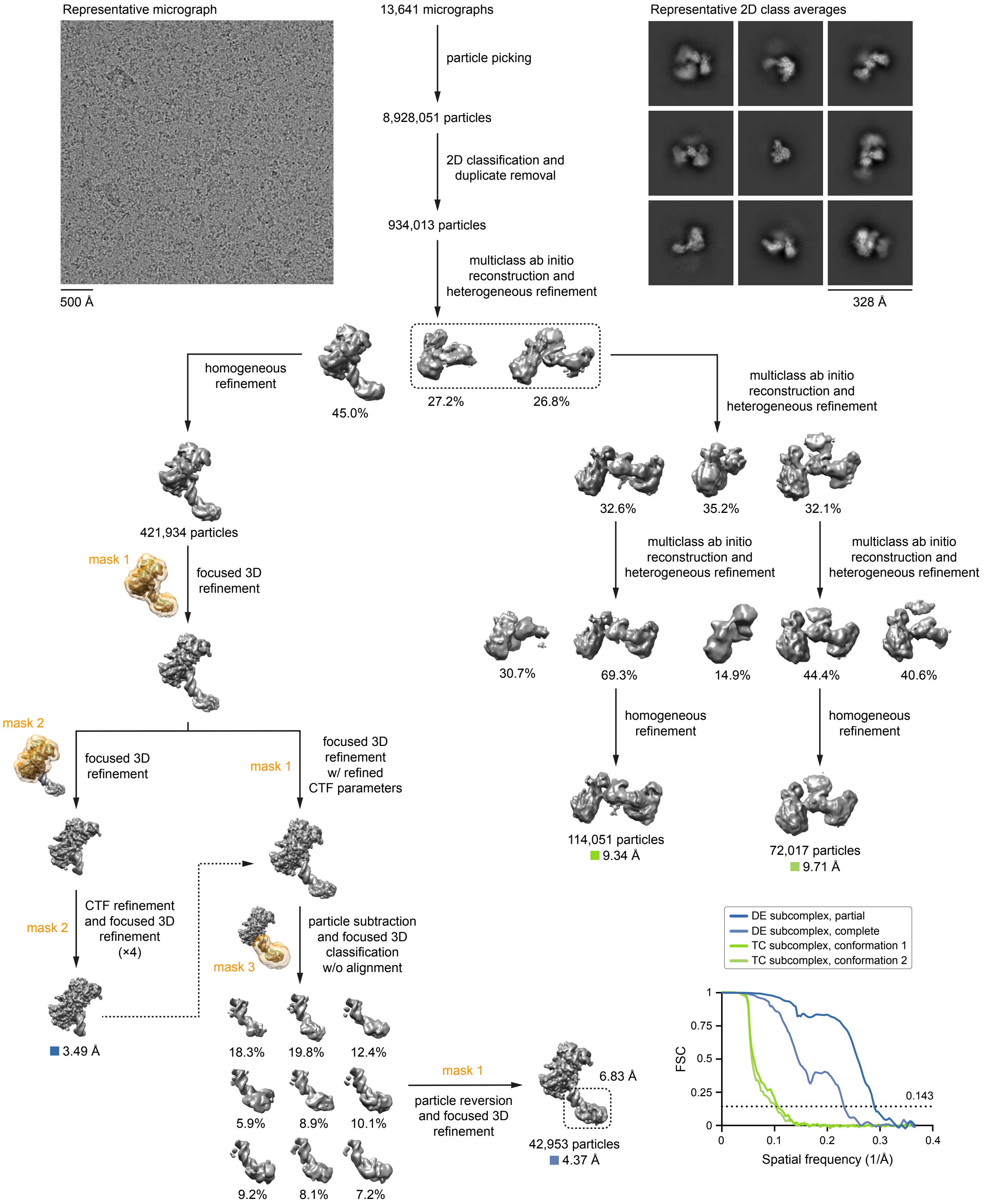


**Supplemental Figure S12. Data-processing workflow for the DNA elongation dataset.** All processing steps were performed in cryoSPARC or RELION. 3D reconstructions and masks are colored gray and orange, respectively. Gold-standard Fourier shell correlation (FSC) curves were calculated from independent half-reconstructions. All final 2D class averages are provided in Supplemental Fig. S13. Additional validation metrics are provided in Supplemental Fig. S15 and Supplemental Table S4.

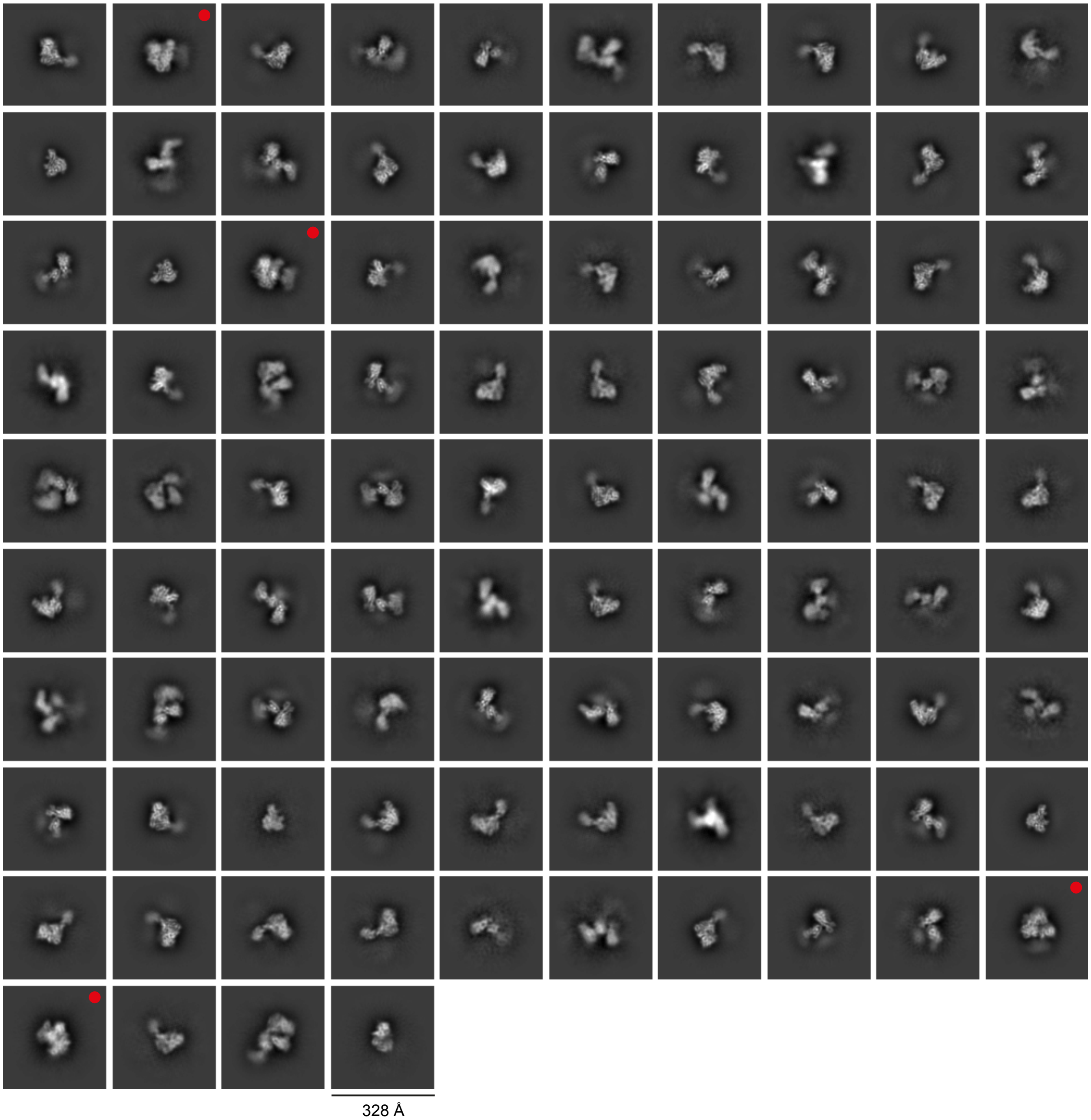


**Supplemental Figure S13. Final 2D class averages from the DNA elongation dataset.** 2D class averages show the DE subcomplex (47% of classes, 48% of particles), the TC subcomplex (43% of classes, 44% of particles), the DE complex in configuration 1 (1% of classes, 1% of particles), the DE complex in configuration 2 (5% of classes, 4% of particles), or the AI complex (4% of classes, 3% of particles). 2D class averages showing the AI complex are indicated with red dots.

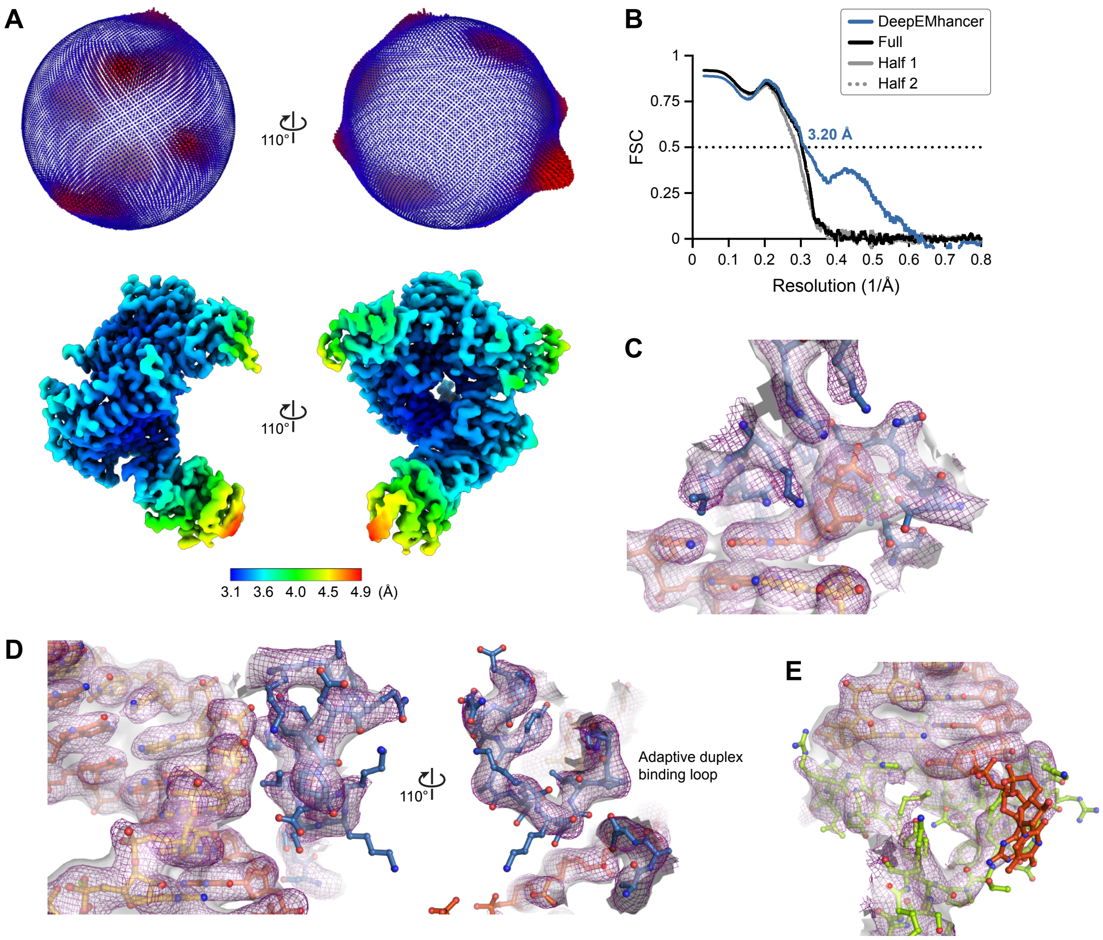


**Supplemental Figure S14. Validation of 3D reconstructions from the DNA initiation dataset. A**. Euler angle distribution and local resolution estimation for the reconstruction of the DI subcomplex. **B**. Model-to-map Fourier shell correlation (FSC) curves for the structure of the DI subcomplex. Prior to correlation calculations, the structure was refined against the DeepEMhancer-sharpened map, the unsharpened full map, or half map 1. Correlation with half map 2 was calculated relative to the structure refined against half map 1. **C**–**E**. Model-to-map fits for the regions of the DI subcomplex shown in Fig. 5. Cryo-EM densities from the DeepEMhancer-sharpened map and the unsharpened full map are shown as a purple mesh and a white surface, respectively. Density for the second and third nucleotides of the 3′-overhang is only apparent at lower contour levels of the unsharpened map.

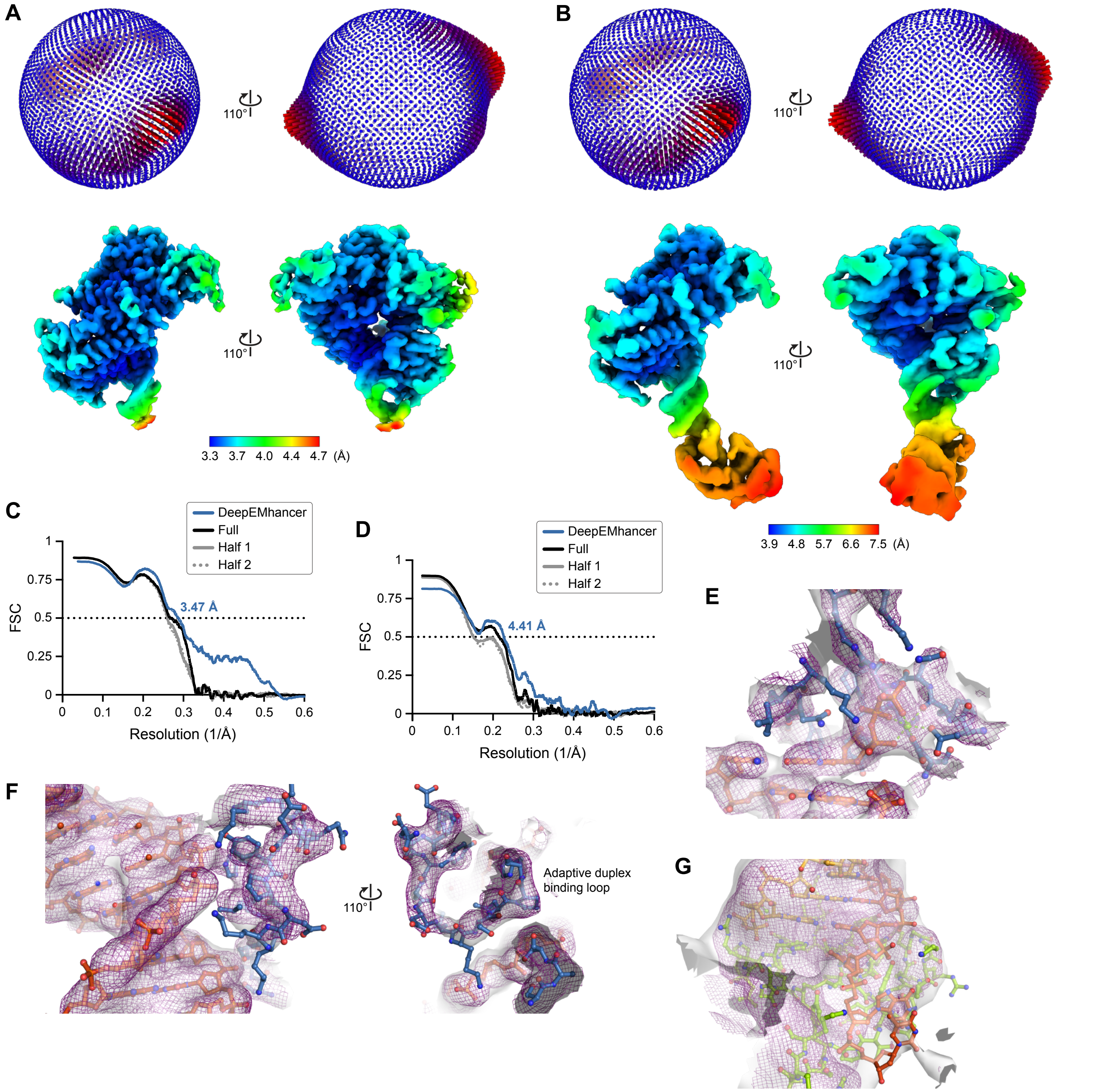


**Supplemental Figure S15. Validation of 3D reconstructions from the DNA elongation dataset. A**,**B**. Euler angle distributions and local resolution estimations for the partial (A) and complete (B) reconstructions of the DE subcomplex. **C**,**D**. Model-to-map Fourier shell correlation (FSC) curves for the partial (C) and complete (D) structures. Prior to correlation calculations, structures were refined against the DeepEMhancer-sharpened map, the unsharpened full map, or half map 1. Correlation with half map 2 was calculated relative to the structures refined against half map 1. **E**–**G**. Model-to-map fits for the regions of the DE subcomplex shown in Fig. 5. Cryo-EM densities from the DeepEMhancer-sharpened map and the unsharpened full map are shown as a purple mesh and a white surface, respectively.

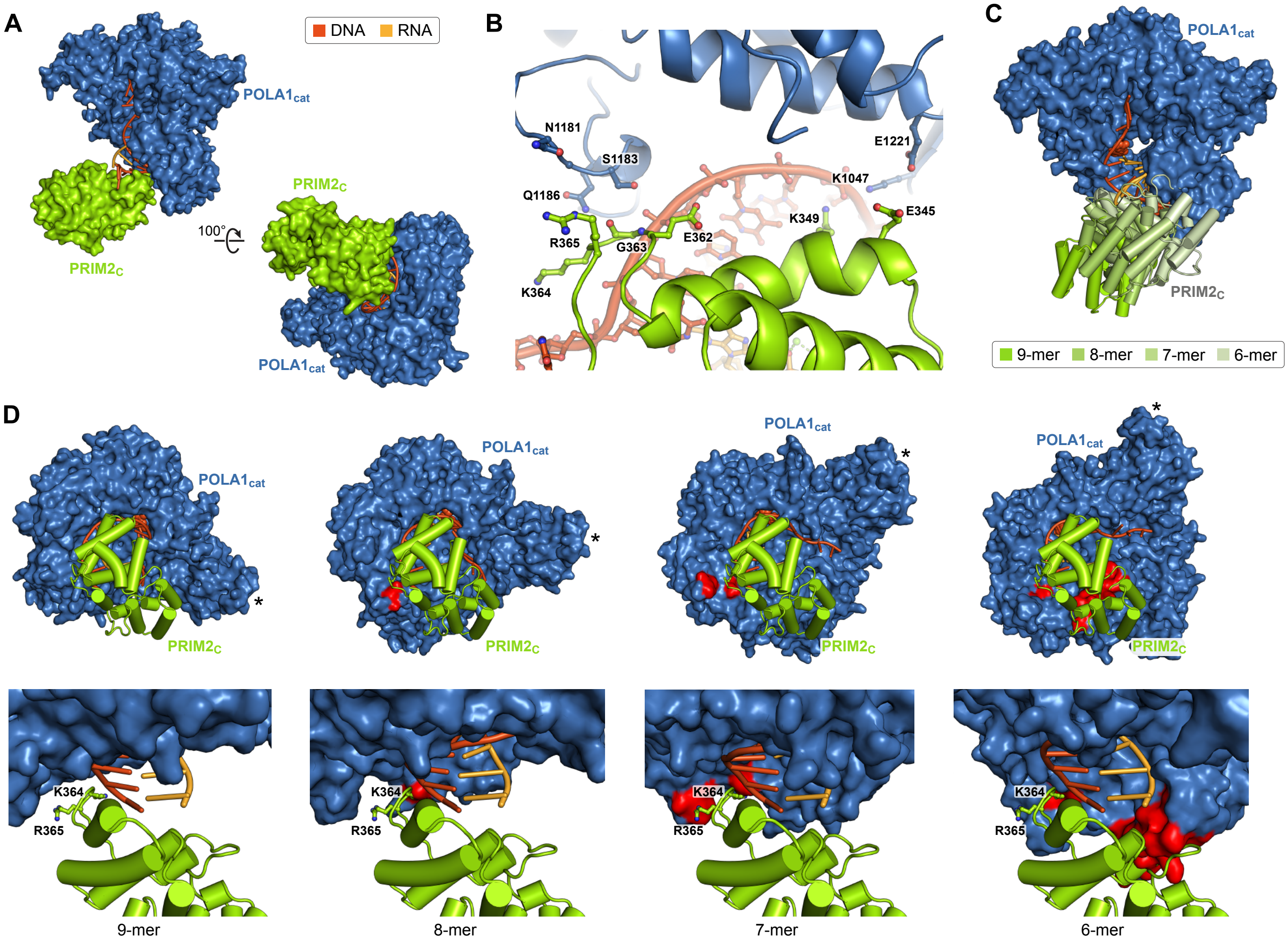


**Supplemental Figure S16. Structural basis for DNA initiation with variable-length RNA primers. A**. DI subcomplex showing surface representations of $POLA1_{cat}$ and $PRIM2_C$ bound to a 9-mer RNA primer, a DNA template, and an incoming dGTP. **B**. Subunit interface between $POLA1_{cat}$ (blue) and $PRIM2_C$ (green) in the DI subcomplex. All residues within 6 Å are shown. Only POLA1 Ser1183 and PRIM2 Gly363 are within van der Waals contact distance. **C**. Overlay of the DI subcomplex and hypothetical DI models containing 8-mer, 7-mer, or 6-mer RNA primers, DNA template, and incoming dGTP, aligned on $POLA1_{cat}$. Models were constructed by serially removing the second base pair from the 5′-end of the primer and then manually repositioning the first base pair, the template 3′-overhang, and $PRIM2_C$ as a rigid body to reform the phosphodiester bonds. As expected for an A-form to intermediate AB-form duplex, reformation of the bonds required the rigid body to be rotated by ~30° and translated by ~3 Å after deletion of each base pair. **D**. Comparison of the subunit interface between $POLA1_{cat}$ and $PRIM2_C$ in the DI subcomplex (left) and the hypothetical DI models containing 8-mer, 7-mer, or 6-mer RNA primers, aligned on $PRIM2_C$. The asterisks denote a common point on $POLA1_{cat}$ in all complexes. Residues in $POLA1_{cat}$ that sterically clash with $PRIM2_C$ are colored red. The extensive steric clashes in the 6-mer model suggest DNA initiation is unlikely with RNA primers ≤ 6 nucleotides in length. However, the minor steric clashes—involving only side chains—in the 7-mer and 8-mer models suggest DNA initiation is possible with RNA primers ≥ 7 nucleotides in length.

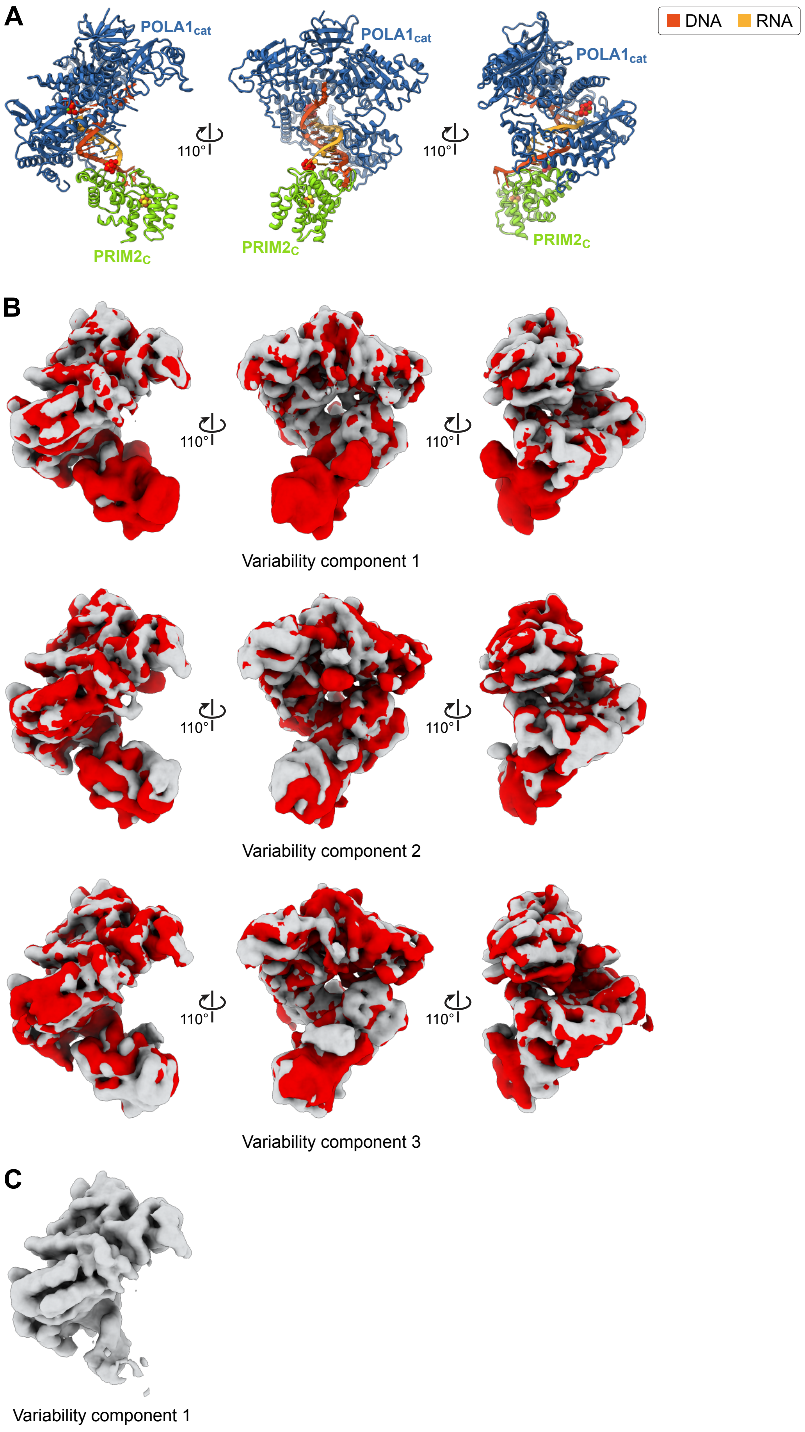


**Supplemental Figure S17. Conformational dynamics in the DNA initiation subcomplex. A**. Structural representations of the DI subcomplex oriented as in panel B. **B**. 3D variability analysis of the 563,079 particles aligned by focused 3D refinement (Supplemental Fig. S10). Maps were filtered to a resolution of 5 Å. Initial and final frames of variability components 1 (top), 2 (middle), and 3 (bottom) are shown in gray and red, respectively. Variability component 1 resolves dissociation of PRIM2$_C$ from the primer/template. Component 2 resolves a bending motion in the oligonucleotide duplex, accompanied by changing interactions between POLA1$_{cat}$ and PRIM2$_C$. Component 3 resolves a second, distinct bending/twisting motion in the duplex. All variability components also resolve conformational changes in the 5′- and 3′-overhangs of the DNA template. **C**. Initial frame of variability component 1 showing dissociation of PRIM2$_C$.

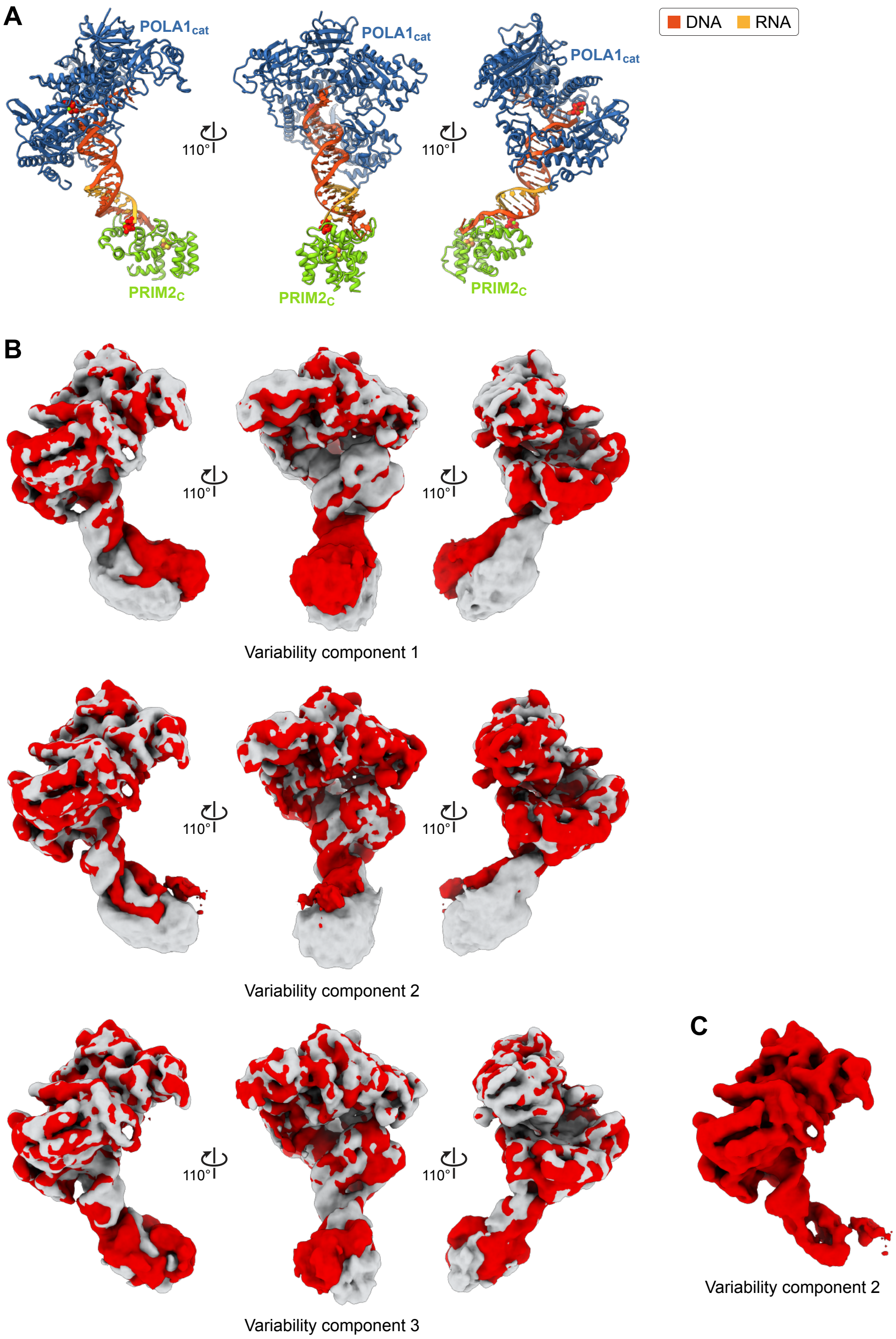


**Supplemental Figure S18. Conformational dynamics in the DNA elongation subcomplex. A**. Structural representations of the DE subcomplex oriented as in panel B. **B**. 3D variability analysis of the 421,934 particles aligned by focused 3D refinement (Supplemental Fig. S12). Maps were filtered to a resolution of 5 Å. Initial and final frames of variability components 1 (top), 2 (middle), and 3 (bottom) are shown in gray and red, respectively. Variability component 1 resolves a bending motion in the duplex, centered at the RNA-DNA junction. Component 2 resolves dissociation of PRIM2$_C$ from the primer/template. Component 3 resolves a second, distinct bending/twisting motion in the duplex. All variability components also resolve conformational changes in the 5′-overhang of the DNA template. Conformational changes in the 3′-overhang are not apparent, but may be obscured by the lower local quality of the reconstructions. **C**. Final frame of variability component 2 showing dissociation of PRIM2$_C$.

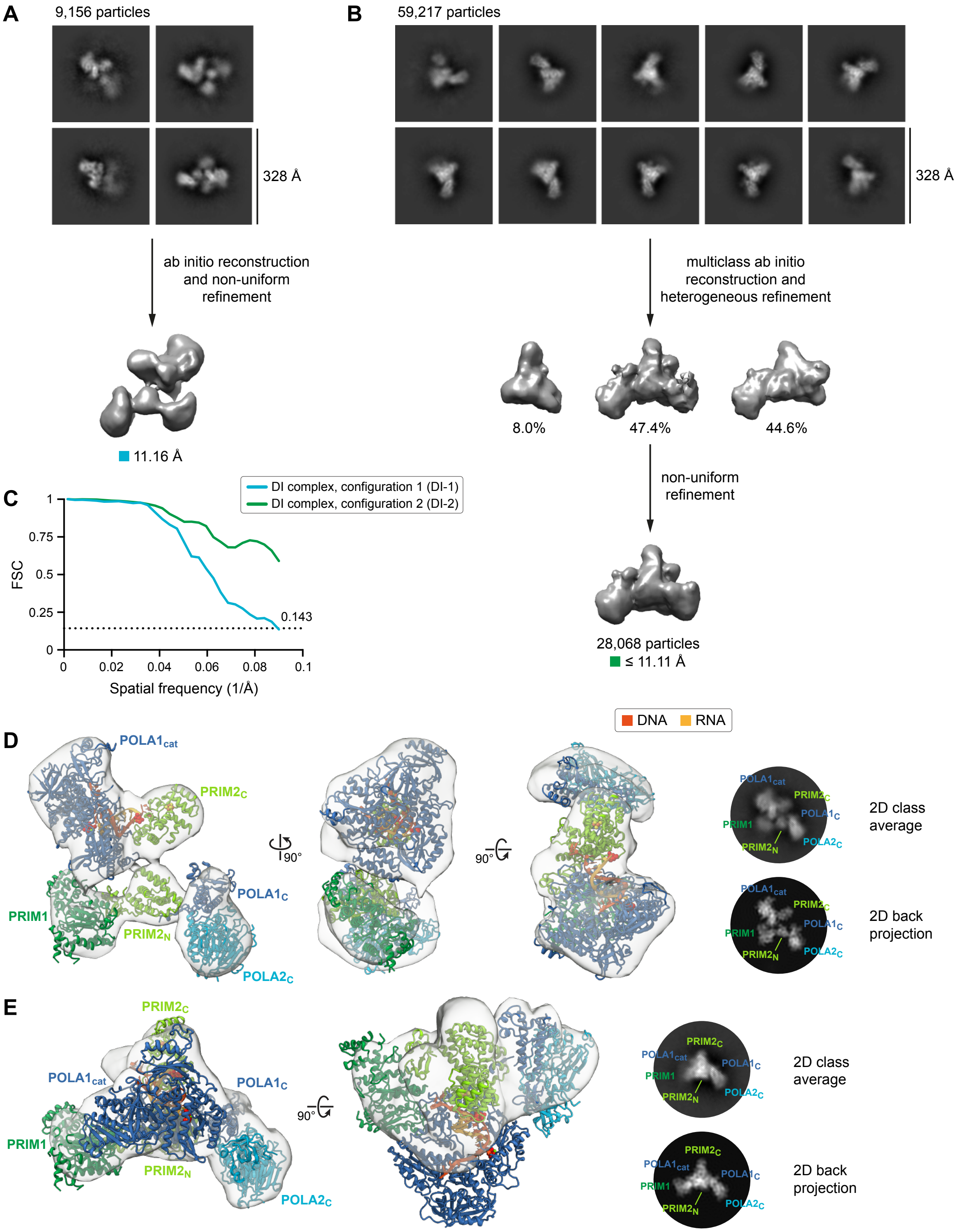


**Supplemental Figure S19. Reconstruction and validation of distinct configurations of the DNA initiation complex. A**,**B**. Data-processing workflows for 3D reconstructions of configurations 1 (A) and 2 (B) of the DI complex. Particles were manually selected from the 2D class averages shown in Supplemental Fig. S11. All processing steps were performed in cryoSPARC. Multiclass ab initio reconstruction and heterogeneous refinement of particles initially assigned to configuration 2 revealed a possible third configuration. **C**. Gold-standard Fourier shell correlation (FSC) curves calculated from independent half-reconstructions. The estimated resolution of configuration DI-2 is limited by the pixel size of the downsampled particles. **D**,**E**. Validation of structures in configurations 1 (D) and 2 (E) of the DI complex. Unsharpened full maps are shown as transparent white surfaces. Structures of the DI subcomplex (one rigid body) and the TC subcomplex (three rigid bodies) were docked into the maps to form the complete DI complex. 2D back projections were generated from the 3D structures and downsampled to a resolution of 10 Å. For configuration DI-2, the discrepancy between the model and the map may be the result of remaining heterogeneity among the particles in the reconstruction or an error in the reconstruction caused by the limited angular distribution of the particles.

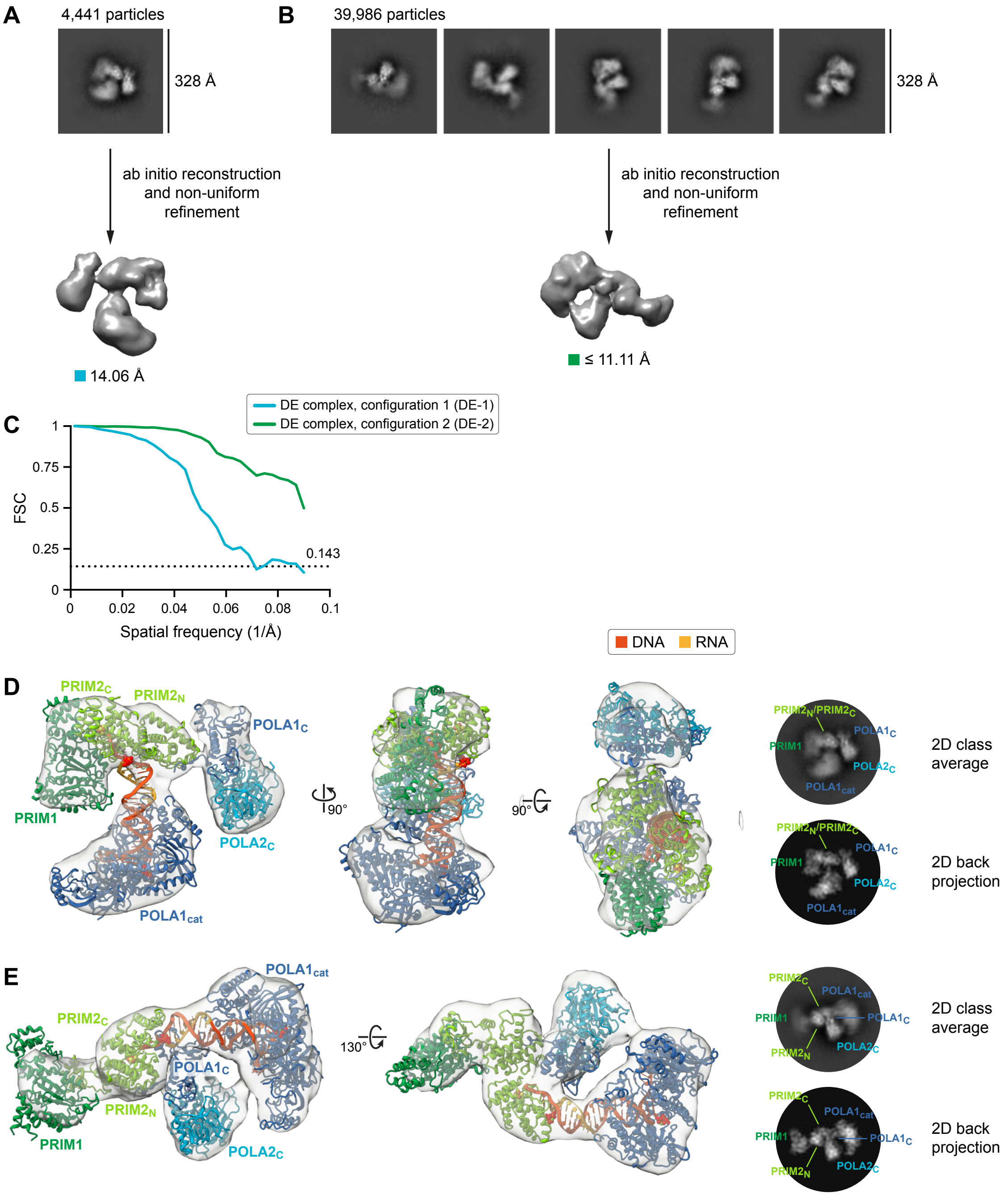


**Supplemental Figure S20. Reconstruction and validation of distinct configurations of the DNA elongation complex. A**,**B**. Data-processing workflows for 3D reconstructions of configurations 1 (A) and 2 (B) of the DE complex. Particles were manually selected from the 2D class averages shown in Supplemental Fig. S13. All processing steps were performed in cryoSPARC. **C**. Gold-standard Fourier shell correlation (FSC) curves calculated from independent half-reconstructions. The estimated resolution of configuration DE-2 is limited by the pixel size of the downsampled particles. **D**,**E**. Validation of structures in configurations 1 (D) and 2 (E) of the DE complex. Unsharpened full maps are shown as transparent white surfaces. Structures of the DE subcomplex (one rigid body) and the TC subcomplex (three rigid bodies) were docked into the maps to form the complete DE complex. 2D back projections were generated from the 3D structures and downsampled to a resolution of 10 Å. For configuration DE-1, the discrepancy in duplex position between the model and the map may be indicative of heterogeneity among the small number of particles in the reconstruction or an error in the reconstruction resulting from the limited angular distribution of the particles.

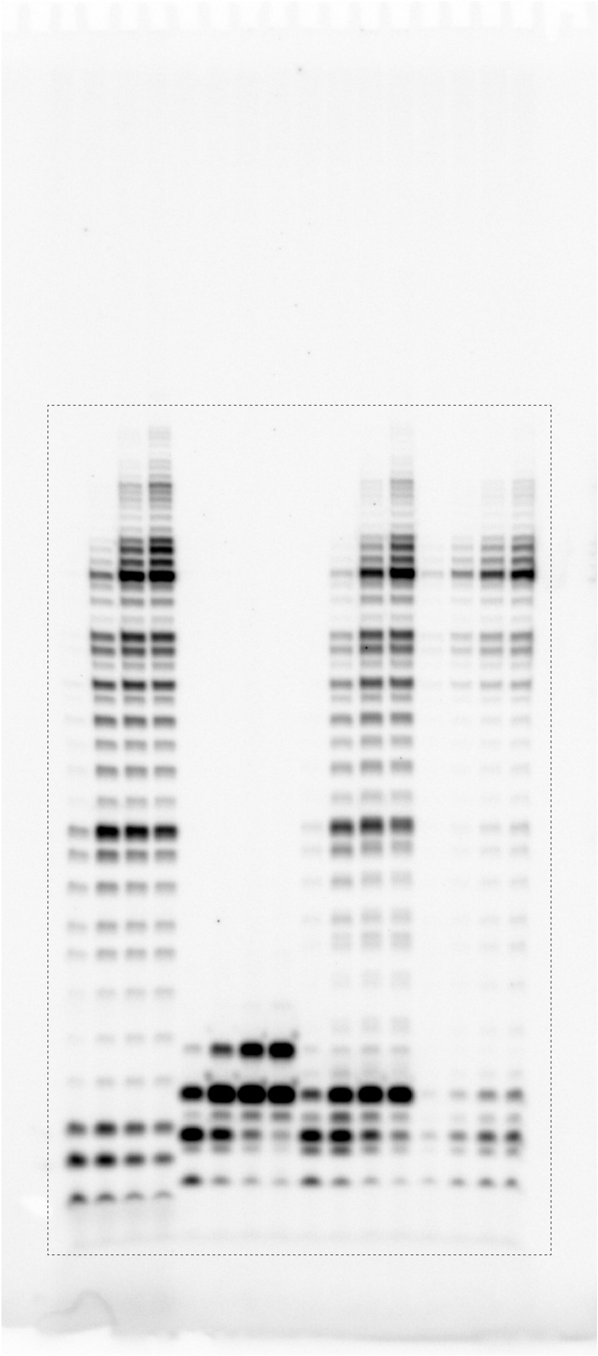

**Supplemental Figure S21. Uncropped gel.** The gel image shown in Fig. 4A was cropped from a larger image. The cropped area is indicated with a dashed line. Unincorporated [α-$^{32}$P]-ATP and [α-$^{32}$P]-dATP were run off the bottom of the gel during electrophoresis.

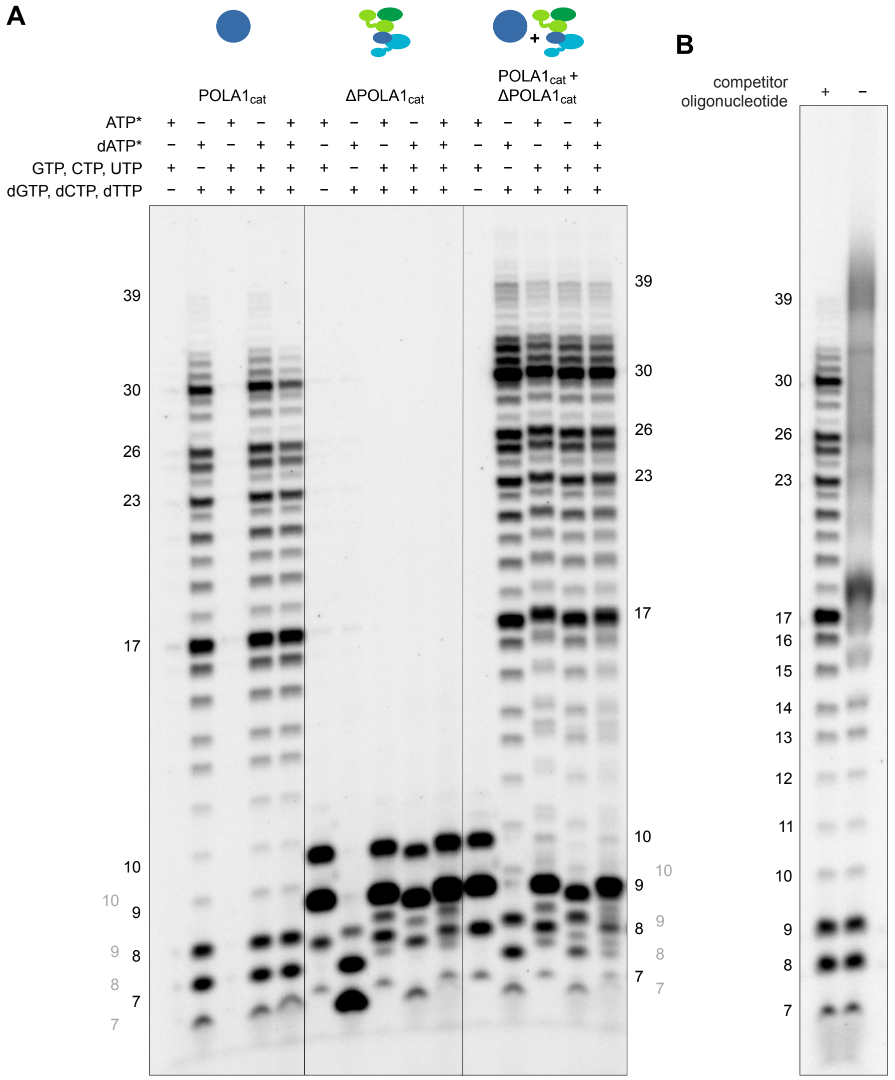


**Supplemental Figure S22. Control experiments for the primer elongation assays. A**. Selective incorporation of NTPs and dNTPs by polα ($POLA1_{cat}$) and primase ($\Delta POLA1_{cat}$). Polα failed to incorporate NTPs at detectable levels. However, primase readily incorporated both NTPs and dNTPs, producing intermediate and doublet bands in reactions with both substrates. Because polα is unable to incorporate NTPs, all DNA elongation products in the reactions containing both $POLA1_{cat}$ and $\Delta POLA1_{cat}$ but lacking dATP were initially extended by primase and then transferred to polα via an intermolecular handoff. **B**. Denaturation of the primer/template. Identical aliquots from the primer elongation assays were worked up with (+) and without (−) a competitor oligonucleotide for subsequent denaturing gel electrophoresis. In the absence of competitor, incomplete denaturation and/or partial re-annealing of the primer/template caused smearing of elongation products ≥ 15 nucleotides in length. In both control experiments, reactions contained the same primer/template depicted in Fig. 4B and were incubated at 22°C for 15 min. Experiments were performed in duplicate.

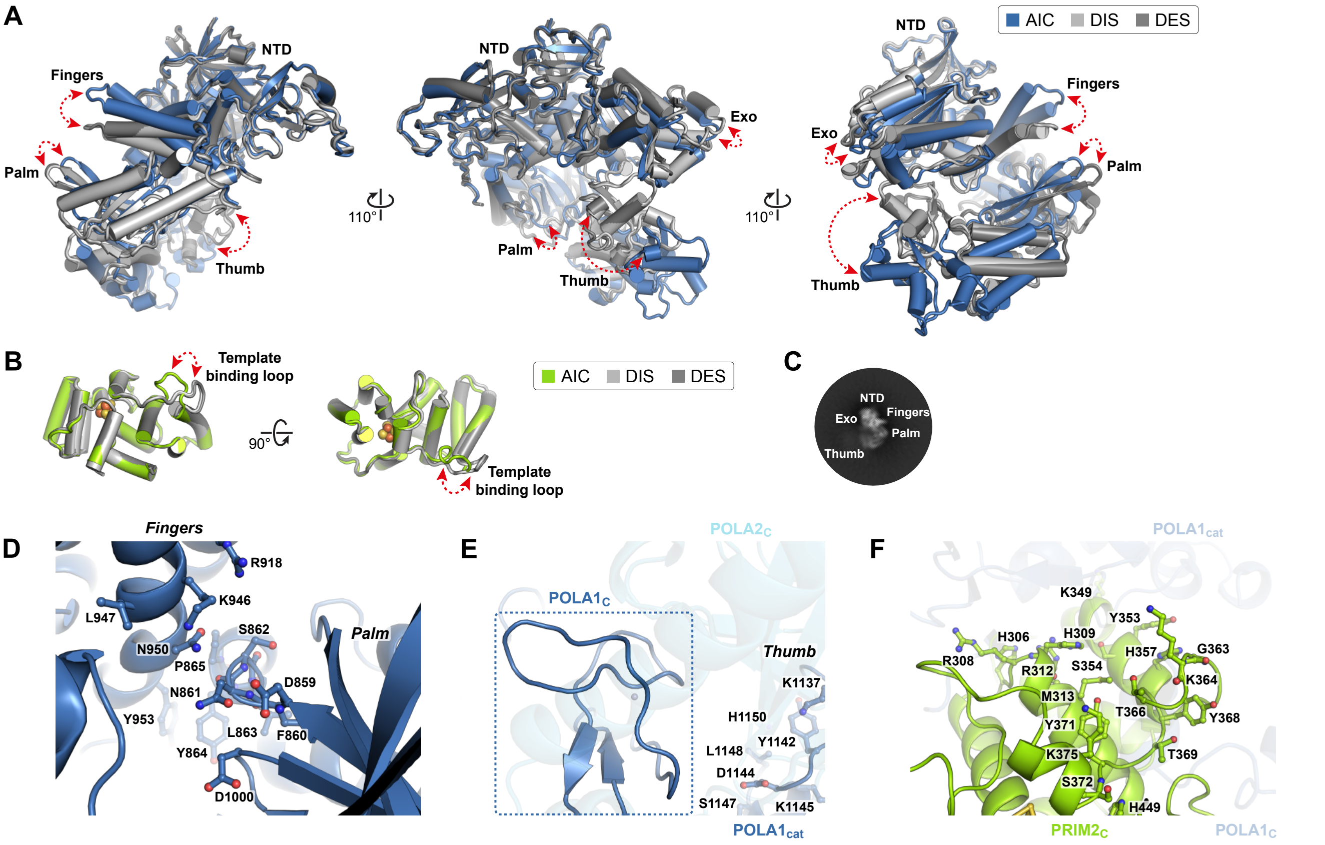


**Supplemental Figure S23. Substrate-dependent conformational changes in POLA1$_{cat}$ and PRIM2$_{C}$. A**,**B**. Comparison of POLA1$_{cat}$ (A) and PRIM2$_{C}$ (B) from the AI, DI, and DE (sub)complexes. Recognition of an RNA/DNA or DNA/DNA duplex by POLA1$_{cat}$ causes conformational changes in the thumb and the palm that in turn form the dNTP binding site. Subsequent recognition of an incoming dNTP results in closure of the fingers, creating a catalytically competent conformation poised for phosphodiester bond formation. Recognition of an RNA/DNA duplex by PRIM2$_{C}$ requires only conformational changes in the template-binding loop. **C**. Selected 2D class average showing POLA1$_{cat}$ from the substrate-free dataset. **D**–**F**. Substrate-binding surfaces in the substrate-free AI complex. Images are oriented as in Fig. 5. **D**. Unformed dNTP-binding pocket created by the fingers and palm domains in the open conformation of POLA1$_{cat}$. **E**. Thumb domain in the open conformation of POLA1$_{cat}$ (blue). The thumb forms protein-protein contacts with POLA1$_{C}$ (blue) and POLA2$_{C}$ (cyan) that sterically occlude substrate binding by POLA1$_{cat}$. **F**. Substrate-binding surface of PRIM2$_{C}$ (green). Helices α17 (residues 344–349) and α18 (residues 352–358) and the template-binding loop (residues 359–372) form protein-protein contacts with POLA1$_{cat}$ (blue) and POLA1$_{C}$ (blue) that sterically block substrate binding by PRIM2$_{C}$.

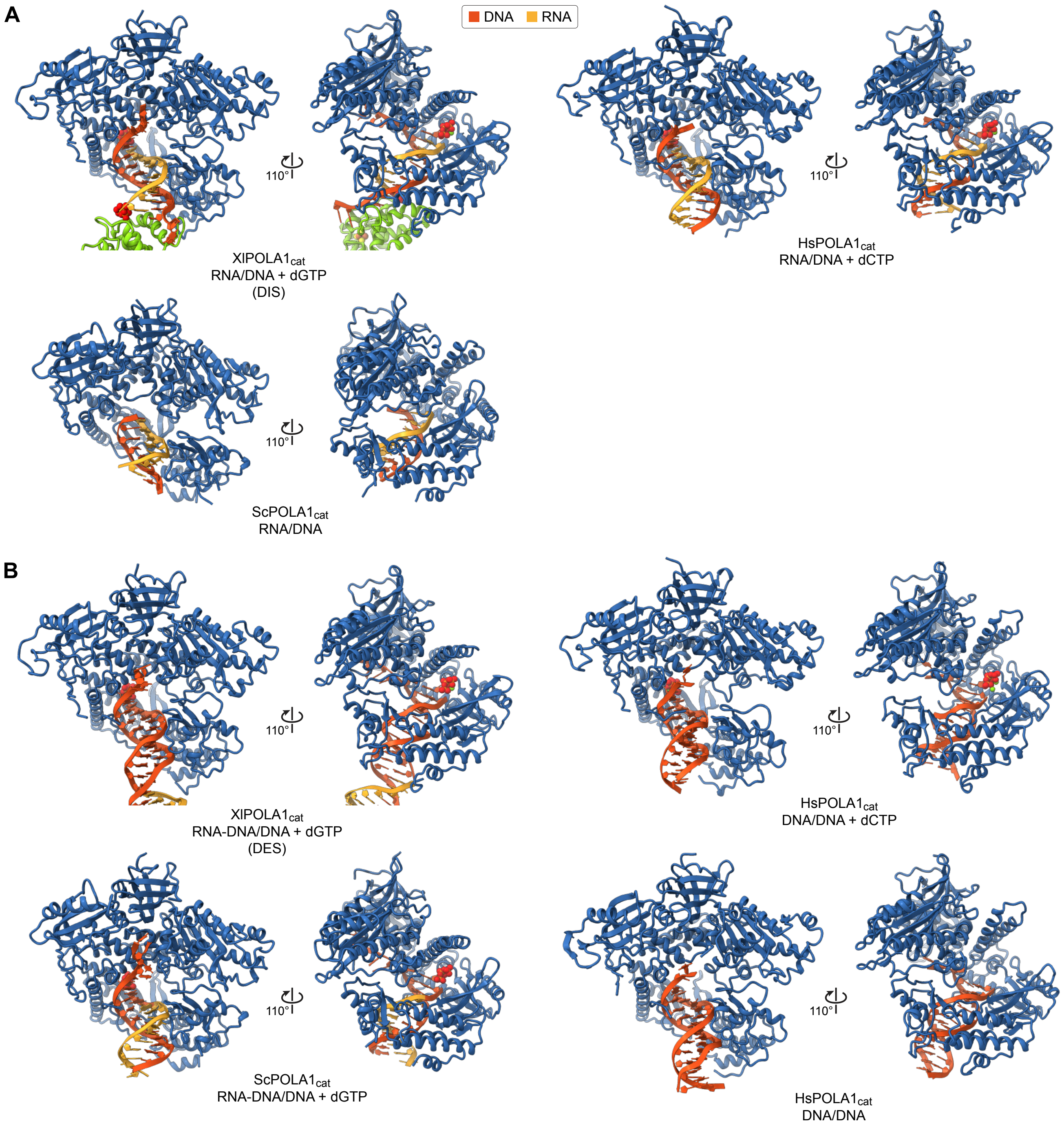


**Supplemental Figure S24. Comparison of substrate-bound POLA1$_{cat}$ structures. A**. DNA initiation complexes containing RNA/DNA duplexes. The *Xenopus laevis* and *Homo sapiens* ternary complexes (PDB accession 4QCL) are fully closed on the primer/template and the incoming dNTP, while the *Saccharomyces cerevisiae* binary complex (PDB accession 4FXD) is partially open in the absence of an incoming dNTP to induce closure of the fingers. POLA1$_{cat}$ and PRIM2$_{C}$ are colored blue and green, respectively. **B**. DNA elongation complexes containing RNA-DNA/DNA or DNA/DNA duplexes. Only the *X. laevis* and S. cerevisiae ternary complexes (PDB accession 4FYD) are fully closed. As expected, in the absence of an incoming dNTP, the *H. sapiens* binary complex (PDB accession 5IUD) is partially open. However, unexpectedly, the *H. sapiens* ternary complex (PDB accession 6AS7) is also partially open. Due to crystallographic lattice contacts, the fingers are not closed on the incoming dCTP.

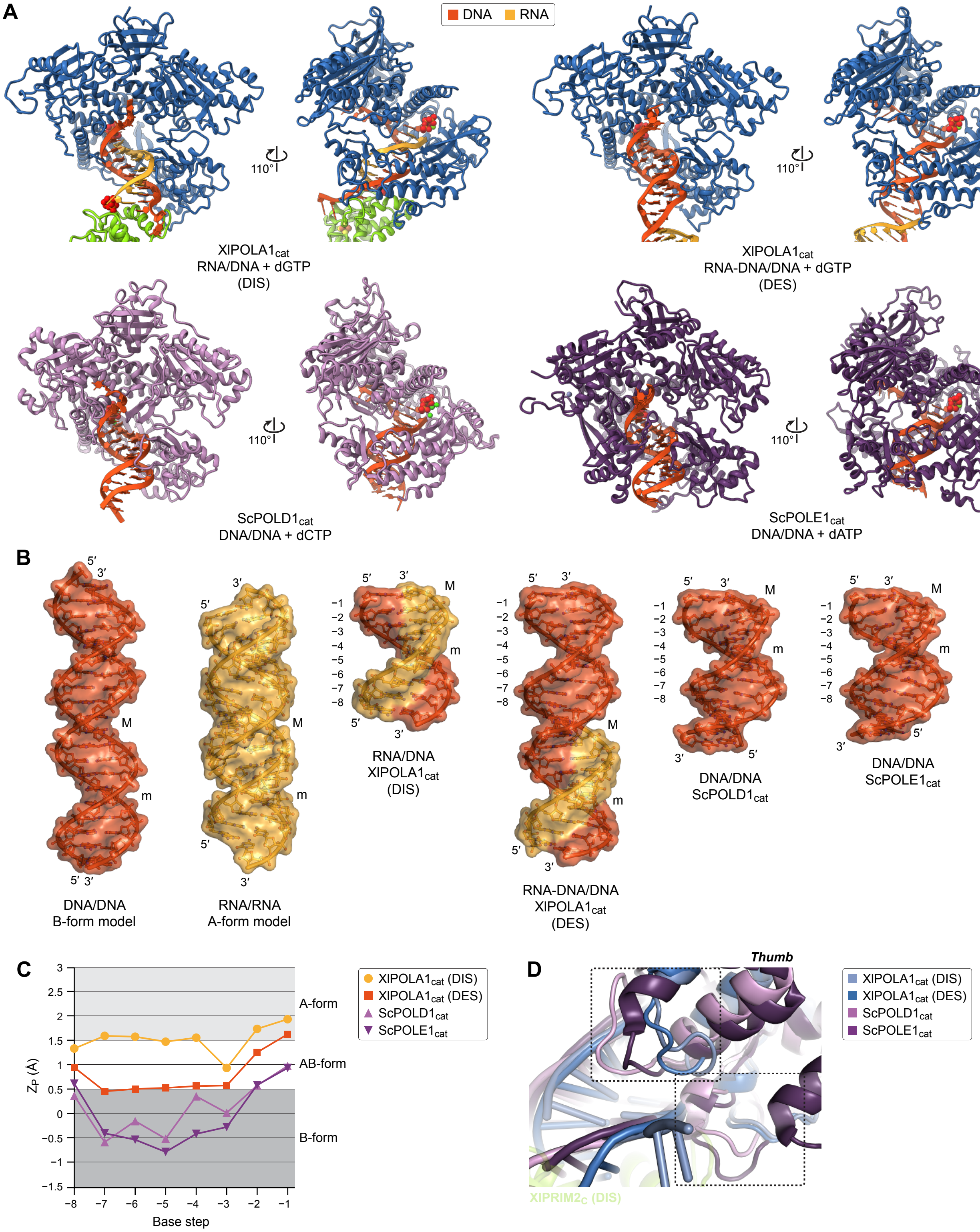


**Supplemental Figure S25. Substrate recognition by polα, polδ, and polε. A**. Comparison of ternary RNA/DNA (DIS) and RNA-DNA/DNA (DES) complexes of *Xenopus laevis* $POLA1_{cat}$ and ternary DNA/DNA complexes of *Saccharomcyes cerevisiae* $POLD1_{cat}$ (PDB accession 3IAY) and $POLE1_{cat}$ (PDB accession 4M8O). $POLA1_{cat}$ and $PRIM2_C$ are colored blue and green, respectively. **B**. Comparison of oligonucleotide duplexes extracted from the structures shown in panel A. Ideal B-form DNA/DNA and A-form RNA/RNA duplexes are shown to the left. Major (M) and minor (m) grooves are indicated. Numbers to the left of the duplexes indicate the base step relative to the 3′-end of the primers. **C**. Analysis of duplex conformation in the structures shown in panels A and B. $Z_P$ is the interstrand phosphate-phosphate distance after projection onto the helical (Z) axis [Hassan and Calladine, *Phil. Trans. R. Soc. Lond. A* (1997) 355: 43; Lu et al, *J. Mol. Biol.* (2000) 300: 819]. **D**. Superposition of the structures shown in panel A. Structural differences in the thumb domain of $POLA1_{cat}$ create a wider substrate-binding cleft for preferential recognition of wider intermediate AB-form duplexes.

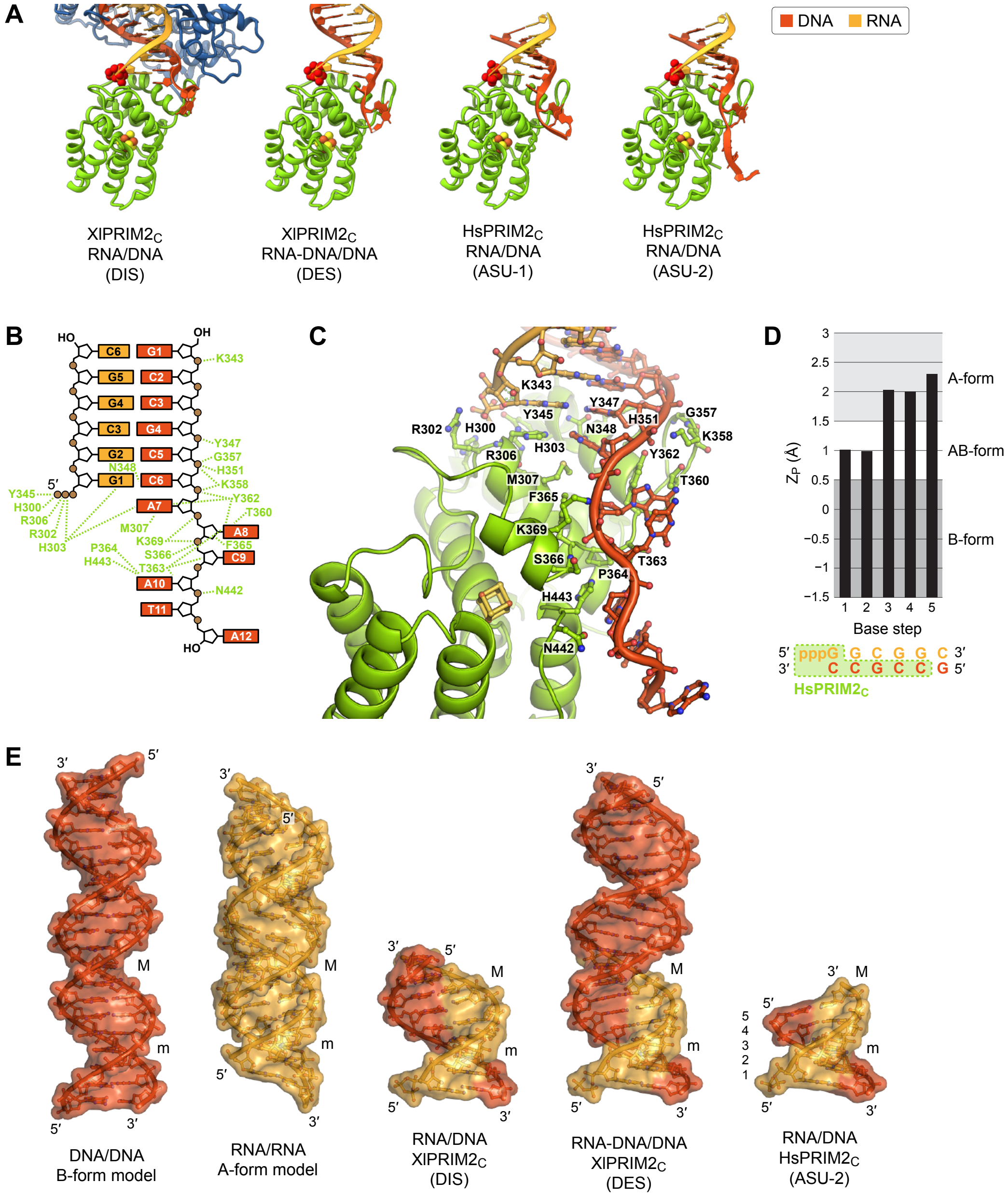


**Supplemental Figure S26. Substrate recognition by PRIM2$_C$. A**. Comparison of *Xenopus laevis* PRIM2$_C$ bound to RNA/DNA (DIS) and RNA-DNA/DNA (DES) duplexes and *Homo sapiens* PRIM2$_C$ bound to an RNA/DNA duplex (PDB accession 5F0Q). The asymmetric unit (ASU) of the HsPRIM2$_C$ crystal structure contains two crystallographically unique complexes (ASU-1 and ASU-2). POLA1$_{cat}$ and PRIM2$_C$ are colored blue and green, respectively. **B**,**C**. Schematic (B) and molecular (C) depictions of substrate-binding interactions with HsPRIM2$_C$ (ASU-2). Due to crystallographic lattice contacts, interactions with the last three nucleotides (A10–A12) of the DNA template differ between the two HsPRIM2$_C$ complexes. **D**. Analysis of duplex conformation in the crystal structure of HsPRIM2$_C$ bound to RNA/DNA (ASU-2). $Z_P$ is the interstrand phosphate-phosphate distance after projection onto the helical (Z) axis [Hassan and Calladine, *Phil. Trans. R. Soc. Lond. A* (1997) 355: 43; Lu et al, *J. Mol. Biol.* (2000) 300: 819]. **E**. Comparison of oligonucleotide duplexes extracted from the structures shown in panel A. Ideal B-form DNA/DNA and A-form RNA/RNA duplexes are shown to the left. Major (M) and minor (m) grooves are indicated. Numbers to the left of the RNA/DNA duplex from the HsPRIM2$_C$ complex (ASU-2) indicate the base step relative to the 5′-end of the primer.

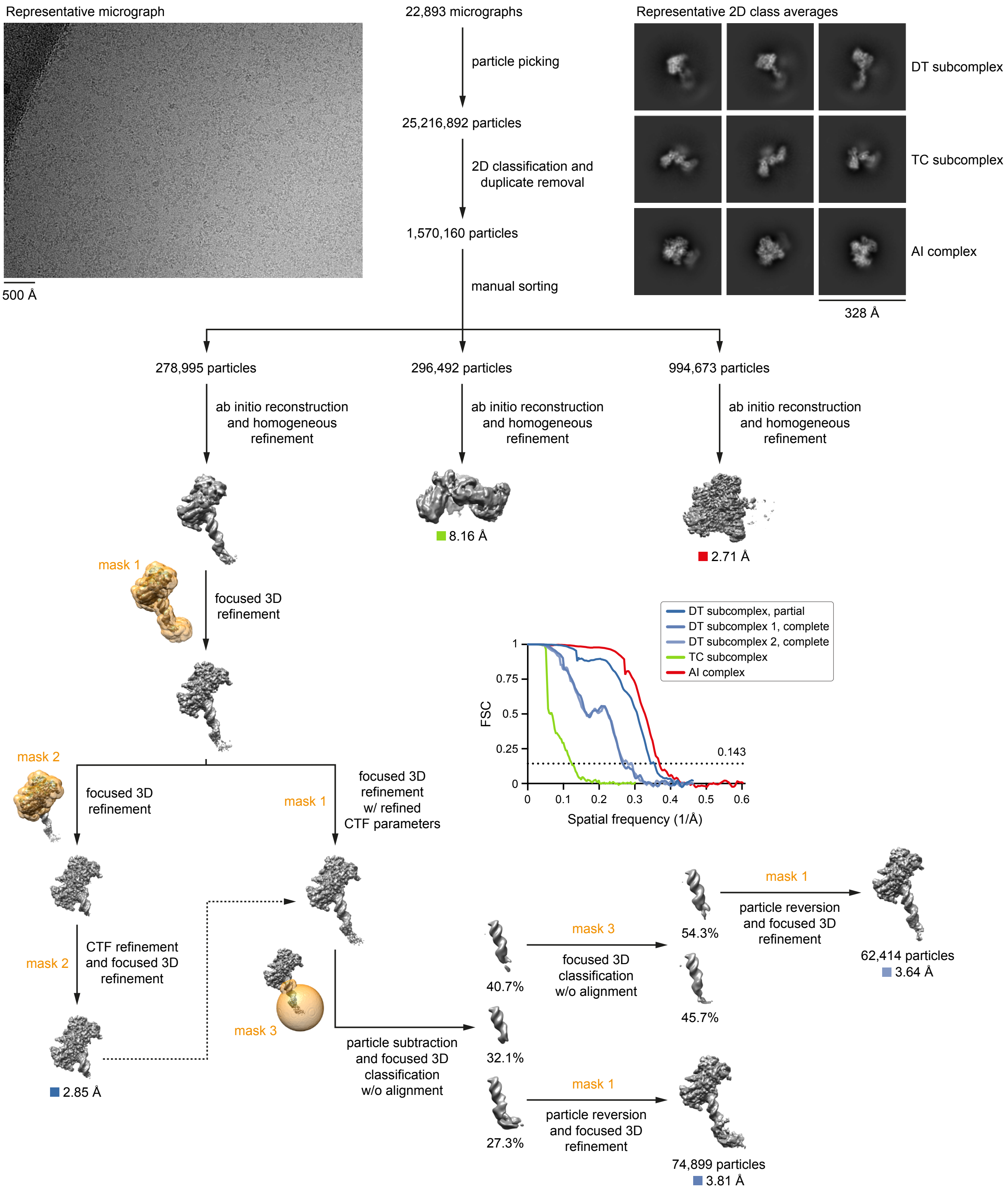


**Supplemental Figure S27. Data-processing workflow for the DNA termination dataset.** All processing steps were performed in cryoSPARC or RELION. 3D reconstructions and masks are colored gray and orange, respectively. Gold-standard Fourier shell correlation (FSC) curves were calculated from independent half-reconstructions. All final 2D class averages are provided in Supplemental Fig. S28. Additional validation metrics are provided in Supplemental Fig. S29 and Supplemental Table S6.

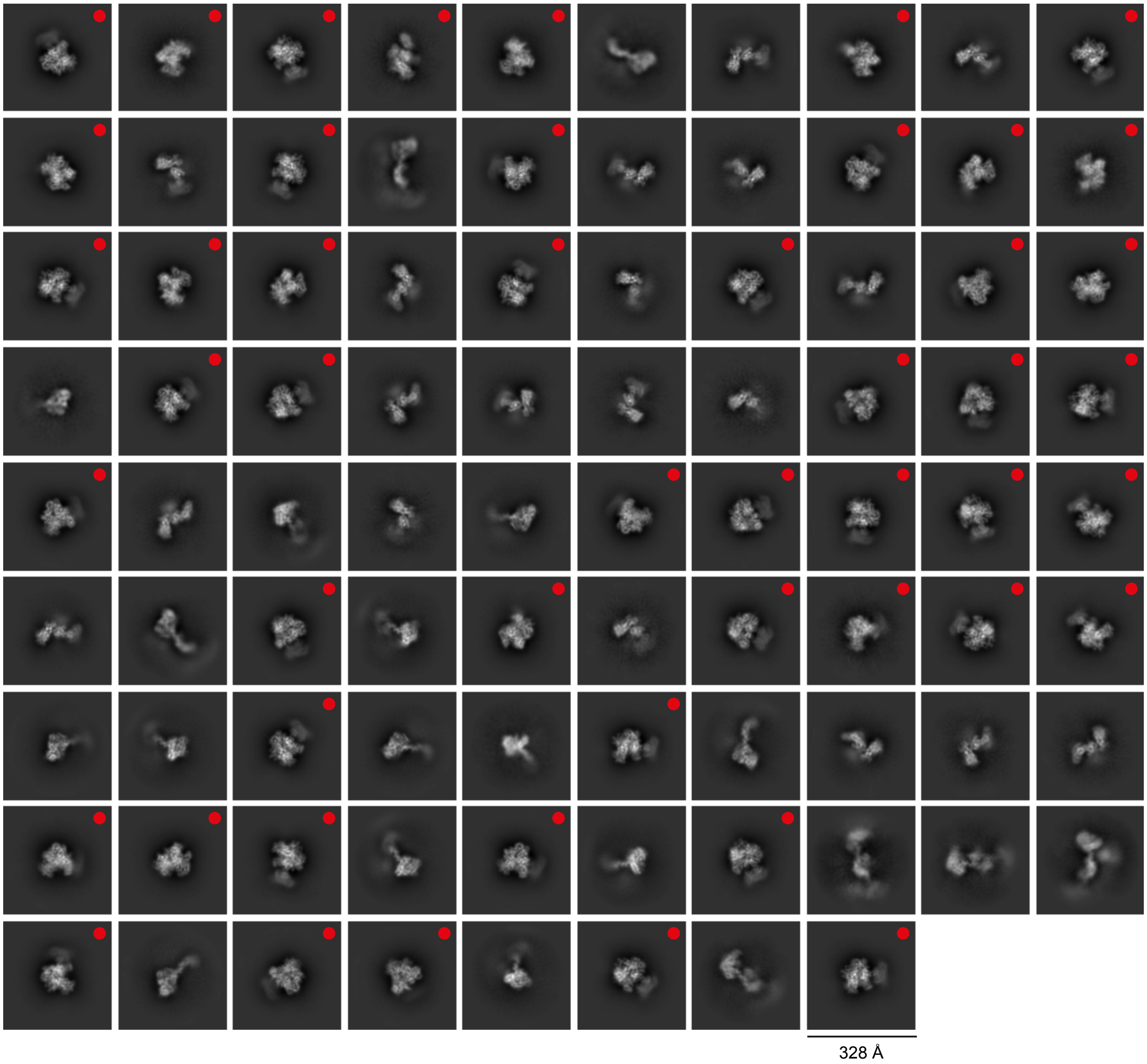


**Supplemental Figure S28. Final 2D class averages from the DNA termination dataset.** 2D class averages show the DT subcomplex (23% of classes, 18% of particles), the TC subcomplex (22% of classes, 19% of particles), or the AI complex (56% of classes, 63% of particles). 2D class averages showing the AI complex are indicated with red dots.

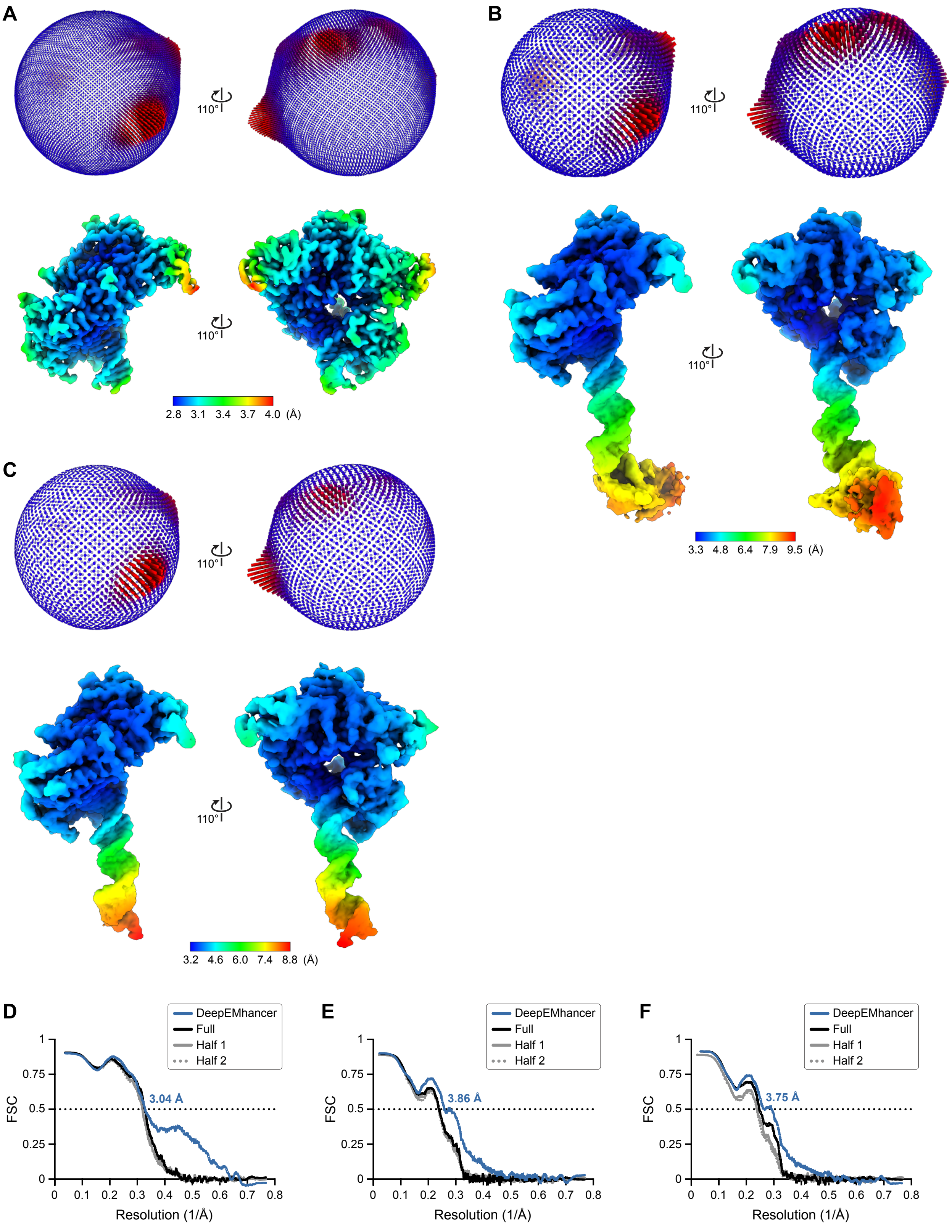
A
110°
2.8 3.1 3.4 3.7 4.0 (Å)
B
110°
3.3 4.8 6.4 7.9 9.5 (Å)
C
110°
3.2 4.6 6.0 7.4 8.8 (Å)
D
DeepEMhancer
Full
Half 1
Half 2
3.04 Å
FSC
Resolution (1/Å)
E
3.86 Å
F
3.75 Å

**(cont.)**

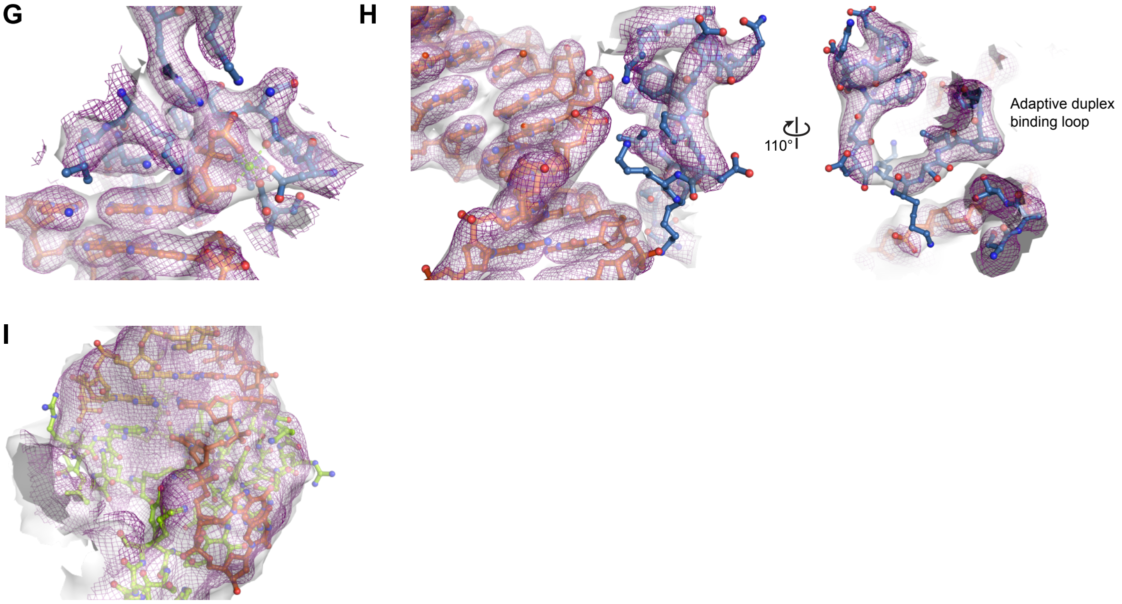


**Supplemental Figure S29. Validation of 3D reconstructions from the DNA termination dataset. A–C**. Euler angle distributions and local resolution estimations for reconstructions of the partial DT subcomplex (A), DT subcomplex 1 (B), and DT subcomplex 2 (C). **D–F**. Model-to-map Fourier shell correlation (FSC) curves for structures of the partial subcomplex (D), subcomplex 1 (E), and subcomplex 2. Prior to correlation calculations, structures were refined against the DeepEMhancer-sharpened map, the unsharpened full map, or half map 1. Correlation with half map 2 was calculated relative to the structures refined against half map 1. **G–I**. Model-to-map fits for selected regions of the DT subcomplex. Cryo-EM densities from the DeepEMhancer-sharpened map and the unsharpened full map are shown as a purple mesh and a white surface, respectively.

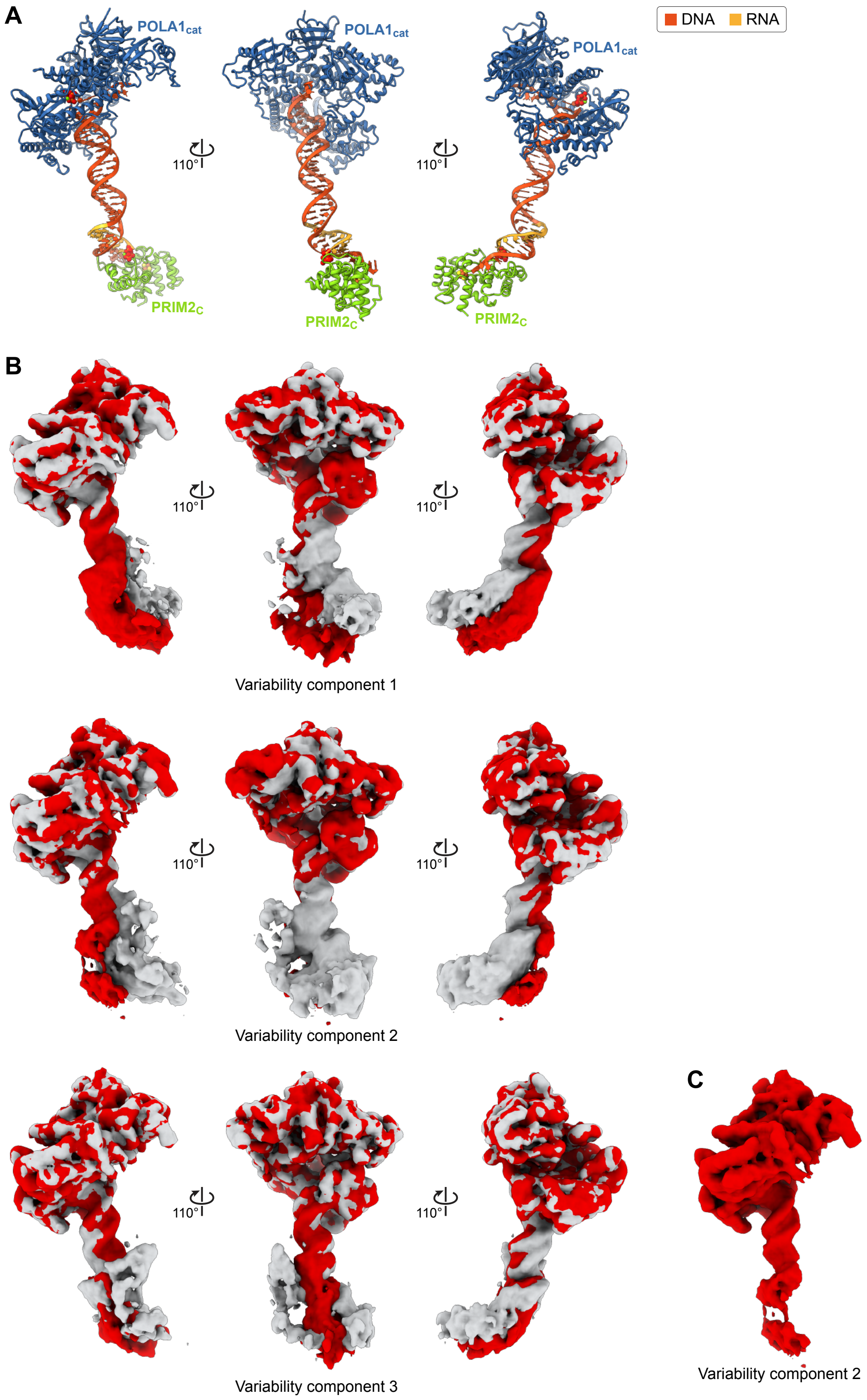


**Supplemental Figure S30. Conformational dynamics in the DNA termination subcomplex. A**. Structural representations of the DT subcomplex oriented as in panel B. **B**. 3D variability analysis of the 278,995 particles aligned by focused 3D refinement (Supplemental Fig. S27). Maps were filtered to a resolution of 5 Å. Initial and final frames of variability components 1 (top), 2 (middle), and 3 (bottom) are shown in gray and red, respectively. Variability component 1 resolves a bending motion in the duplex. Component 2 resolves a similar bending motion accompanied by dissociation of $PRIM2_C$ from the primer/template. Component 3 resolves a third, smaller bending/twisting motion in the duplex. All variability components also resolve conformational changes in the 5′-overhang of the DNA template, while conformational changes in the 3′-overhang are likely obscured by the lower local quality of the reconstructions. Localized noise near $PRIM2_C$ and the unbound portion of the primer/template may be indicative of transient association of the TC subcomplex. Density for $PRIM2_C$ is weak in all components. **C**. Final frame of variability component 2 showing dissociation of $PRIM2_C$.

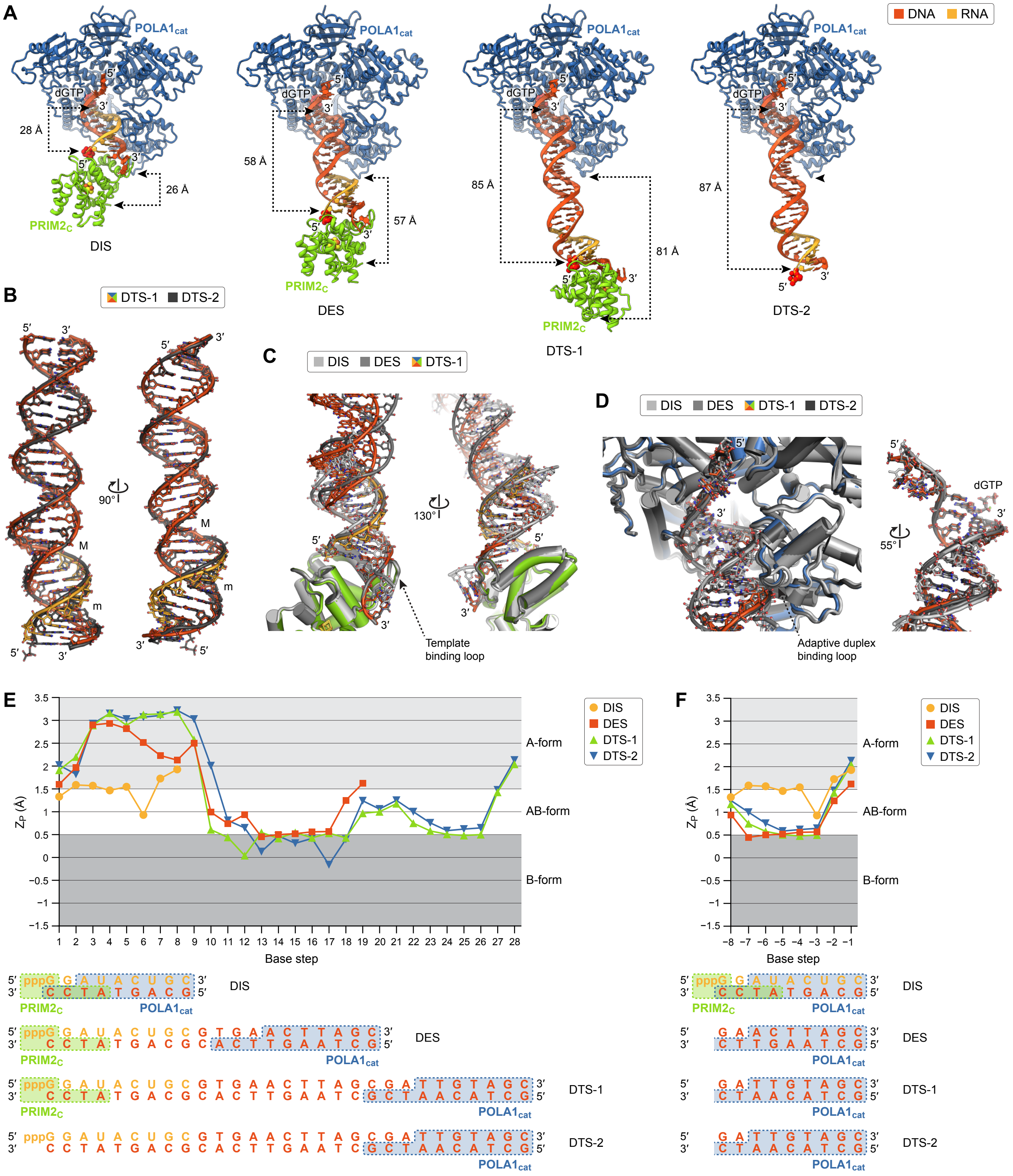
A
POLA1cat
DNA RNA
dGTP
28 Å
26 Å
PRIM2C
DIS
58 Å
57 Å
DES
85 Å
81 Å
DTS-1
87 Å
DTS-2
B
DTS-1 DTS-2
90°
M
m
C
DIS DES DTS-1
130°
Template binding loop
D
DIS DES DTS-1 DTS-2
55°
Adaptive duplex binding loop
E
ZP (Å)
A-form
AB-form
B-form
Base step
F
DIS
DES
DTS-1
DTS-2
5′ pppG G A U A C U G C 3′
3′ C C T A T G A C G 5′
5′ pppG G A U A C U G C G T G A A C T T A G C 3′
3′ C C T A T G A C G C A C T T G A A T C G 5′
5′ pppG G A U A C U G C G T G A A C T T A G C G A T T G T A G C 3′
3′ C C T A T G A C G C A C T T G A A T C G C T A A C A T C G 5′
5′ G A A C T T A G C 3′
3′ C T T G A A T C G 5′
5′ G A T T G T A G C 3′
3′ C T A A C A T C G 5′

(cont.)

**Supplemental Figure S31. Conformational changes in the DNA synthesis assembly. A**. Domain separation during DNA synthesis. As the length of the RNA/DNA duplex increases by 57 Å, the distance between $POLA1_{cat}$ and $PRIM2_C$ (measured at the attachment points of the interdomain linkers) increases by 55 Å. **B**. Conformational differences in the RNA-DNA/DNA duplexes from the DT subcomplexes. In the absence of $PRIM2_C$, the base-pair inclination of the DTS-2 substrate is reduced in the RNA/DNA region of the duplex. **C**. Substrate recognition by $PRIM2_C$. The conformation of $PRIM2_C$ remains unchanged as the conformation of the substrates varies. In the DI subcomplex, duplex remodeling by $POLA1_{cat}$ substantially alters interactions between $PRIM2_C$ and the DNA template. For clarity, $POLA1_{cat}$ is omitted. **D**. Substrate recognition by $POLA1_{cat}$. $POLA1_{cat}$ remodels all substrates to achieve similar (near) intermediate AB-form conformations. Modest narrowing of the minor groove (DIS, 15.1 Å; DES, 13.5 Å; DTS-1, 12.9 Å; DTS-2, 13.4 Å) in the DNA/DNA region of the DE and DT substrates is correlated with conformational changes in the adaptive duplex-binding loop. For clarity, $PRIM2_C$ is omitted. **E**,**F**. Conformational analysis of the DI, DE, and DT substrates. $Z_P$ is the interstrand phosphate-phosphate distance after projection onto the helical (Z) axis [Hassan and Calladine, Phil. Trans. R. Soc. Lond. A (1997) 355: 43; Lu et al, J. Mol. Biol. (2000) 300: 819]. The base step indicated in the plots is relative to either the 5′-end (E) or 3′-end (F) of the primer. The regions of the duplexes bound by $POLA1_{cat}$ (blue) and $PRIM2_C$ (green) are shown below the plots.

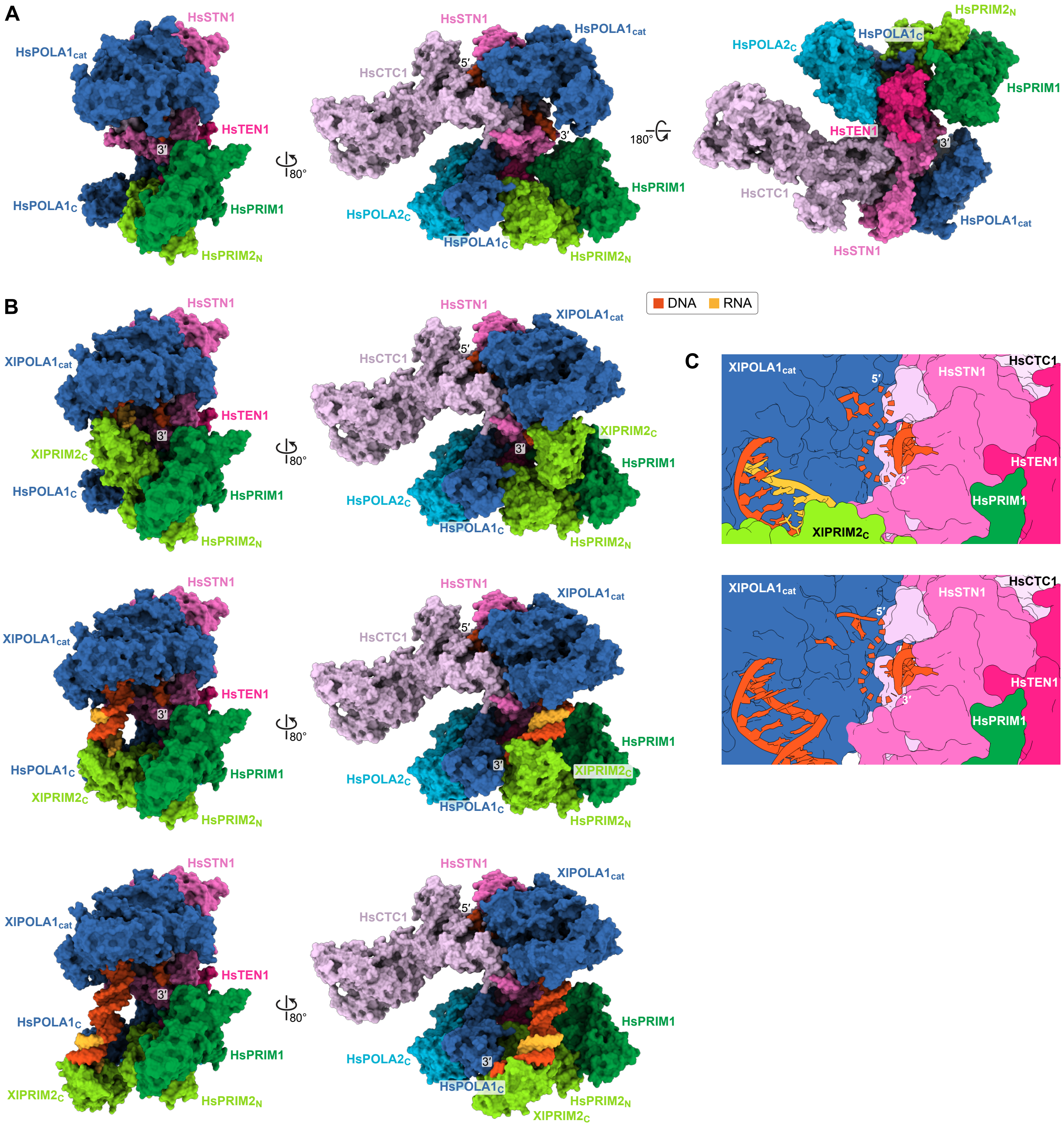


**Supplemental Figure S32. Structural models for DNA synthesis by the polα–primase/CST complex. A**. Cryo-EM structure of *Homo sapiens* polα–primase/CST bound to single-stranded DNA in a pre-initiation complex (PDB accession 8D0B). Due to weak density resulting from conformational heterogeneity, neither the C-terminal domain of PRIM2 nor the thumb domain of POLA1$_{cat}$ were modeled. **B**. Chimeric models of polα–primase/CST bound to primer/template in hypothetical DNA initiation (top), DNA elongation (middle), and DNA termination (bottom) complexes. Models were constructed by aligning POLA1$_{cat}$ in the *Xenopus laevis* DI, DE, and DT subcomplexes to POLA1$_{cat}$ in the *H. sapiens* polα–primase/CST complex. In the closed, substrate-bound conformation of XlPOLA1$_{cat}$, the thumb domain forms additional protein-protein contacts with HsCTC1 and HsSTN1. In the DI complex, XlPRIM2$_{C}$ fits between XlPOLA1$_{cat}$, HsPRIM1, and HsPRIM2$_{N}$. In the DE complex, XlPRIM2$_{C}$ is moved away from XlPOLA1$_{cat}$ and HsPRIM1 and into contact with HsPRIM2$_{N}$, creating a severe steric clash. In the DT complex, XlPRIM2$_{C}$ is moved past HsPRIM2$_{N}$. **C**. Cross-sections through the hypothetical DNA initiation (top) and DNA elongation (bottom) complexes shown in panel B. The dashed lines connect the DNA template in the *H. sapiens* polα–primase/CST complex to the DNA templates in the *X. laevis* DI and DE subcomplexes.

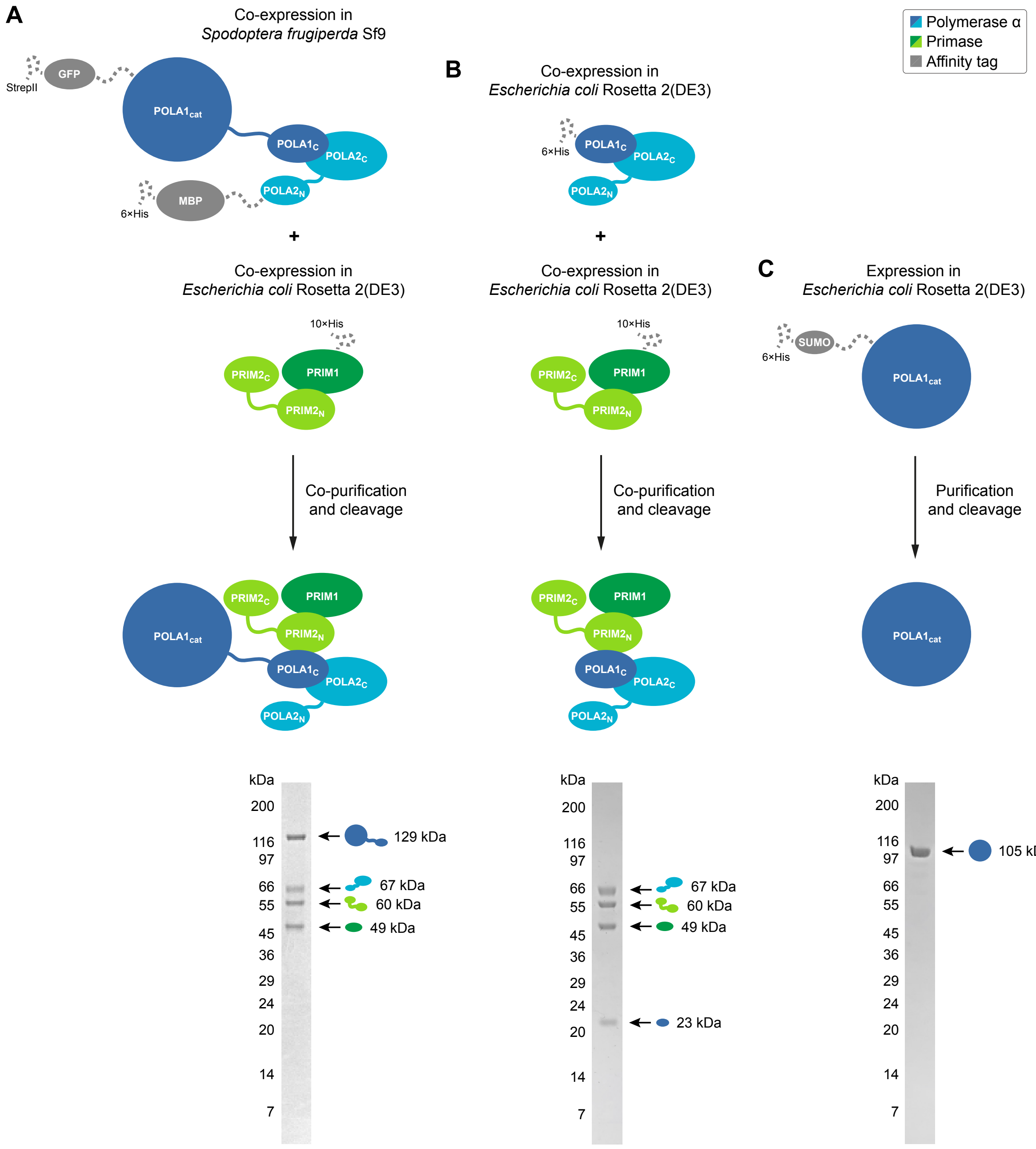


**Supplemental Figure S33. Polα–primase constructs. A**–**C**. Expression and purification of polα–primase (A), ΔPOLA1$_{cat}$ (B), and POLA1$_{cat}$ (C). Protein constructs were (co-)expressed in either *Spodoptera frugiperda* Sf9 or *Escherichia coli* Rosetta2(DE3) cells and then (co-)purified. Affinity tags were cleaved during purification. Purified proteins were characterized by denaturing gel electrophoresis.

**Supplemental Table S1.** Cryo-EM data collection, processing, refinement, and validation statistics for the substrate-free dataset.

| | AI complex, partial (EMD-29862) (PDB 8G99) | AI complex, complete (EMD-29864) (PDB 8G9F) | TC subcomplex, conformation 1 (EMD-29888) | TC subcomplex, conformation 2 (EMD-29889) | TC subcomplex, conformation 3 (EMD-29891) |
|---|---|---|---|---|---|
| **Data collection and processing** | | | | | |
| Magnification (×) | 105,000 | 105,000 | 105,000 | 105,000 | 105,000 |
| Voltage (kV) | 300 | 300 | 300 | 300 | 300 |
| Electron exposure (e−/Å$^2$) | 54.8 | 54.8 | 54.8 | 54.8 | 54.8 |
| Defocus range (μm) | −0.8 – −2.0 | −0.8 – −2.0 | −0.8 – −2.0 | −0.8 – −2.0 | −0.8 – −2.0 |
| Pixel size (Å) | 0.82 | 0.82 | 0.82 | 0.82 | 0.82 |
| Symmetry imposed | C1 | C1 | C1 | C1 | C1 |
| Initial particle images (no.)* | 23,007,566 | 23,007,566 | 23,007,566 | 23,007,566 | 23,007,566 |
| Particle images after 2D classification (no.)[†] | 2,683,442 | 2,683,442 | 2,683,442 | 2,683,442 | 2,683,442 |
| Final particle images (no.) | 334,066 | 47,277 | 64,911 | 80,155 | 68,842 |
| Map resolution (Å) | 2.80 | 3.22 | 9.22 | 8.56 | 8.99 |
| FSC threshold | 0.143 | 0.143 | 0.143 | 0.143 | 0.143 |
| Map resolution range (Å) | 2.6 – 7.0 | 2.9 – 5.9 | — | — | — |
| **Structure refinement and validation** | | | | | |
| Initial model used (PDB code) | 5EXR | 4BPU, 8G99 | 8G9F | 8G9F | 8G9F |
| Model resolution (Å) | 2.66 | 3.05 | — | — | — |
| FSC threshold | 0.5 | 0.5 | — | — | — |
| Map sharpening *B*-factor (Å$^2$)[‡] | — | — | — | — | — |
| Model composition (no.) | | | | | |
| Non-hydrogen atoms | 15,096 | 18,782 | — | — | — |
| Protein residues | 1,875 | 2,321 | — | — | — |
| RNA/DNA nucleotides | 0 | 0 | — | — | — |
| Cofactors/ions | 3 | 4 | — | — | — |
| Avg. *B*-factors (Å$^2$) | | | | | |
| Protein | 51.0 | 75.8 | — | — | — |
| RNA/DNA | — | — | — | — | — |
| Cofactor/ion | 67.5 | 108.9 | — | — | — |
| R.m.s. deviations | | | | | |
| Bond lengths (Å) | 0.003 | 0.003 | — | — | — |
| Bond angles (°) | 0.458 | 0.491 | — | — | — |
| Ramachandran plot (%) | | | | | |
| Favored | 96.93 | 96.91 | — | — | — |
| Allowed | 3.07 | 3.09 | — | — | — |
| Disallowed | 0.00 | 0.00 | — | — | — |
| MolProbity score | 1.19 | 1.25 | — | — | — |
| Clashscore | 2.32 | 2.77 | — | — | — |
| Poor rotamers (%) | 0.00 | 0.00 | — | — | — |

*Particles were picked using three separate methods. Initial particle images include duplicates and triplicates.

[†]Duplicate and triplicate particle images were removed before final 2D classification.

[‡]Maps were locally sharpened using DeepEMhancer.

**Supplemental Table S2.** Oligonucleotide constructs.

| Sequence* | Strand | Application |
|---|---|---|
| pGAUACUGCdd | Primer | Synthesis (DI) |
| GTCTCACACAGCAGUAUCCAA | Template | Synthesis (DI) |
| UUGGAUACUGCTGTGTGAGAC | Competitor | Synthesis (DI) |
| pGAUACUGCGTGAACTTAGCdd | Primer | Synthesis (DE) |
| GCAGUAUCCAA | Template | Synthesis (DE) |
| UUGGAUACUGC | Competitor | Synthesis (DE) |
| pGAUACUGCGTGAACTTAGCGATTGTAGCdd | Primer | Synthesis (DT) |
| GCAGUAUCCAA | Template | Synthesis (DT) |
| UUGGAUACUGC | Competitor | Synthesis (DT) |
| pGCGGC | Primer | Synthesis |
| GCAGCCGCCAA | Template | Synthesis |
| UUGGCGGCUGC | Competitor | Synthesis |
| pppGGAUACUGCdd | Primer | Cryo-EM (DI) |
| TGTATGTATGTATGTCGCTAAGTTCACGCAGTATCCTGTATGTATGTATG | Template | Cryo-EM (DI) |
| pppGGAUACUGCGTGAACTTAGCdd | Primer | Cryo-EM (DE) |
| TGTATGTATGTATGTCGCTAAGTTCACGCAGTATCCTGTATGTATGTATG | Template | Cryo-EM (DE) |
| pppGGAUACUGCGTGAACTTAGCGATTGTAGCdd | Primer | Cryo-EM (DT) |
| TGTATGTATGTATGTCGCTACAATCGCTAAGTTCACGCAGTATCCTGTATGTATGTATG | Template | Cryo-EM (DT) |
| pppGGCGGC | Primer | Elongation |
| ACACAGGAACAGGAGAACACAGGAACAGAGGGACGAAGGACAGAAGGGACAGAAGACAG GAGAAACAGACGACAAGATGCCGCCACAGAGACACAGAGA | Template | Elongation |
| TCTCTGTGTCTCTGTGGCGGCATCTTGTCGTCTGTTTCTCCTGTCTTCTGTCCCTTCTGTCC TTCGTCCCTCTGTTCCTGTGTTCTCCTGTTCCTGTGT | Competitor | Elongation |
| GGATACTGCdd | Primer | Binding |
| Cy5-TGTATGTATGTATGTCGCAGTATCCTGTAT | Template | Binding |
| pppGGAUACUGCdd | Primer | Binding (DI) |
| Cy5-TGTATGTATGTATGTCGCAGTATCCTGTAT | Template | Binding (DI) |
| pppGGAUACUGCGTGAACTTAGCdd | Primer | Binding (DE) |
| Cy5-TGTATGTATGTATGTCGCTAAGTTCACGCAGTATCCTGTAT | Template | Binding (DE) |
| pppGGAUACUGCGTGAACTTAGCGATTGTAGCdd | Primer | Binding (DT) |
| Cy5-TGTATGTATGTATGTCGCTACAATCGCTAAGTTCACGCAGTATCCTGTAT | Template | Binding (DT) |

*Sequences are listed 5′→3′ with RNA and DNA colored gray and black, respectively. 5′-Monophosphorylated and triphosphorylated nucleotides are indicated with "p" and "ppp". 2′,3′-Dideoxynucleotides are indicated with "dd". Constructs used in multiple applications and/or as part of multiple substrates are listed multiple times.

**Supplemental Table S3.** Cryo-EM data collection, processing, refinement, and validation statistics for the DNA initiation dataset.

| | DI subcomplex<br>(EMD-29871)<br>(PDB 8G9L) | TC subcomplex | DI complex, configuration 1 (DI-1) | DI complex, configuration 2 (DI-2) |
|---|---|---|---|---|
| **Data collection and processing** | | | | |
| Magnification (×) | 45,000 | 45,000 | 45,000 | 45,000 |
| Voltage (kV) | 200 | 200 | 200 | 200 |
| Electron exposure (e−/Å$^2$) | 45.5, 50.6, 52.3, 54.4 | 45.5, 50.6, 52.3, 54.4 | 45.5, 50.6, 52.3, 54.4 | 45.5, 50.6, 52.3, 54.4 |
| Defocus range (μm) | −1.5 – −2.5 | −1.5 – −2.5 | −1.5 – −2.5 | −1.5 – −2.5 |
| Pixel size (Å) | 0.91 | 0.91 | 0.91 | 0.91 |
| Symmetry imposed | C1 | C1 | C1 | C1 |
| Initial particle images (no.)* | 11,200,341 | 11,200,341 | 11,200,341 | 11,200,341 |
| Particle images after 2D classification (no.)† | 1,191,043 | 1,191,043 | 1,191,043 | 1,191,043 |
| Final particle images (no.) | 202,946 | 63,237 | 9,156 | 28,068 |
| Map resolution (Å) | 3.31 | 9.71 | 11.16 | ≤ 11.11 |
| FSC threshold | 0.143 | 0.143 | 0.143 | 0.143 |
| Map resolution range (Å) | 3.1 – 4.9 | — | — | — |
| **Structure refinement and validation** | | | | |
| Initial model used (PDB code) | 4QCL, 5F0Q | — | 8G9F, 8G9L | 8G9F, 8G9L |
| Model resolution (Å) | 3.20 | — | — | — |
| FSC threshold | 0.5 | — | — | — |
| Map sharpening *B*-factor (Å$^2$)‡ | — | — | — | — |
| Model composition (no.) | | | | |
| Non-hydrogen atoms | 9,027 | — | — | — |
| Protein residues | 1,054 | — | — | — |
| RNA/DNA nucleotides | 26 | — | — | — |
| Cofactors/ions | 3 | — | — | — |
| Avg. *B*-factors (Å$^2$) | | | | |
| Protein | 75.6 | — | — | — |
| RNA/DNA | 68.7 | — | — | — |
| Cofactor/ion | 98.9 | — | — | — |
| R.m.s. deviations | | | | |
| Bond lengths (Å) | 0.002 | — | — | — |
| Bond angles (°) | 0.483 | — | — | — |
| Ramachandran plot (%) | | | | |
| Favored | 96.08 | — | — | — |
| Allowed | 3.92 | — | — | — |
| Disallowed | 0.00 | — | — | — |
| MolProbity score | 1.33 | — | — | — |
| Clashscore | 2.75 | — | — | — |
| Poor rotamers (%) | 0.00 | — | — | — |

*Particles were picked using three separate methods. Initial particle images include duplicates and triplicates.

†Duplicate and triplicate particle images were removed before final 2D classification.

‡Maps were locally sharpened using DeepEMhancer.

**Supplemental Table S4.** Cryo-EM data collection, processing, refinement, and validation statistics for the DNA elongation dataset.

| | DE subcomplex, partial (EMD-29872) (PDB 8G9N) | DE subcomplex, complete (EMD-29873) (PDB 8G9O) | TC subcomplex, conformation 1 | TC subcomplex, conformation 2 | DE complex, configuration 1 (DE-1) | DE complex, configuration 2 (DE-2) |
|---|---|---|---|---|---|---|
| **Data collection and processing** | | | | | | |
| Magnification (×) | 45,000 | 45,000 | 45,000 | 45,000 | 45,000 | 45,000 |
| Voltage (kV) | 200 | 200 | 200 | 200 | 200 | 200 |
| Electron exposure (e−/Å$^2$) | 41.7, 53.0, 55.2 | 41.7, 53.0, 55.2 | 41.7, 53.0, 55.2 | 41.7, 53.0, 55.2 | 41.7, 53.0, 55.2 | 41.7, 53.0, 55.2 |
| Defocus range (μm) | −1.5 – −2.5 | −1.5 – −2.5 | −1.5 – −2.5 | −1.5 – −2.5 | −1.5 – −2.5 | −1.5 – −2.5 |
| Pixel size (Å) | 0.91 | 0.91 | 0.91 | 0.91 | 0.91 | 0.91 |
| Symmetry imposed | C1 | C1 | C1 | C1 | C1 | C1 |
| Initial particle images (no.)* | 8,928,051 | 8,928,051 | 8,928,051 | 8,928,051 | 8,928,051 | 8,928,051 |
| Particle images after 2D classification (no.)† | 934,013 | 934,013 | 934,013 | 934,013 | 934,013 | 934,013 |
| Final particle images (no.) | 421,934 | 42,953 | 114,051 | 72,017 | 4,441 | 39,986 |
| Map resolution (Å) | 3.49 | 4.37 | 9.34 | 9.71 | 14.06 | ≤ 11.11 |
| FSC threshold | 0.143 | 0.143 | 0.143 | 0.143 | 0.143 | 0.143 |
| Map resolution range (Å) | 3.3 – 4.7 | 3.9 – 7.5 | — | — | — | — |
| **Structure refinement and validation** | | | | | | |
| Initial model used (PDB code) | 8G9L | 5F0Q, 8G9L, 8G9N | — | — | 8G9F, 8G9O | 8G9F, 8G9O |
| Model resolution (Å) | 3.47 | 4.41 | — | — | — | — |
| FSC threshold | 0.5 | 0.5 | — | — | — | — |
| Map sharpening *B*-factor (Å$^2$)‡ | — | — | — | — | — | — |
| Model composition (no.) | | | | | | |
| Non-hydrogen atoms | 7,427 | 9,481 | — | — | — | — |
| Protein residues | 868 | 1,054 | — | — | — | — |
| RNA/DNA nucleotides | 23 | 48 | — | — | — | — |
| Cofactors/ions | 1 | 3 | — | — | — | — |
| Avg. *B*-factors (Å$^2$) | | | | | | |
| Protein | 79.1 | 160.0 | — | — | — | — |
| RNA/DNA | 57.7 | 97.1 | — | — | — | — |
| Cofactor/ion | 70.0 | 203.3 | — | — | — | — |
| R.m.s. deviations | | | | | | |
| Bond lengths (Å) | 0.003 | 0.003 | — | — | — | — |
| Bond angles (°) | 0.455 | 0.522 | — | — | — | — |
| Ramachandran plot (%) | | | | | | |
| Favored | 95.01 | 95.32 | — | — | — | — |
| Allowed | 4.99 | 4.68 | — | — | — | — |
| Disallowed | 0.00 | 0.00 | — | — | — | — |
| MolProbity score | 1.47 | 1.58 | — | — | — | — |
| Clashscore | 3.33 | 4.86 | — | — | — | — |
| Poor rotamers (%) | 0.00 | 0.00 | — | — | — | — |

*Particles were picked using three separate methods. Initial particle images include duplicates and triplicates.

†Duplicate and triplicate particle images were removed before final 2D classification.

‡Maps were locally sharpened using DeepEMhancer.

**Table S5.** Substrate dissociation constants for $POLA1_{cat}$.

| | $K_d$* (M) |
|---|---|
| $DNA^9/DNA^{30\ \dagger,\ddagger}$ | $(3.0 \pm 0.3) \times 10^{-7}$ |
| $RNA^9/DNA^{30\ \ddagger}$ | $(3.3 \pm 0.2) \times 10^{-8}$ |
| $RNA^9$-$DNA^{11}/DNA^{41\ \ddagger}$ | $(2.6 \pm 0.7) \times 10^{-8}$ |
| $RNA^9$-$DNA^{20}/DNA^{50\ \ddagger}$ | $(9.4 \pm 0.5) \times 10^{-8}$ |

Data are presented as the mean ± SD (n=3 biologically independent experiments). Source data are presented as a Source Data file.

*$K_d$ is the equilibrium dissociation constant determined from reaction mixtures containing 10 nM oligonucleotide duplex, 20 µM dGTP, and 0.625–5,120 nM $POLA1_{cat}$.

†The DNA primer lacks the 5′-triphosphate group present on the RNA and RNA-DNA primers.

‡Primer and template lengths are indicated with superscripts. Templates were varied to maintain consistent overhangs with all primers.

**Supplemental Table S6.** Cryo-EM data collection, processing, refinement, and validation statistics for the DNA termination dataset.

| | DT subcomplex, partial (EMD-42140) (PDB 8UCU) | DT subcomplex 1, complete (EMD-42141) (PDB 8UCV) | DT subcomplex 2, complete (EMD-42142) (PDB 8UCW) | TC subcomplex | AI complex |
|---|---|---|---|---|---|
| **Data collection and processing** | | | | | |
| Magnification (×) | 105,000 | 105,000 | 105,000 | 105,000 | 105,000 |
| Voltage (kV) | 300 | 300 | 300 | 300 | 300 |
| Electron exposure (e−/Å$^2$) | 55.8 | 55.8 | 55.8 | 55.8 | 55.8 |
| Defocus range (μm) | −0.8 – −2.0 | −0.8 – −2.0 | −0.8 – −2.0 | −0.8 – −2.0 | −0.8 – −2.0 |
| Pixel size (Å) | 0.82 | 0.82 | 0.82 | 0.82 | 0.82 |
| Symmetry imposed | C1 | C1 | C1 | C1 | C1 |
| Initial particle images (no.)* | 25,216,892 | 25,216,892 | 25,216,892 | 25,216,892 | 25,216,892 |
| Particle images after 2D classification (no.)† | 1,570,160 | 1,570,160 | 1,570,160 | 1,570,160 | 1,570,160 |
| Final particle images (no.) | 278,995 | 74,899 | 62,414 | 296,492 | 994,673 |
| Map resolution (Å) | 2.85 | 3.81 | 3.64 | 8.16 | 2.71 |
| FSC threshold | 0.143 | 0.143 | 0.143 | 0.143 | 0.143 |
| Map resolution range (Å) | 2.8 – 4.0 | 3.3 – 9.5 | 3.2 – 8.8 | — | — |
| **Structure refinement and validation** | | | | | |
| Initial model used (PDB code) | 8G9N | 8G9O, 8UCU | 8G9O, 8UCU | — | — |
| Model resolution (Å) | 3.04 | 3.86 | 3.75 | — | — |
| FSC threshold | 0.5 | 0.5 | 0.5 | — | — |
| Map sharpening *B*-factor (Å$^2$)‡ | — | — | — | — | — |
| Model composition (no.) | | | | | |
| Non-hydrogen atoms | 7,468 | 9,850 | 8,264 | — | — |
| Protein residues | 868 | 1,054 | 868 | — | — |
| RNA/DNA nucleotides | 25 | 66 | 63 | — | — |
| Cofactors/ions | 1 | 3 | 2 | — | — |
| Avg. *B*-factors (Å$^2$) | | | | | |
| Protein | 65.1 | 133.1 | 118.5 | — | — |
| RNA/DNA | 50.2 | 113.0 | 129.5 | — | — |
| Cofactor/ion | 79.3 | 194.7 | 177.3 | — | — |
| R.m.s. deviations | | | | | |
| Bond lengths (Å) | 0.003 | 0.003 | 0.002 | — | — |
| Bond angles (°) | 0.444 | 0.534 | 0.475 | — | — |
| Ramachandran plot (%) | | | | | |
| Favored | 96.98 | 96.85 | 96.98 | — | — |
| Allowed | 3.02 | 3.15 | 3.02 | — | — |
| Disallowed | 0.00 | 0.00 | 0.00 | — | — |
| MolProbity score | 1.25 | 1.43 | 1.31 | — | — |
| Clashscore | 2.84 | 4.61 | 3.44 | — | — |
| Poor rotamers (%) | 0.00 | 0.00 | 0.00 | — | — |

*Particles were picked using three separate methods. Initial particle images include duplicates and triplicates.

†Duplicate and triplicate particle images were removed before final 2D classification.

‡Maps were locally sharpened using DeepEMhancer.
